# Intein-assisted bisection mapping systematically splits proteins for Boolean logic and inducibility engineering

**DOI:** 10.1101/2020.11.30.381921

**Authors:** Trevor Y. H. Ho, Alexander Shao, Zeyu Lu, Harri Savilahti, Filippo Menolascina, Lei Wang, Neil Dalchau, Baojun Wang

## Abstract

Split inteins are powerful tools for seamless ligation of synthetic split proteins. Yet, their use remains limited because the already intricate split site identification problem is often complicated by the requirement of extein junction sequences. To address this, we augmented a mini-Mu transposon-based screening approach and devised the intein-assisted bisection mapping (IBM) method. IBM robustly revealed clusters of split sites on five proteins, converting them into AND or NAND logic gates. We further showed that the use of inteins expands functional sequence space for splitting a protein. We also demonstrated the utility of our approach over rational inference of split sites from secondary structure alignment of homologous proteins. Furthermore, the intein inserted at an identified site could be engineered by the transposon again to become partially chemically inducible, and to some extent enabled post-translational tuning on host protein function. Our work offers a generalizable and systematic route towards creating split protein-intein fusions and conditional inteins for protein activity control.

## Introduction

Synthetic split proteins are useful tools for biologists^1^. Often, they are created to serve as sensors for protein-protein interaction^2^, detectors for biomolecules^3, 4^, molecular switches^5, 6^, and logic gates in synthetic circuits^7–10^. In other instances, proteins are split to be endowed with temporal-spatial^11^ or user-defined controls^12^, or to reduce their sizes for viral delivery^13^. Generation of split proteins invariably demands the resulting bipartite sections remain individually inactive, and that they can reconstitute spontaneously or with external assistance, followed by the restoration of protein function^1^.

A specific niche of split protein engineering concerns the use of an intein as the driver for split parts reconstitution. Inteins are <u>in</u>ternal pro<u>tein</u> elements that are expressed as part of a larger precursor protein^14–16^. Upon proper folding, an intein excises itself from the precursor protein and ligates the flanking <u>ex</u>ternal prot<u>eins</u> (exteins) with a peptide bond, producing a product as if the intein was absent from the original gene sequence. An intein split at an appropriate position generates a bipartite intein that can spontaneously self-assemble and undergo the splicing process. Split inteins thus enable reconstitution of separate coding sequences with minimal scarring, reducing the chances of additionally inserted residues or domains that might compromise the original functionality upon protein reconstitution. Documented uses of protein split by inteins include in vivo DNA sensors^17^,protein-based logic gates for bio-computation^18, 19^ and enforcing dual conditions in directed evolution^20^.

One of the major challenges in splitting proteins is the identification of split sites that ensure function loss in split parts but permit protein reconstitution. The use of inteins introduces an extra layer of complexity – inteins require specific extein junction sequences for efficient splicing^21, 22^. Hence, the composition of amino acid residues around a chosen split site needs to be carefully considered. Alternatively, one can insert characterized extein junctions at a putative split site, or modify host protein residues around a putative split site to satisfy extein junction requirements. Doing so could risk perturbations to the protein structure and function. By either approach, the split site design space is often sparsely sampled by educated guesses with split sites tested through trial-and-error^6, 23, 24^, which is inefficient. A better solution would be to predict split sites computationally from protein crystal structures. A method was recently developed to predict intein-insertable split sites by searching flexible regions on protein structures and regions that lack functional conservation^25^. This was demonstrated on antibiotic resistance genes. Another method, SPELL^26^, takes protein structures, calculates split energies and identifies surface-exposed loops that contains low conservation in sequences to predict split sites. SPELL was designed to split proteins with a pair of chemically inducible dimerization (CID) domain, but the principle behind might be general enough for use with inteins. While these computational methods provided better rationality in testing split sites, they require 3D structures. If crystal structures are unavailable homology modelling could be used to generate predicted protein structures, but using predicted structures as inputs might reduce prediction accuracy.

To facilitate split site identification without knowing protein structures, we customized and improved previously described mini-Mu transposon-based approaches^27–31^. We developed a bisection mapping method that involves the fusion of a pair of split intein to the bisected host protein parts. The technique was applied to four proteins and revealed novel split sites for achieving the AND and NAND logic. We highlighted the advantage of using an intein compared to interacting domains in splitting proteins, employed our method to evaluate a single case of split site prediction from protein structural homology, and described suppressing uninduced activities by splitting highly active proteins. Finally, we demonstrated in principle that, once functional split sites for an intein are identified, some degree of post-translational inducibility can be engineered into the intein to achieve drug-dependent control of protein function.

## Results

### Designing the IBM workflow for split site screening

In pursuit of a systematic protocol to search for split sites for inteins, we built upon existing methods utilizing the mini-Mu transposon, bisection mapping^27^ (BM) and domain-insertion profiling^28, 29^ (DIP), and incorporated features from the latter into the former. In brief, the method (**Fig. 1a**, **Supplementary Fig. 1**) starts with an in vitro transposition reaction that randomly inserts a BbsI and SapI-flanked transposon into a staging vector, which hosts a slightly trimmed, BsaI-flanked coding DNA sequence (CDS) of interest (**Supplementary Fig. 2**). This is followed by size selection of the insertion library such that only CDS fragments with insertions would be isolated and ligated into a vector for protein expression. A Golden Gate reaction is then used to irreversibly substitute the transposon with a DNA fragment. The fragment carries a selection marker, a split intein, and transcription and translation initiation elements for carboxyl-lobe (C-lobe) expression. In-frame insertions in the right orientation would thus split a CDS into two with the amino-lobes (N-lobes) and C-lobes of the split intein as fusion partners, under separate control of two inducible promoters. The final library is then screened for individual clones that display functional activities only when the chemical inducers for both promoters are present. The clones are then sequenced at the fusion joints to reveal the split sites.

**Fig. 1.**
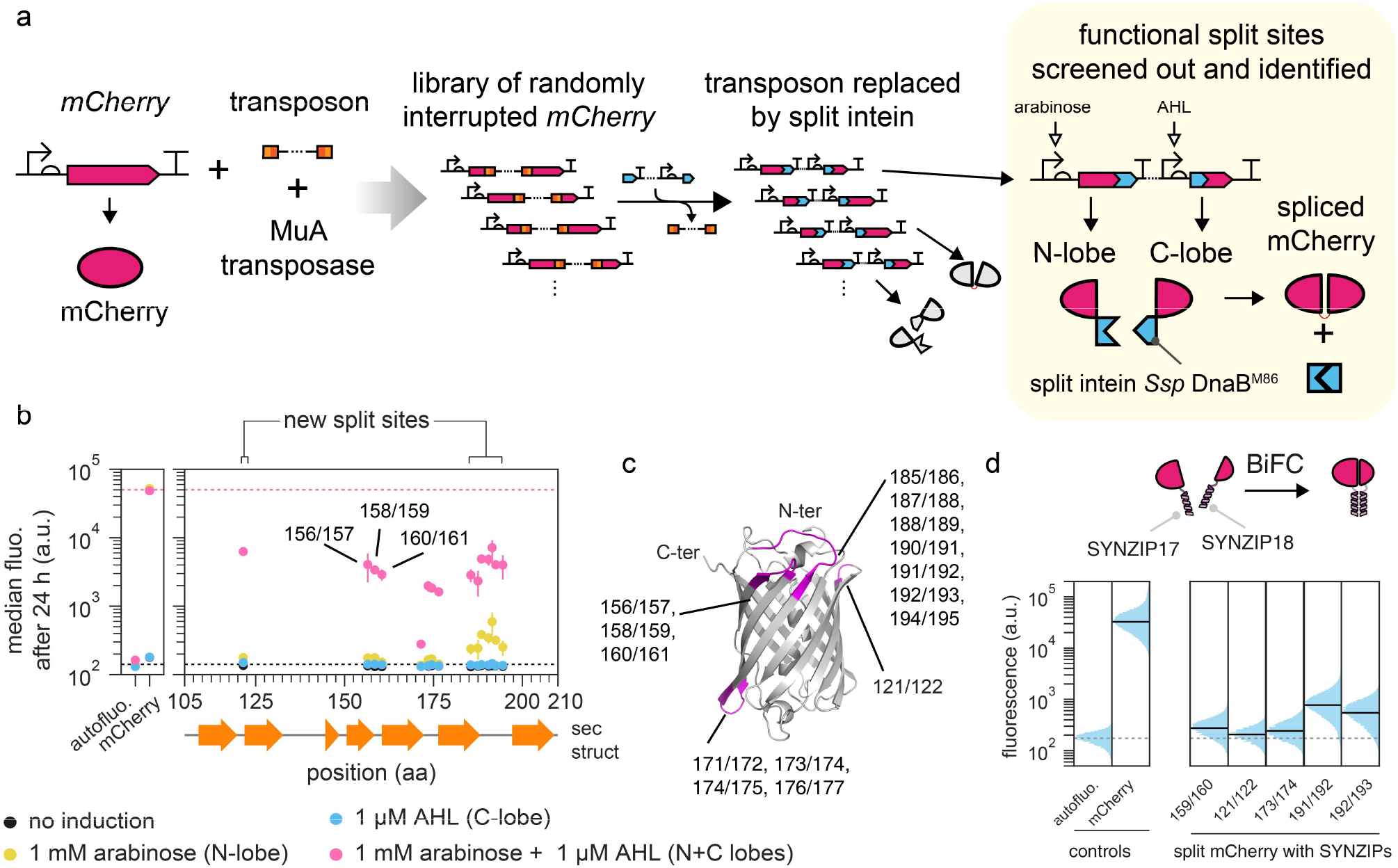
Proof-of-concept of Intein-assisted Bisection Mapping (IBM) on mCherry to recover known split sites for BiFC and discover new ones. **a.** Workflow. A transposon was randomly inserted into the mCherry CDS, and was then substituted with a DNA fragment containing the split intein Ssp DnaB^M86^. This generated a library of mCherry-split intein fusions that can be screened for fluorescence only when both the N and C-lobes were expressed. **b.** Two new loops for split sites were identified within mCherry and two existing ones were recovered. Each vertical group of spots represents an identified split site, aligned to the mCherry secondary structure below. The majority of sites are between the β-sheets of the barrel. y locations and error bars are mean and std of median fluorescence from experiments performed on 3 different days. Horizontal dashed lines bound the range of fluorescence that split mCherry could yield. See **Supplementary Fig. 5** for site distributions and activities at 5 h. **c.** Split sites mapped to a reconstructed mCherry 3D structure (PDB: 2H5Q). Each split site has the −1 and the +1 amino acid residues colored. **d.** Representative split sites from each loop on mCherry permitted biomolecular fluorescence complementation (BiFC). Single-cell fluorescence values were pooled from 3 biological replicates. Solid black horizontal lines denote population median, except for autofluorescence which was denoted by dotted grey lines.

For the split intein, we selected the *Ssp* DnaB^M86^ intein^32^ (thereafter referred as the M86 intein) since it only requires the −1 and +1 extein residues for splicing. These extein residues are incorporated into the substitution insert. Successful splicing of the M86 intein would leave behind a highly predictable four-residue peptide linker at the split site of the original protein (**Supplementary Fig. 3**). Our method is an augmentation of BM by an intein - hence the name intein-assisted bisection mapping (IBM).

### mCherry as a proof of concept for the IBM workflow

To preempt potential difficulties in troubleshooting if the IBM workflow returned no functional split sites, we first carried out a proof of principle test utilizing mCherry as the target protein to be split. mCherry has been employed as a reporter for Bimolecular Fluorescent Complementation (BiFC) and two split sites, 159/160 and 174/175, were known to create bipartite lobes that would regain functionality if brought into proximity^33^. We created a split mCherry construct that simulated the known split site 159/160 being sampled by IBM. This construct only gave increased fluorescence when the inputs for N-(inducible by arabinose) and C-lobes (inducible by AHL) were present (**Supplementary Fig. 4**). Thus, provided enough library coverage, a successful execution of the IBM workflow should generate the simulated control as a member within the final library, and the control should be recoverable afterwards.

Given the assurance, we proceeded to run the IBM workflow on the mCherry protein (**Fig. 1a**). The resulting library was induced with both arabinose and AHL. Cells with fluorescence above autofluorescence were sorted by fluorescence-activated cell sorting (FACS), plated and isolated as single colonies. Individual strains were then assayed for responses in the absence or presence of the two inducers, and those that showed AND logic behavior were subsequently sequenced to identify the split sites. Pooling the results yielded an intein-bisection map (**Fig. 1b**, **Supplementary Fig. 5**). A total of 15 split sites were identified. At all sites, protein splicing was proven by a Western blot (**Supplementary Fig. 6**). All split sites clustered into 4 seams, which were mostly located on loops between the β sheets of the barrel. The second and the third seams should cover sites 159/160 and 174/175, though curiously, site 159/160 was not sampled by this IBM attempt. This site was present in the library we screened (**Supplementary Table 1**) and was left out fortuitously, likely due to under sampling.

To our knowledge, the seams 121/122 and 185/186 - 194/195 were never described to contain functional split sites before. Of equal interest was the fact that four split sites, while close to the loops, were found between amino acids that constituted the β sheets (**Fig. 1c**), and this could suggest tolerance of either structural disruption or inserted linkers, both of which are unlikely to be ever attempted if split sites were designed rationally. Together, these two observations showcased that IBM has the potential to discover novel and unexpected split sites.

We also noticed that sites within seam 185/186 - 194/195 yielded a low level of fluorescence when the cells were grown under prolonged induction (24 h) of the N-lobe alone. Other bipartite mCherry split at other seams did not show such behavior. This might be explained by a leaky P/_ux2_ promoter, and the fact that mCherry split at the last seam would produce relatively shorter C-lobes that were easier to transcribe and translate.

Since the literature reported split sites on mCherry were developed for bimolecular fluorescent complementation^33^, we asked whether the new split sites identified could serve the same purpose. We arbitrarily selected one or two representative split sites from each seam. We also included three additional sites that were found within β-sheets. For each site, we built split mCherry constructs where the split M86 intein was removed or replaced by a pair of synthetic and heterodimerizing coil-coiled domains, SYNZIP17 and SYNZIP18^34^ (**Fig. 1d**, **Supplementary Fig. 7**). For all constructs where split sites were on flexible loops, increase in fluorescence could be observed when both lobes were expressed with SYNZIPs, demonstrating that for mCherry, tolerances of the IBM-identified split sites to protein fusion were not unique to the intein. Sites within β-sheets, except 176/177, did not yield an increase in fluorescence, likely due to structural disruptions to the β-barrel. To test whether the N- and C-lobes could complement without external help from SYNZIPs, we removed SYNZIP17 from the N-lobes and conducted the experiment. Results showed no increase in fluorescence and Western blots proved that it was not due to lack of protein expression.

### IBM identified a computationally unpredicted split site on β-lactamase

Since a computational method^25^ exists to predict split sites for the gp41-1 intein^35^ on antibiotic resistance genes, we asked how a library approach through IBM would perform in comparison to a computational approach. We focused on the case of TEM-1 β-lactamase (also abbreviated as BLA in figures). It has a resolved crystal structure and a well-established split site at 194/196^36^ or 195/196^37^. The previously mentioned computational method identified a new split site at 104/105, which when used with the split gp41-1 intein, allowed co-selection of two plasmids with just ampicillin^25^. We therefore sought to recover these two sites with IBM. We employed the gp41-1 intein with the −2, −1 and +1, +2 minimal extein residues (GY/SS)^38^ and passed β-lactamase through the IBM pipeline. Split clones that conferred ampicillin resistance were enriched through selection and outgrowth. Subsequent sequencing on candidate clones revealed 6 split sites on 2 split seams (**Supplementary Fig. 8**). The first seam, consisting sites 192/193 and 196/197 corresponded to the established site of 195/196. Whereas the second seam (260/261, 261/262, 264/265, 267/268) represented a previously unreported split location. We noted that the absence of the computationally predicted site 104/105 could be due to the clones being outcompeted during the enrichment process. Hence, IBM was not necessarily superior to computational methods, but could complement them in identifying more intein-insertable split sites.

### Applying IBM to engineer AND and NAND logic gates

Having established the IBM workflow, we then sought to demonstrate its universality in engineering protein-based logic gates^10, 39, 40^. We focused on transcription factors because their responses could be directly converted to an assayable fluorescent output (**Fig. 2a**). We chose the repressor TetR and its homolog SrpR from the same protein family^41^, and an activator, the extra cytoplasmic sigma factor 20 (ECF20)^42^. Each protein was fed into the IBM workflow using the M86 intein and a corresponding intein-bisection map was generated. 3 split seams and 32 split sites were found for TetR; 4 seams and 13 sites for SrpR; 3 seams and 17 sites for ECF20 (**Fig. 2b-d**, **Supplementary Fig. 10-12**). Most of the split sites for TetR clustered around loop regions between helices from the TetR crystal structure (PDB: 4AC0, **Supplementary Fig. 13**). It is noteworthy that the same was observed for SrpR and ECF20 even though their shown secondary structures were only predictions that we generated from JPred4^43^. The performance of the logic gates in aspect of on and off states strongly depends on the split protein, the split site locations as well as the time elapsed since induction. Across most split sites found in TetR and ECF20, the split proteins would show qualitative NAND and AND gate behavior with good repression and activation strengths at both 5 and 24 h. For SrpR, most split sites yielded NAND gate behavior at 5 h post-induction with observable levels of repression, but at 24 h, expression of C-lobes alone sufficed to evoke a strong repression rendering the circuit more like a single input responsive gate (**Supplementary Fig. 11**). This was caused by an accumulation of the N-lobes from leaky *ParaBAD* expression over time and the NAND behavior could be restored by eliminating the leakiness (**Supplementary Fig. 13**), proving that either N- or C-lobe alone was capable of repression. Our results thus demonstrated that the IBM workflow is generally applicable and could return multiple sites for choosing the most appropriate gates for user-defined applications.

**Fig. 2.**
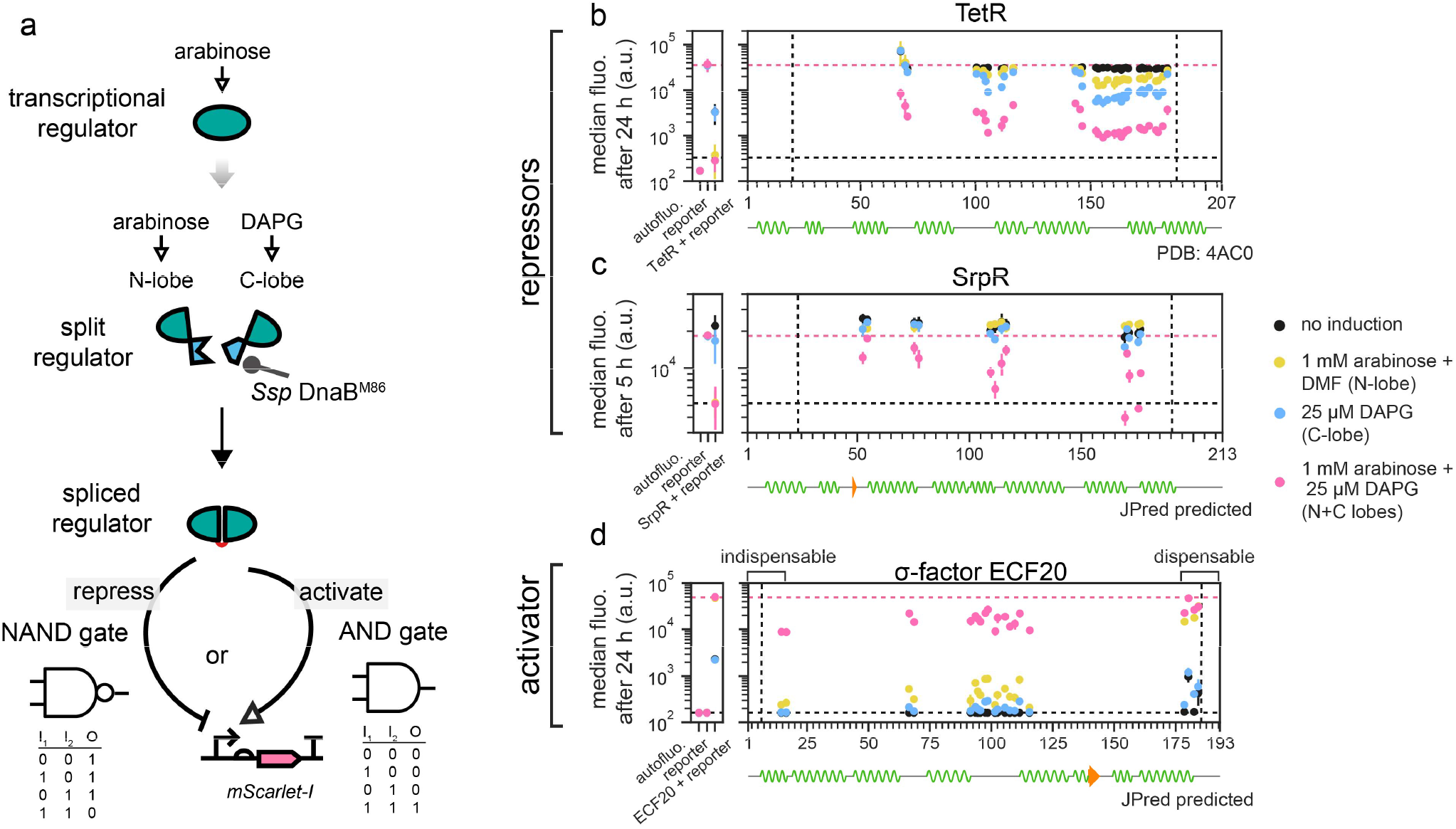
IBM as a universal method to exhaust split sites for AND and NAND logic gate engineering. **a.** Any transcription factor with a function that can be wired to an assay-friendly output could be subjected to IBM for logic gate engineering. **b-d.** Intein-bisection maps for TetR (**b**, 3 seams identified), SrpR (**c**, 4 seams), and ECF20 (**d**, 3 seams). Split clones of TetR and SrpR (or ECF) achieved major off (or on) activities when both the N- and C-lobes were present. y locations and error bars are mean and std of median fluorescence from experiments performed on 3 different days. Vertical dashed lines bound the permitted transposition window and horizontal dashed lines bound the ranges of activities that could be attained by the split proteins. **d.** By-products of IBM revealed a truncatable region of ECF20 which when removed did not adversely affect activation. **b-d.** See **Supplementary Fig. 6-8** for site distributions and activities at 5 h (TetR and ECF20) and 24 h (SrpR). See **Supplementary Fig. 9** for explanations of controls (leftmost subplots).

### IBM indirectly defined functional boundaries on the ECF20 activator protein

While screening the colonies for AND gates in ECF20, we observed that 80% yielded strong activation activities from the expression of N-lobes alone (data not shown), which emulated the responses of an intact protein and addition of C-lobes did not further improve activities. We thus surmised that they could be truncations at the C-termini and sequenced some of them. Indeed, those split sites were clustered at 178185 (**Fig. 2d**) and approximately corresponded to the end of the last helix on ECF20, suggesting that the residues beyond position 178 could be trimmed without loss of function. In contrast, the first helix was crucial since the first AND gate split site was found immediately after it. These two observations suggested that IBM could be repurposed to determine the minimal functional size of a protein.

### IBM expands the range of split sites discoverable in TetR

Previously reported approaches in bisection mapping for logic gates engineering utilized protein-protein interacting domains like the SYNZIPs as fusion partners^30^. We hypothesized that inteins make a better choice because they would be excised from the splice product, whereas additional domains could exert steric hindrance, especially when the host protein function requires multimerization or interactions with other proteins. To test our hypothesis, a representative site from each split seam identified on split TetR was selected and the split M86 intein was replaced by SYNZIPs in a similar manner as above (**Fig. 3a**). Of the three tested sites, site 166/167 showed a stronger level of repression (~ 4 fold) compared to the other two (~ 2 fold), despite having the least possible amount of reconstituted protein (**Supplementary Fig. 15**). The differences in repression strengths between split sites, when SYNZIPs were used, demonstrated their differential tolerances towards additional domains. Whereas in IBM, all three sites showed good levels of repressions that were strong enough to be identified from a single screen. Hence, the use of an intein could enable split sites that might be inaccessible by protein-protein interacting domains and expands the range of split sites that could be identified for logic gate engineering.

**Fig. 3.**
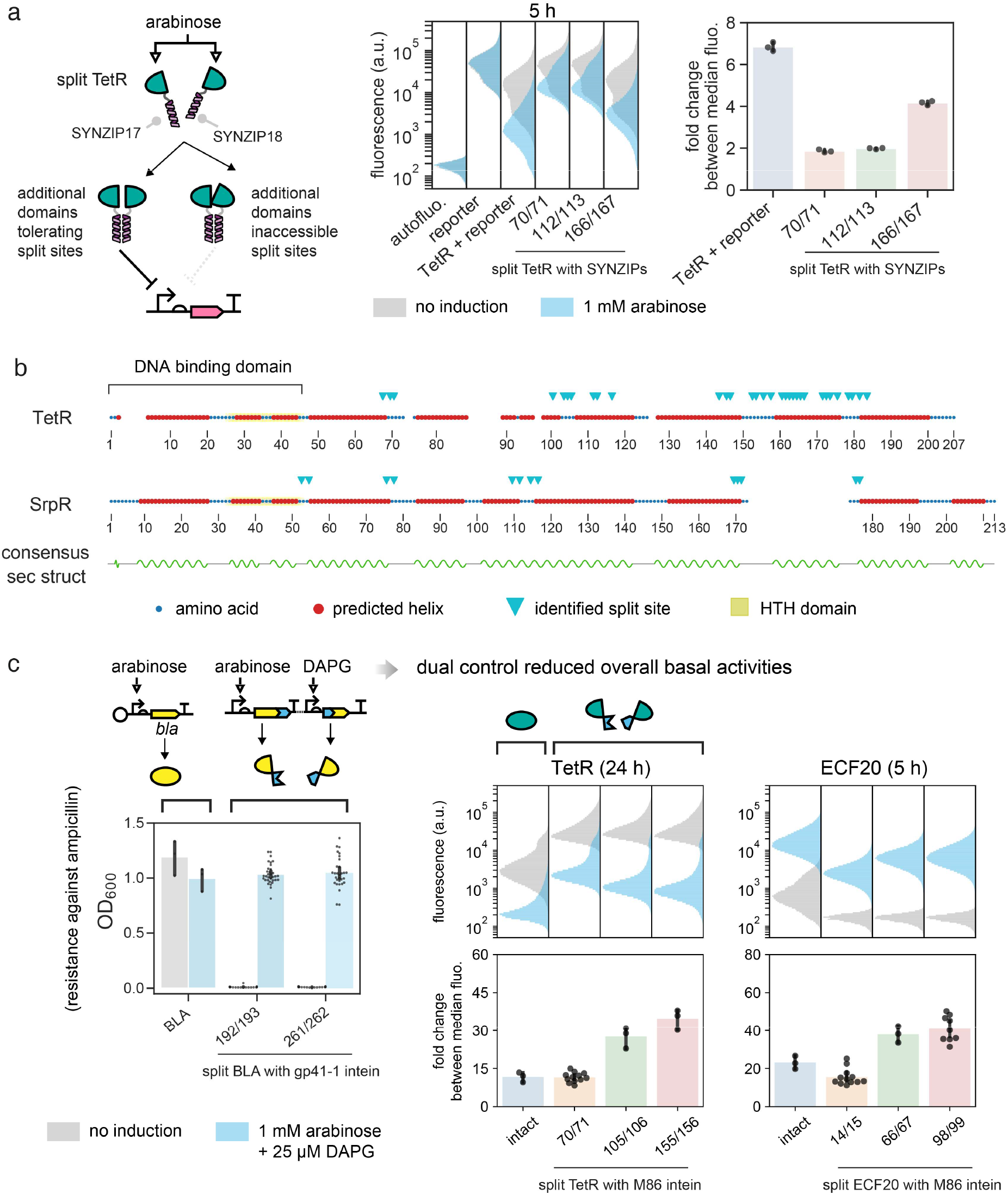
Benefits of IBM and reduction of basal activities by IBM. **a.** Substitution of the split M86 intein inserted in split TetR by SYNZIP. Representative sites were chosen within the 3 identified split seams. Results showed different split sites had differential tolerances towards additional protein domains, but all split sites functioned well when the intein was used (**Fig. 2b**). Single-cell fluorescence values were pooled from 3 biological replicates. **b**. IBM-identified split sites of homologous TetR and SrpR do not necessarily map to loops between consensus helical domains, highlighting the limitation of guessing split sites for a new protein by secondary structure alignment. **c**. Splitting highly active proteins can reduce their basal activities. Left panel: leaky expression of BLA led to ampicillin resistance in absence of induction, which could be improved by splitting BLA. Middle and right panels: Fluorescence distributions and fold changes between fully on and fully off states of intact and split TetR and ECF20 were shown. Representative sites were chosen from each identified seam. In most cases the split clones had lower basal activities and therefore larger fold changes between on and off states. **b-c.** Data reused from Fig. 2 and Supplementary Fig. 8 but further analyzed. **a, c.** Single-cell fluorescence values were pooled from experiments performed on 3 different days. In fold change calculations, bar heights and error bars represent mean and std.

### IBM revealed limitations in inferring split sites from secondary structure alignment

When choosing split sites for a new protein from a known protein family, a conventional approach is to align the sequences and their predicted secondary structures, and find common loops between homologous structures to identify putative split sites^44^. Upon the completion of the intein-bisection maps of TetR and SrpR, we realized our data might shed light on how reliable the approach is, and therefore aligned the two protein sequences by secondary structures^45^ along with the identified split sites (**Fig. 3b**, **Supplementary Fig. 16**). The alignment indicated that only two split seams are shared between TetR and SrpR. Sites on seam 2 of TetR and seam 3 of SrpR were not aligned and would be overlooked if the alignment approach were taken. Likewise, on SrpR split sites were found on the loop that demarcates the DNA binding domain from the dimerization domain, but those sites would be unexpected if TetR served as the reference model. Our results therefore suggest the secondary structure alignment approach works but could miss other potentially useful split sites, which could be discovered by IBM.

### Mitigation of undesirable basal activities in highly active proteins through IBM

Splitting highly active proteins could suppress their background activities. This was best illustrated by the IBM generated split β-lactamases (**Fig. 3c**). When the full CDS of β-lactamase was placed under the control of *P_araBAD_,* the hosting bacteria could grow in ampicillin regardless of arabinose addition, proving that leaky expression of β-lactamase was sufficient to confer resistance. In contrast, split β-lactamases did not lead to cell growth if inducers were absent. This tightening of protein expression control could also be concluded from further analyses of single-cell fluorescence data from **Fig. 2**. Intact TetR and SrpR had stronger repression than their bipartite counterparts at 5 h. At 24 h, however, in the absence of induction the fluorescence of unrepressed cells was much lower (**Fig. 3c, Supplementary Fig. 17**). Whereas, many bipartite repressors at 24 h had higher unrepressed fluorescence, better separation of populations between on and off states and hence high fold changes. This phenomenon was even more pronounced on the ECF20 activator - at 5 h, basal activities already gave a strong off-state fluorescence and four split constructs started to benefit from fold change improvement, and at 24 h the average fold change of the worst performing split ECF20 construct was around 54 compared to 22 of the intact ECF20 (**Supplementary Fig. 17**). These data from β-lactamase, repressors and activators implied that intact proteins had accumulated over time due to leaky expression from a single promoter, but when they were split by IBM and placed under independent promoters, the conferred AND logic led to a lower probability of assembling a functional protein, thereby reducing the overall undesirable basal activities at the off states.

### Transposon-mediated engineering of post-translational inducibility into inteins to control protein function

In theory, our IBM workflow should allow one to use a conditional intein^46–48^ in place of a split intein, and then screen for insertion or split sites that would enable post-translational control of protein function. We thus synthesized and tested three reported chemogenic conditional inteins^49–51^ from the literature, but they did not work well under our specific context (**Supplementary Fig. 18**). Given the circumstances we decided to take a different approach and asked whether we could create inducible inteins de novo, by transposing drug-controlled domains into the M86 intein. We reasoned that a conversion of the spontaneous split intein into a conditional one would allow us to exert control over protein activity through addition or removal of small molecules (**Fig. 4a**). To do so, we synthesized the ligand binding domain of human estrogen receptor (ER-LBD)^29^ and the camelid anti-caffeine VHH (acVHH) antibodies^52, 53^. For the former, we hypothesized that similar to the previous study, possible insertion sites exist within the cis-intein, where intein folding would be obstructed by the interrupting ER-LBD until 4-hydroxytamoxifen (4-HT) binding elicits a structural change that relieves the effect. For the latter, we hypothesized that at certain split sites, the split intein lobes would suffer from reduced interacting surfaces and hence diminished affinities for spontaneous assemblies, but could be rescued by an externally supplied source upon ligand-induced dimerization of acVHH.

**Fig. 4.**
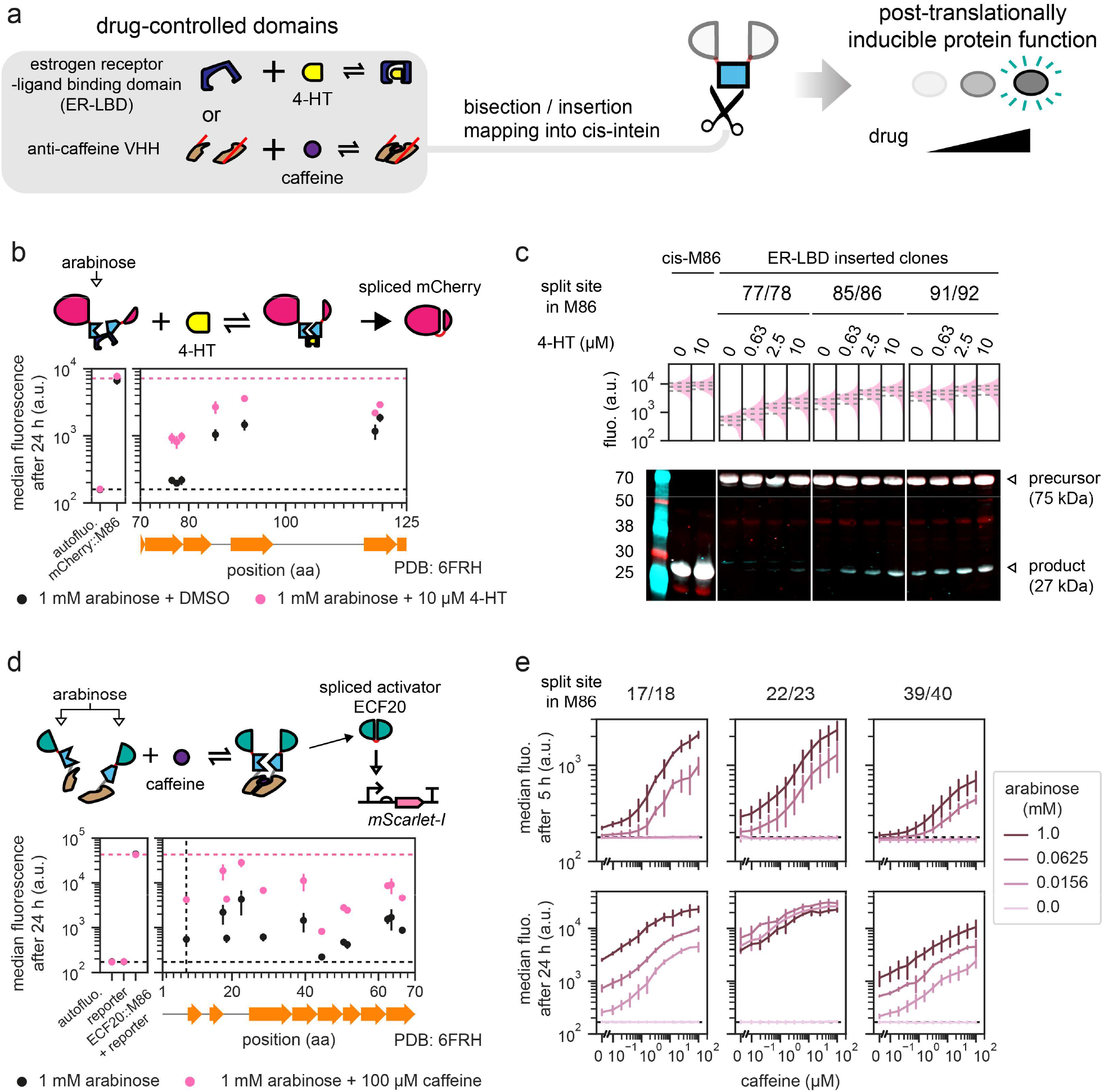
Transposon as a tool to engineer inducibility into an intein for protein function control. **a.** The M86 intein in its cis-form can be inserted in an identified split site within the protein of interest, and then bisected or inserted with drug-controlled domains, which leads to inducible splicing and inducible function of the host protein. **b**. 7 insertion sites identified via domain-insertion mapping of the estrogen receptor-ligand binding domain into the M86 intein within mCherry interrupted at 192/193, where addition of 4-HT led to increased fluorescence. **c**. Selected clones from (b) showed gradual increase in fluorescence and spliced product formation as the concentrations of 4-HT increased, despite the fact that most precursors were unspliced. Single-cell fluorescence values were obtained from one biological sample. Interquartile ranges are denoted by horizontal dashed lines. mCherry N-lobes were labelled in red and C-lobes were labelled in turquoise, and their superpositions give a white color. **d.** 11 split sites for anti-caffeine VHH identified on the M86 intein that interrupted the activator ECF20 at 101/102. In those clones, addition of caffeine led to increased activation activities. **e.** Selected clones from (d) had increased activation activities as caffeine concentration increased, and leakiness due to spontaneous assembly could be mitigated by lowering the total split protein concentrations. **b, d.** Each vertical group of spots represents an identified insertion/split site, aligned to the M86 intein secondary structure. See **Supplementary Fig. 19, 22** for site distributions and activities at 5 h. **b, d, e.** y locations and error bars are mean and std of median fluorescence from experiments performed on 3 different days. Horizontal dashed lines bound the maximum (b and d only) and minimum fluorescence that could be achieved by the split or inserted constructs.

### Engineering a 4-HT-inducible mCherry by domain-insertion mapping

In our first example we aimed to control splicing and fluorescence of split mCherry using a slightly modified version of domain-insertion profiling. We first inserted the cis-M86 intein between amino residues 192 and 193 of mCherry since this mCherry split site performed well in the fluorescence complementation experiment (**Fig. 1d**) and was at the center of the flexible loop. We then performed the domain-insertion of the ER-LBD into the cis-M86 intein, now between mCherry(1-192) and mCherry(193-236), via random transposition, and then replaced the min Mu-transposon by the CDS of the ER-LBD domain (**Fig. 4b**). We induced protein expression by the addition of arabinose, performed a series of positive and negative cell sorting with or without 4-HT, obtained individual strains, characterized and then sequenced them. Of the 7 insertion sites identified, none yielded any activities at 5 h post-induction (**Supplementary Fig. 19**), but differential responses between 4-HT-uninduced and induced states could be observed at 24 h. Some recovered sufficiently high fluorescence comparable to the intact M86 intein but not without sacrificing basal activities.

We then selected three representative sites that spanned the activity ranges and proceeded to verify dosedependent activities. Cells were subjected to a gradient of 4-HT induction and harvested for fluorescence and protein content analysis (**Fig. 4c**). All three strains showed gradual upshift of fluorescent populations as 4-HT concentration increased. Similarly, a Western blot of whole-cell lysates also indicated a 4-HT-dependent increase in spliced mCherry formation, even though most precursors did not undergo splicing regardless of split sites or inducer concentrations. The exception is the strain where ER-LBD was inserted at site 77/78 within the M86 intein, in which the bands of the spliced products were too weak to be detected, and the band for the spliced products under 10 μM 4-HT was barely visible. The Western blot also confirmed the existence of spliced mCherry in the absence of 4-HT and explained the strong basal fluorescence of the inserted clones.

We further asked whether the novel conditional inteins are transferrable to another host protein, and so cloned all seven ER-LBD-inserted inteins into the identified site 101/102 of ECF20. Clones were subjected to either 0 or 10 μM 4-HT induction under various arabinose concentrations (**Supplementary Fig. 20**). In the best-case scenario, strain 76/77 under 0.0625 mM arabinose yielded a small but statistically significant increase in fluorescence when 4-HT was added. Nevertheless, spontaneous splicing already generated sufficient amount of spliced ECF20, which were high active and resulted in strong basal activities, and rendered the circuit impractical for further use.

### Engineering caffeine-inducible activation by acVHH-assisted bisection mapping

In the second example of post-translational inducibility engineering we generated a caffeine-inducible activation by transposing two acVHH domains, each with a 10-residue linker, into the cis-M86 intein at site 101/102 of ECF20. Consequently, the intein was split and the resulting split protein parts were then placed under the control of two arabinose inducible promoters, one for the N-lobe and another for the C-lobe, simplifying the effort to tune protein expression strengths. Prior to the bisection mapping experiment, we established comparable performance between caffeine-inducible dimerization and the rapamycin-inducible FRB/FKPB dimerization in reconstituting an evolved split T7 RNA polymerase^54^ (**Supplementary Fig. 21**). FRB/FKBP had lower propensity to self-associate without ligand but we chose acVHH since caffeine is inexpensive.

We then carried out the acVHH-assisted bisection mapping on the M86 intein, following a similar workflow as we did for the domain-insertion mapping of ER-LBD. 12 split sites close to the M86 N-terminus were identified and they had increased fluorescence when cells were grown under 100 μM caffeine for 24 h when compared to the lack of caffeine induction (**Fig. 4d**, **Supplementary Fig. 22**). Again, all strains had either strong basal and strong maximum activities or weak activities for both states. Among them, three strains displayed discernable differences in fluorescence at 5 h and thus were subjected to tests for dosedependent activation.

From previous literature and the experiment on split T7 RNA polymerase we were cognizant that spontaneous assembly positively correlated with intracellular protein concentrations and could be mitigated by tuning down protein expression. We therefore grew the three strains in a 2-dimension gradient of caffeine and arabinose, and characterized their fluorescence at 5 and 24 h (**Fig. 4e**, **Supplementary Fig. 23**). Activation was an increasing function of caffeine and to some extent the response resembled a hill-curve. Apart from site 22/23 at 24 h post-induction, reducing arabinose concentration could reduce leaky activation at the expense of the overall activities. We then attempted to test those inducible inteins’ transferability, by cloning them back into mCherry split at 192/193 for fluorescence and Western blot assays, but neither showed any signs of splicing (**Supplementary Fig. 24**). We speculated the splicing efficiencies under caffeine induction were too low and only worked with ECF20 given its extreme potency in activation.

The examples of transposing ER-LBD and acVHH into a spontaneously splicing intein generated prototypes of inducible inteins with admittedly limited performance and transferability, but they were only the first steps. With these two examples we proved in principle, there exists a systematic approach to introduce inducibility into an intein, and hence into a protein of interest, provided that its function could be screened conveniently.

## Discussion

We have established IBM as a useful tool for split protein-intein engineering. Thus far we have only employed the Ssp DnaB^M86^ and the gp41-1 intein. Repeating our IBM experiments using different split inteins may reveal even more split sites at new positions, since extein junctions with different amino acid compositions and lengths would be incorporated. This may alter the rigidity of the resulting linkers and hence functionalize other unprobed split sites. If shorter linkers are desired, a “promiscuous” intein^55^ could be helpful, since a designated +2 extein residue could be omitted and supplied by the host protein.

The computational method by Palanisamy et al^25^, and the SPELL algorithm^26^, both mentioned above, suggest split sites based on protein 3D structures and works in silico, whereas our IBM/acVHH-assisted BM workflow takes on an empirical approach and addresses the same issue with different constraints. Sometimes solved 3D structures may be unavailable and de novo structure prediction might not sufficiently reflect the multimerization required in some proteins like TetR. Conversely, IBM/acVHH-assisted BM requires the protein function to be manifested as an easily screen-able output and would be tremendous difficulty or outright impossible if the function of interest is, for example, chemotaxis. It also requires a chassis capable of creating a complex library through highly efficiently transformation, and so for the time being it only applies to bacteria or yeasts. These limitations are absent in computational methods for split site predictions, however. Furthermore, data from IBM/acVHH-assisted BM may be fed back into and refine those algorithms. We therefore advocate IBM/acVHH-assisted BM not as a competitor, but rather, a complement to the computation approaches.

In this work we created two novel inducible variants of the M86 intein. At the current stage, these two inteins have limited dynamic ranges and strong spontaneous splicing activities, and thus are specific to their protein contexts where they were screened. Therefore, our work was a proof of concept, and only the first step towards a systematic approach in converting spontaneously splicing inteins to conditional ones. So far only the M86 intein was tested as the engineering precursor. Other inteins standardized by our group lately might lead to conditional inteins with better on-off characteristics. There are also other synthetic ligand binding domains, for example, uniRapR^56^ and iFKBP/FRB^57^ that are potential replacements for ER-LBD and acVHH. Given so, the performance of any inducible intein engineered through this method may depend on complex interactions between the intein of choice, the drug-responsive domain, and the insertion/split positions. Each could influence the other two factors. Hence, future efforts in engineering more inducible inteins would likely require a combinatorial exploration of all three factors to identify the optimal structure drug-induced protein splicing.

The limited performance of the two inducible intein from this study could also be improved via directed evolution^23, 32, 49, 58, 59^, which can mutate these inteins such they splicing with better efficiencies and with lower basal activities, and allow them to perform sufficiently well when inserted into other host proteins. We foresee a powerful combination between domain-insertion/acVHH-assisted bisection mapping and directed evolution, where the former identifies the optimal sites for differential activities. The sites can then be exploited by the latter which often requires differential responses to prime the evolution process. This should create more inducible inteins which would be valuable tools when additional domains on a protein cannot be tolerated (**Supplementary Fig. 25**).

Our IBM workflow employed random transposition to diversify DNA insertion. This method has known issues including sequence bias^60–64^ and inexhaustive space search^28, 29, 64, 65^. Despite so, in our final libraries prior to screening, sequenced by Next Generation Sequencing (NGS), we obtained at least 87% coverage on possible amino acid split / insertion positions (**Supplementary Fig. 26, Supplementary Table 1**), which sufficiently explored the sequence space. If unbiased and full position coverage is desired, the step of random transposition could be replaced by the recently developed Saturated Programmable Insertion Engineering (SPINE)^64^, which hardcodes all insertion possibilities into oligonucleotide pools. Incorporation of SPINE into the IBM workflow would return bisection / insertion maps with even higher confidences.

In summary, we presented a robust method to screen and identify protein split sites for the insertion of a split intein. We have recently characterized a library of orthogonal split inteins^16^. With the help of IBM, a large number of orthogonal AND or NAND gates could be created in a streamlined fashion, expediting the development of biocomputing units. Moreover, we demonstrated in principle that transposing drug-controlled domains could create inducible inteins to control host protein function post-translationally. Together, they constitute an empirical and systematic approach towards split protein and protein function control engineering, and should benefit general biologists who seek to use inteins on split proteins.

## Methods

### Strains, media, and inducers

*Escherichia coli* TOP10 (Invitrogen) was used for routine cloning. For all bisection/insertion mapping, the first insertion libraries were always transformed into the electrocompetent *E. coli* NEB 10-beta (C3020K, NEB). For the rest of the workflow the strain was switched back to TOP10. The only exception was the IBM experiment on mCherry and its outcome simulation experiment, where NEB 10-beta was used in all steps. All strains were grown in the Miller’s Lysogeny Broth (LB, 10 g/L tryptone, 5 g/L yeast extract, 10 g/L sodium chloride) in liquid medium or agar supplemented with the appropriate antibiotics (unless noted otherwise) at the final concentrations of: kanamycin (K4000, Sigma-Aldrich), 50 μg/mL; chloramphenicol (C0378, Sigma-Aldrich), 25 μg/mL; ampicillin (A9518, Sigma-Aldrich), 100 μg/mL; tetracycline (T8032, Sigma-Aldrich), 10 μg/mL, spectinomycin (ab141968, Abcam), 50 μg/mL.

For preparation of stock inducers, powder of L-(+)-Arabinose (A3256, Sigma-Aldrich, 1 M), N-(3- Oxohexanoyl)-L-homoserine lactone (AHL, K3007, Sigma-Aldrich, 25 mM), or caffeine (A10431.22, VWR, 10 mM) was dissolved in water; 2,4-Diacetylphloroglucinol (DAPG, 16345, Cambridge BioScience, 25 mM), in dimethylformamide (D4551, Sigma-Aldrich); Rapamycin (S1039-SEL, Stratech, 10 mM), (Z)-4- hydroxytamoxifen (4-HT, H7904, Sigma-Aldrich, 10 mM), 3,3’,5-triiodo-L-thyronine (T3, HY-A0070A, Cambridge BioScience, 10 mM) in dimethyl sulfoxide (D8418, Sigma-Aldrich), with the stock concentrations denoted in brackets. For inductions involving inducers dissolved in organic solvents, the volumes of inducer were less than or equal to 1% of the final volume.

### DNA assemblies and purification

Synthetic DNA constructs were built using Gibson Assembly, Golden Gate Assembly and conventional subcloning using restriction digestion and ligation, with the method chosen depending on their individual needs. Whenever necessary, synonymous mutations were introduced to remove internal BsaI, BbsI or SapI restriction sites. Standard molecular biology protocols were observed. The ZymoPURE II Plasmid Midiprep Kit (D4200, Zymo) was used for DNA library extractions. For the purification of DNA, the Monarch Nucleic Acid Purification Kits (T1020 and T1030, NEB) were used. All restriction enzymes and ligases were bought from NEB. MuA protein was purified in collaboration with Domus Biotechnologies (Turku, Finland) essentially as described^66^.

### Cell growth for fluorescence assays and OD measurements

Cells were routinely cultured in 96-well plates (655096, Greiner Bio-One) sealed with breathable membranes (Z380059, Sigma-Aldrich), and incubated at 37 °C in plate shakers (AS-03020-00, Allsheng) with 1000 rpm orbital shaking motion. An assay of synthetic constructs began with an inoculation of a single colony from an agar plate into a well with 200 μL of medium, which was then grown for 16-18 h. The next day, 2 μL of the overnight culture was diluted 1:100 into 198 μL of fresh medium with or without inducers and grown for 5 h. The membrane was then removed and 2 μL of the culture was sampled. A new seal was applied, and the plate was returned to the shaker to further grow until the total time of inoculation was 24 h. Afterwards, 2 μL (5 h) or 0.5 μL (24 h) of the culture was sampled. Changes to growth time were noted in individual figures where appropriate. Exception to the above applies to the split mCherry splicing experiment, the mCherry BiFC experiment and the split TetR-SYNZIP experiment, where overnight cultures were diluted 1:100 into 1mL of fresh medium in 96-deepwell plates (E2896-2110, Starlab). For the assays of strains identified from bisection/insertion mapping experiments, the overnight culture was inoculated from the saved glycerol stocks (described below). Assays measuring resistance against ampicllin were performed in a similar manner, the only difference was that ampicillin was added at the same time as inducers, and growth was only measured after 24 h.

### Optical density measurements by plate reader

End-point optical densities at 600 nm (OD600) were measured with a FLUOstar Omega plate reader (BMG Labtech). The software Omega Control v5.11 (BMG Labtech) was used for data acquisition and Omega MARS v3.32 (BMG Labtech) was used for data export. The optical densities of blank wells from the same plate were subtracted from all other wells.

### Fluorescence measurement by flow cytometry

Prior to analysis, sampled cultures were diluted into 1× phosphate-buffered saline (K813-500ML, VWR) with 2 mg/mL kanamycin to a total volume of 200 μL. Diluted cells sampled at 5 h were incubated at 4 °C for a minimum of one hour to promote fluorophore maturation, whereas those at 24 h were directly assayed. Cells were passed into the Attune NxT Flow Cytometer (Thermo Fisher) equipped with the Autosampler for analysis. For each well, 100 μL of diluted cells were run at 500 μL/min and at least 10^5^ events were recorded. Red fluorescence was acquired on the YL2-H channel (excitation 561 nm, emission 615/25 nm). Exported FCS files were processed using an in-house Python script dependent on the FlowCytometryTools package v0.5.0. All samples were gated on FCS-H, FCS-A, SSC-H and SSC-A for events between 10^3^ - 10^5^ arbitrary units, followed by gating on YL2-A and YL2-H between 1 to 10^6^ arbitrary units.

### Transposition and bisection/insertion library preparations

The mini-Mu transposon used in this study was modified from the one used by Segall-Shaprio et al.^27^ with BbsI and SapI sites incorporated into the R1 recognition sites. Prior to transposition the transposon was released from its host vector by restriction digestion using BglIl followed by purification from agarose gels. The coding DNA sequence of interest was trimmed at the N- and C-termini before being subcloned into a staging vector. In vitro transposition reaction was set up following an established protocol^67^ with slight customization: 150 ng of the staging plasmid and 150 ng transposon were mixed with 660 ng of MuA. For each insertion library 5-6 reactions of 25 μL each were prepared and incubated at 30 °C in a thermocycler for 6 h, followed by heat inactivation at 80 °C for 10 min. All reactions were pooled, purified, and then eluted in 10 μL of nuclease-free water. The resulting DNA was then electroporated into a total of 200 μL of NEB 10-beta cells in four separate cuvettes and recovered following manufacturer’s protocol. Then, 10 μL of the recovered cells (~ 2 mL) were removed, serially diluted into 0.85% sodium chloride (w/v) and spread onto LB agar with kanamycin and chloramphenicol for colony counting. The library coverage was defined as the total number of obtainable transformants / (size of staging plasmid in bp × 2) and were at least 20-fold for all experiments. Libraries that did not meet the criteria were discarded and transposition reactions were repeated. For libraries with sufficient coverage, the rest of the recovered cells were spread onto LB agar. Bacterial lawns were then washed down by 0.85% sodium chloride and a small aliquot was saved as a glycerol stock. The rest were pelleted, and the DNA was extracted by midiprep.

10 μg of the midiprepped DNA from the initial insertion library was digested by BsaI and then resolved on agarose gels until bands were well-separated. The band corresponding to the trimmed coding DNA sequences with insertions was then excised, purified, and ligated to the linearized expression vector in 1:2 molar ratio for insert:vector. The overnight ligated product was then purified and electroporated into 100 μL of in-house-prepared electrocompetent cell, which were recovered in 2 mL SOC for 1 h, concentrated and then spread onto LB agar and grown overnight. Library coverage estimation and DNA extraction of the library were performed similar to that in transposition, except that the size of the insertable positions equals to the size of trimmed coding DNA sequence in bp. At this stage library coverages were typically > 500-fold.

To replace the inserted transposon with split inteins or drug-controlled domains, 60 ng of the midiprepped DNA from the open reading frame (ORF) insertion library was mixed with substitution inserts (released from the cloning plasmids) in a 1:5 molar ratio for plasmid:insert, and added to a Golden Gate reaction mixture^68^ with 20 units of BbsI, 10 units of SapI and 400 units of T4 DNA ligase. The reaction was then run in a thermocycler with the following program: (37 °C for 3 min, 16 °C for 4 min) × 25 cycles, 37 °C for 30 min, and 65 °C for 20 min. Usually 5-6 reactions were run, pooled, purified and electroporated into 100 μL of inhouse-prepared electrocompetent cells. Electrocompetent cells carried a reporter plasmid wherever required. Cells were recovered and the library coverage estimation was performed in the same manner as the preparation of the ORF insertion library. At this stage library coverages were typically > 100-fold.

### Library screening

For IBM on mCherry, recovered cells from the final library were first induced with both arabinose and AHL, and sorted by fluorescence activated cell sorting (see below). Retrieved cells were then spread onto LB agar for colony picking. For IBM on β-lactamase, recovered cells from the final library were first induced with arabinose and DAPG overnight, the culture was then diluted 1:100 in fresh medium containing arabinose, DAPG and ampicillin, and was grown for another overnight. The resulting culture was serially diluted onto solid medium with inducers and ampicillin for isolating single colonies. For IBM on TetR, SrpR and ECF20, recovered cells from the final library were serially diluted such that single colonies could be observed when they were spread onto LB agar with arabinose and DAPG. For TetR and SrpR, functional reconstitution of the bipartite protein represses expression of mScarlet-I and therefore yielded visibly white or pale pink colonies. The opposite was true for ECF20. These colonies were picked directly. For the M86 intein inserted with ER-LBD or the M86 intein bisected by acVHH, recovered cells from the final library were first induced with arabinose and 4-HT or caffeine. They were then sorted for populations with fluorescence higher than the library without 4-HT or caffeine induction (positive sort). Sorted cells were regrown in the presence of arabinose only and then sorted for populations with lower fluorescence (negative sort). The positive sort was repeated once, and the retrieved cells were spread onto LB agar to obtain single colonies for picking.

### Fluorescence activated cell sorting (FACS) experiment

For the initial sort, 100 μL of the library was inoculated into 25 mL of medium with inducers and grown overnight for 18 - 24 h. The next day, 10 μL of the culture was diluted into 10 mL 1× phosphate-buffered saline and then passed into the cytometer. Cell sorting was performed on a FACS Aria II cytometer (BD-Biosciences) through the red fluorescent channel (excitation 561 nm, emission 610/20 nm), under the Purity Mode. Cells were first gated on irregularly shaped FSC-A and SSC-A gates to exclude non-cellular materials, and then gated on boundaries defined by the previous libraries with or without induction. Gate sizes and positions were tailored to individual experiments. Typically, 0.5 - 1 million gated events were collected into a 15 mL conical tube with 5 mL of LB supplemented with 1 % D-Glucose (10117, VWR) and without antibiotics. Collected cells were recovered for 2 h at 37 °C with 160 rpm shaking. Then, the volume was topped to 15 mL using LB with the next set of inducers and grown overnight for 16-18 h for the next sorting experiment. After the final cell sorting, the overnight culture was diluted and plated onto LB agar to obtain single colonies for strain isolation.

### Candidate strain isolation, characterization and split/insertion site mapping

In most cases > 500 single colonies with desirable traits were individually picked into 96-well plates with 200 μL of LB medium for 16-18 h of growth, which were subjected to 16-24 h of induction assays to look for AND logic (mCherry, β-lactamase and ECF20), NAND logic (TetR and SrpR), or differential expression (with the M86 intein inserted or bisected). The fluorescence of the candidate clones was then measured on the FLUOstar Omega plate reader (BMG Labtech) and ranked. The best clones with desirable traits (~96 for IBM and ~40 for M86 intein engineering) were isolated with assistance from an OT-2 robot (Opentrons). The shortlisted clones were saved as temporary glycerol stocks and then subjected to fluorescence assays as described above for proper characterization, which generated the data for plotting the bisection/insertion maps. Strains that showed strong experiment-to-experiment variations in fluorescence were excluded for further use. A small aliquot of each cell strain in liquid suspension was then subjected to polymerase chain reactions (PCR), which amplified the N-terminal (mCherry) or the C-terminal (all others) joints. The PCR products were purified and sent for Sanger sequencing. Poor sequencing results or reads that suggested non-single clones were discarded. The rest of the sequencing results were analyzed with a customized Python script utilizing the Biopython package v1.76^69^ to deduce split or insertion sites by local alignment of sequences. Sites were mapped back to their fluorescence profiles and protein secondary structures (rendered using the Biotite package v0.20.1^70^).

### SDS-PAGE and Western blots for whole-cell lysate analysis

Cells cultured for Western blots were grown in 30 mL universal tubes (E1412-3011, Starlab) placed inside a shaker (Infors HT) maintained at 37 °C, 160 rpm. For each sample, a single colony was inoculated into 2 mL of medium and grown for 16-18 h. The next day, the overnight culture was diluted 1:100 into 2 mL of fresh medium with the appropriate inducers and grown for 24 h unless specified otherwise. Then, where appropriate, 0.5 μL of culture was removed for fluorescence measure for flow cytometry, with the pre-lysis fluorescent distributions displayed in the same figure. For detection of protein expression, in most cases 1 mL of culture was harvested. Exception to the above applies to the split mCherry-split M86 intein splicing experiment, the mCherry BiFC experiment and the split TetR-SYNZIP experiment, where the volumes of bacterial culture harvested were adjusted by optical densities to standardize the amount of cellular materials used in cell lysis. Bacterial cells were centrifugation at 17, 000 × g and resuspended in 50 μL of 1× Laemmli sample buffer (1610747, Bio-Rad), boiled at 100 °C for 10 min, and centrifuged at 17, 000 × g for 10 min. 5 or 10 μL of the supernatant were resolved on an Any kD TGX Stain-Free protein gel (4568126, BioRad) alongside a Chameleon Duo Pre-Stained Protein Ladder (928-60000, Li-cor). Protein contents were then transferred to a nitrocellulose membrane (1704270, Bio-Rad) through a semi-dry transfer protocol (1704150, Bio-Rad). The manufacturer’s protocol (Doc. #988-13627, Licor) was followed for blocking, antibody incubation, washing and detection of near-infrared probes on secondary antibody. We used 5% (w/v) skimmed milk in 1× tris-buffered saline (1706435, Bio-Rad) as the blocking reagent. The mCherry constructs involved in the Western blot assays carried a hexahistidine tag at the C-terminus and an epitope (residues 27-41) exists within the N-lobe. Bipartite proteins were detected using the rabbit anti-mCherry (A00682, GenScript, 1:3000 diluted), the mouse anti-His (A00186, GenScript, 1:5000 diluted), and the rabbit anti-HA (902303, Biolegend, 1:1000 diluted) antibodies. They reacted against the IRDye 680RD goat antirabbit (925-68071, Licor) or the IRDye 800CW goat anti-mouse (925-32210, Li-cor) secondary antibodies, both diluted at 1:20,000. Membrane imaging was performed on the Odyssey CLx Infrared Imaging System (Li-cor) and the resulting images were processed using Image Studio Lite v5.2.5 software (Li-cor) and ImageJ^71^.

### Secondary structure alignments by amino acids and protein 3D structures

Secondary structures of SrpR and ECF20 were predicted using the JPred4 Server^43^, or modelled via SWISS-MODEL^72^. SrpR and TetR amino acid sequences were aligned with the known TetR structure (PDB: 4AC0) using PROMALS3D^45^ with default parameters. 3D structures were rendered using the software PyMOL v1.7.6.7 (Schrodinger).

### Library preparation, Next Generation Sequencing and data analysis

Glycerol stocks of the final library were thawed and for each library, 200 μL of the stock was inoculated into 50 mL of fresh medium for overnight growth. Subsequently the plasmid DNA libraries were extracted by midiprep. For IBM libraries of mCherry, TetR, SrpR and ECF20, appropriate combinations of restriction enzymes were used to release a minimal length of DNA fragments that contain mixed insertions at various positions. Digested DNA were resolved on agarose gels and the fragments with mixed insertions, which migrated as a single band, was excised and purified. For the domain-insertion library of ER-LBD into the M86 intein which was within mCherry, the region with insertions were amplified by PCR, then resolved and purified from agarose gels. Purified DNA were sent to Novogene (UK) for fragmentation and sequencing to obtain at least 7 million reads of 150 bp paired-end per library.

The resulting data was processed by in-house developed Python scripts. 12 bp, each at the 5’ and 3’ termini of the final inserted DNA were defined as signature sequences. Raw FASTQ files were filtered for reads that contained perfect matches to these signature sequences. Then, for each filtered read, the signature sequences were aligned and the adjacent sequence (12 bp) was extracted from the read, which were then aligned back to the CDS of the target protein to determine insertion positions. Only unique and perfect matches were considered authentic and a split/insertion site was called. Any fragments where the forward and reverse reads reported different split/insertion sites were removed, and fragments where the forward and reverse reads pointed to the same split/insertion site were deduplicated to avoid double counting. Rare instances of sites mapped beyond the permitted transposition window were also removed. Similar to previous works, a productive split or insertion site on the amino acid sequence was called only if the insertion orientation was forward and the insert was in-frame.

### Data Processing and Statistics

All data were processed and graphed in Python. Whenever displayed, fluorescence distributions shown within the same subpanel were normalized to their individual modes. We used two-tailed t-tests for independent samples assuming unequal variances in comparisons of fold changes between median fluorescent values, and in comparisons of median fluorescent values from populations. Calculation was done in Python using the SciPy package v1.4.1^73^. Owing to the large number of statistical tests performed within a single figure panel, we did not report the individual statistics and P-values but rather the summary statistics: n.s., not significant; *, *P* ≤ 0.05; **, *P* ≤ 0.01; ***, *P* ≤ 0.001. Exact sample sizes (n) were described in figure legends, except for **Supplementary Fig. 17** where sample sizes (n ≥ 3) of different sites differed greatly between groups and were too numerous to report as exact values.

## Data availability

Source data, including uncropped Western blot images and Python scripts for generating figures, are deposited to the Edinburgh DataShare (https://doi.org/10.7488/ds/2954). Raw sequencing data of IBM and DIM final libraries from NGS are deposited to the Sequence Read Archive under the project accession code PRJNA678813. List of constructs used in this study are detailed in Supplementary Data 1, and their sequences are available on SynBioHub^74^ (https://synbiohub.org/public/Intein assisted Bisection Mapping/Intein assisted Bisection Mapping coll ection/1). Representative key constructs used in this study, which allow researchers to conduct IBM of their own, are deposited at Addgene (ID 161937-161955). A detailed protocol for carrying out IBM is available at protocols.io (https://dx.doi.org/10.17504/protocols.io.bpqdmms6).

## Code availability

Python scripts for analyzing Sanger sequencing results to determine split sites at the final step of IBM, and for analyzing split or insertion site coverages from NGS data, is available at GitHub (https://github.com/tyhho/IBM).

## Acknowledgements

We thank Dr Martin Waterfall from the Flow Cytometry Core Facility at the University of Edinburgh for performing cells sorting, Dr Hung-Ju Chang and Dr Jerome Bonnet for advice on acVHH-related experiments, Prof. Bryan C Dickinson for providing plasmids encoding the evolved split T7 RNA polymerase, members of the Wang and Menolascina labs for helpful discussions, and Dr. Jamie Gilman and Dr. James Bryson for feedback on the manuscript. We would also like to thank the SBS BioRDM Team for their help with data deposits and data curation. This work was supported by the UK Research and Innovation Future Leaders Fellowship [MR/S018875/1], Leverhulme Trust grant [RPG-2020-241] and UK Biotechnology and Biological Sciences Research Council grant [BB/N007212/1] to B.W.. T.Y.H.H. was supported by the Darwin Trust of Edinburgh and the Edinburgh Global Research Scholarship. A.S. was supported by a Microsoft Research PhD Scholarship.

## Author Contributions

B.W. and T.Y.H.H conceived the study. T.Y.H.H developed the methods and designed the experiments with inputs from B.W.. T.Y.H.H, A.S., and Z.L. performed the experiments. T.Y.H.H. analyzed the data with inputs from B.W., N.D., L.W. and F.M.. H.S. provided the reagents for transposition. All authors took part in the interpretation of results and preparation of materials for the manuscript. T.Y.H.H. and B.W. wrote the manuscript with comments from all authors. B.W. supervised and acquired the funding of the study.

## References

1. Shekhawat, S.S. & Ghosh, I. Split-protein systems: beyond binary protein-protein interactions. Current Opinion in Chemical Biology 15, 789–797 (2011).

2. Kamiyama, D. et al. Versatile protein tagging in cells with split fluorescent protein. Nature Communications 7, 11046 (2016).

3. Guo, Z. et al. Engineered PQQ-Glucose Dehydrogenase as a Universal Biosensor Platform. Journal of the American Chemical Society 138, 10108–10111 (2016).

4. Hicks, M., Bachmann, T.T. & Wang, B. Synthetic Biology Enables Programmable Cell-Based Biosensors. ChemPhysChem 21, 132–144 (2020).

5. Gray, D.C., Mahrus, S. & Wells, J.A. Activation of Specific Apoptotic Caspases with an Engineered SmallMolecule-Activated Protease. Cell 142, 637–646 (2010).

6. Weinberg, B.H. et al. High-performance chemical- and light-inducible recombinases in mammalian cells and mice. Nature Communications 10, 4845 (2019).

7. Gao, X.J., Chong, L.S., Kim, M.S. & Elowitz, M.B. Programmable protein circuits in living cells. Science 361, 1252–1258 (2018).

8. Fink, T. et al. Design of fast proteolysis-based signaling and logic circuits in mammalian cells. Nature Chemical Biology 15, 115–122 (2019).

9. Xiang, Y., Dalchau, N. & Wang, B. Scaling up genetic circuit design for cellular computing: advances and prospects. Natural Computing 17, 833–853 (2018).

10. Wang, B., Kitney, R.I., Joly, N. & Buck, M. Engineering modular and orthogonal genetic logic gates for robust digital-like synthetic biology. Nature Communications 2, 508 (2011).

11. Schopp, I.M. et al. Split-BioID a conditional proteomics approach to monitor the composition of spatiotemporally defined protein complexes. Nature Communications 8, 15690 (2017).

12. Golob-Urbanc, A., Rajčević, U., Strmšek, Ž. & Jerala, R. Design of split superantigen fusion proteins for cancer immunotherapy. Journal of Biological Chemistry 294, 6294–6305 (2019).

13. Ma, D., Peng, S. & Xie, Z. Integration and exchange of split dCas9 domains for transcriptional controls in mammalian cells. Nature Communications 7, 13056 (2016).

14. Shah, N.H. & Muir, T.W. Inteins: nature’s gift to protein chemists. Chemical Science 5, 446–461 (2014).

15. Elleuche, S. & Pöggeler, S. Inteins, valuable genetic elements in molecular biology and biotechnology. Applied Microbiology and Biotechnology 87, 479–489 (2010).

16. Pinto, F., Thornton, E.L. & Wang, B. An expanded library of orthogonal split inteins enables modular multi-peptide assemblies. Nature Communications 11, 1529 (2020).

17. Slomovic, S. & Collins, J.J. DNA sense-and-respond protein modules for mammalian cells. Nature Methods 12, 1085–1090 (2015).

18. Schaerli, Y., Gili, M. & Isalan, M. A split intein T7 RNA polymerase for transcriptional AND-logic. Nucleic Acids Research 42, 12322–12328 (2014).

19. Bradley, R.W., Buck, M. & Wang, B. Tools and Principles for Microbial Gene Circuit Engineering. Journal of Molecular Biology 428, 862–888 (2016).

20. Wang, T., Badran, A.H., Huang, T.P. & Liu, D.R. Continuous directed evolution of proteins with improved soluble expression. Nature Chemical Biology 14, 972–980 (2018).

21. Amitai, G., Callahan, B.P., Stanger, M.J., Belfort, G. & Belfort, M. Modulation of intein activity by its neighboring extein substrates. Proceedings of the National Academy of Sciences 106, 11005–11010 (2009).

22. Shah, N.H., Eryilmaz, E., Cowburn, D. & Muir, T.W. Extein Residues Play an Intimate Role in the RateLimiting Step of Protein Trans-Splicing. Journal of the American Chemical Society 135, 5839–5847 (2013).

23. Davis, K.M., Pattanayak, V., Thompson, D.B., Zuris, J.A. & Liu, D.R. Small molecule-triggered Cas9 protein with improved genome-editing specificity. Nature Chemical Biology 11, 316 (2015).

24. Jillette, N., Du, M., Zhu, J.J., Cardoz, P. & Cheng, A.W. Split selectable markers. Nature Communications 10, 4968 (2019).

25. Palanisamy, N. et al. Split intein-mediated selection of cells containing two plasmids using a single antibiotic. Nature Communications 10, 4967 (2019).

26. Dagliyan, O. et al. Computational design of chemogenetic and optogenetic split proteins. Nature Communications 9, 4042 (2018).

27. Segall-Shapiro, T.H., Meyer, A.J., Ellington, A.D., Sontag, E.D. & Voigt, C.A. A ‘resource allocator’ for transcription based on a highly fragmented T7 RNA polymerase. Molecular Systems Biology 10, 742 (2014).

28. Nadler, D.C., Morgan, S.-A., Flamholz, A., Kortright, K.E. & Savage, D.F. Rapid construction of metabolite biosensors using domain-insertion profiling. Nature Communications 7, 12266 (2016).

29. Oakes, B.L. et al. Profiling of engineering hotspots identifies an allosteric CRISPR-Cas9 switch. Nature Biotechnology 34, 646 (2016).

30. Zeng, Y. et al. A Split Transcriptional Repressor That Links Protein Solubility to an Orthogonal Genetic Circuit. ACS Synthetic Biology 7, 2126–2138 (2018).

31. Calles, B. & Lorenzo, V.d. Expanding the Boolean Logic of the Prokaryotic Transcription Factor XylR by Functionalization of Permissive Sites with a Protease-Target Sequence. ACS Synthetic Biology 2, 594603 (2013).

32. Appleby-Tagoe, J.H. et al. Highly Efficient and More General cis- and trans-Splicing Inteins through Sequential Directed Evolution. Journal of Biological Chemistry 286, 34440–34447 (2011).

33. Fan, J.-Y. et al. Split mCherry as a new red bimolecular fluorescence complementation system for visualizing protein-protein interactions in living cells. Biochemical and Biophysical Research Communications 367, 47–53 (2008).

34. Thompson, K.E., Bashor, C.J., Lim, W.A. & Keating, A.E. SYNZIP Protein Interaction Toolbox: in Vitro and in Vivo Specifications of Heterospecific Coiled-Coil Interaction Domains. ACS Synthetic Biology 1, 118129 (2012).

35. Carvajal-Vallejos, P., Pallissé, R., Mootz, H.D. & Schmidt, S.R. Unprecedented Rates and Efficiencies Revealed for New Natural Split Inteins from Metagenomic Sources. Journal of Biological Chemistry 287, 28686–28696 (2012).

36. Galarneau, A., Primeau, M., Trudeau, L.-E. & Michnick, S.W. β-Lactamase protein fragment complementation assays as in vivo and in vitro sensors of protein-protein interactions. Nature Biotechnology 20, 619–622 (2002).

37. Wehrman, T., Kleaveland, B., Her, J.-H., Balint, R.F. & Blau, H.M. Protein-protein interactions monitored in mammalian cells via complementation of β-lactamase enzyme fragments. Proceedings of the National Academy of Sciences 99, 3469–3474 (2002).

38. Beyer, H.M., Mikula, K.M., Li, M., Wlodawer, A. & Iwaï, H. The crystal structure of the naturally split gp41-1 intein guides the engineering of orthogonal split inteins from cis-splicing inteins. The FEBS Journal 287, 1886–1898 (2020).

39. Wang, B., Barahona, M. & Buck, M. A modular cell-based biosensor using engineered genetic logic circuits to detect and integrate multiple environmental signals. Biosensors and Bioelectronics 40, 368–376 (2013).

40. Wang, B. & Buck, M. Rapid engineering of versatile molecular logic gates using heterologous genetic transcriptional modules. Chemical Communications 50, 11642–11644 (2014).

41. Stanton, B.C. et al. Genomic mining of prokaryotic repressors for orthogonal logic gates. Nature Chemical Biology 10, 99–105 (2014).

42. Rhodius, V.A. et al. Design of orthogonal genetic switches based on a crosstalk map of σs, anti-σs, and promoters. Molecular Systems Biology 9, 702 (2013).

43. Drozdetskiy, A., Cole, C., Procter, J. & Barton, G.J. JPred4: a protein secondary structure prediction server. Nucleic Acids Research 43, W389–W394 (2015).

44. Aranko, A.S., Wlodawer, A. & Iwaï, H. Nature’s recipe for splitting inteins. Protein Engineering, Design and Selection 27, 263–271 (2014).

45. Pei, J., Kim, B.-H. & Grishin, N.V. PROMALS3D: a tool for multiple protein sequence and structure alignments. Nucleic Acids Research 36, 2295–2300 (2008).

46. Mootz, H.D. & Muir, T.W. Protein Splicing Triggered by a Small Molecule. Journal of the American Chemical Society 124, 9044–9045 (2002).

47. Mootz, H.D., Blum, E.S., Tyszkiewicz, A.B. & Muir, T.W. Conditional Protein Splicing: A New Tool to Control Protein Structure and Function in Vitro and in Vivo. Journal of the American Chemical Society 125, 10561–10569 (2003).

48. Di Ventura, B. & Mootz, H.D. Switchable inteins for conditional protein splicing. Biological Chemistry 400, 467–475 (2019).

49. Peck, Sun H., Chen, I. & Liu, David R. Directed Evolution of a Small-Molecule-Triggered Intein with Improved Splicing Properties in Mammalian Cells. Chemistry & Biology 18, 619–630 (2011).

50. Skretas, G. & Wood, D.W. Regulation of protein activity with small-molecule-controlled inteins. Protein Science 14, 523–532 (2005).

51. Gramespacher, J.A., Burton, A.J., Guerra, L.F. & Muir, T.W. Proximity Induced Splicing Utilizing Caged Split Inteins. Journal of the American Chemical Society 141, 13708–13712 (2019).

52. Bojar, D., Scheller, L., Hamri, G.C.-E., Xie, M. & Fussenegger, M. Caffeine-inducible gene switches controlling experimental diabetes. Nature Communications 9, 2318 (2018).

53. Chang, H.-J. et al. A Modular Receptor Platform To Expand the Sensing Repertoire of Bacteria. ACS Synthetic Biology 7, 166–175 (2018).

54. Pu, J., Zinkus-Boltz, J. & Dickinson, B.C. Evolution of a split RNA polymerase as a versatile biosensor platform. Nature Chemical Biology 13, 432–438 (2017).

55. Stevens, A.J. et al. A promiscuous split intein with expanded protein engineering applications. Proceedings of the National Academy of Sciences 114, 8538–8543 (2017).

56. Dagliyan, O. et al. Rational design of a ligand-controlled protein conformational switch. Proceedings of the National Academy of Sciences 110, 6800–6804 (2013).

57. Karginov, A.V., Ding, F., Kota, P., Dokholyan, N.V. & Hahn, K.M. Engineered allosteric activation of kinases in living cells. Nature Biotechnology 28, 743–747 (2010).

58. Buskirk, A.R., Ong, Y.-C., Gartner, Z.J. & Liu, D.R. Directed evolution of ligand dependence: Smallmolecule-activated protein splicing. Proceedings of the National Academy of Sciences of the United States of America 101, 10505–10510 (2004).

59. Brödel, A.K., Jaramillo, A. & Isalan, M. Intracellular directed evolution of proteins from combinatorial libraries based on conditional phage replication. Nature Protocols 12, 1830–1843 (2017).

60. Mizuuchi, M. & Mizuuchi, K. Target Site Selection in Transposition of Phage Mu. Cold Spring Harbor Symposia on Quantitative Biology 58, 515–523 (1993).

61. Haapa-Paananen, S., Rita, H. & Savilahti, H. DNA Transposition of Bacteriophage Mu: A QUANTITATIVE ANALYSIS OF TARGET SITE SELECTION IN VITRO. Journal of Biological Chemistry 277, 2843–2851 (2002).

62. Manna, D., Deng, S., Breier, A.M. & Higgins, N.P. Bacteriophage Mu Targets the Trinucleotide Sequence CGG. Journal of Bacteriology 187, 3586–3588 (2005).

63. Green, B., Bouchier, C., Fairhead, C., Craig, N.L. & Cormack, B.P. Insertion site preference of Mu, Tn5, and Tn7 transposons. Mobile DNA 3, 3 (2012).

64. Coyote-Maestas, W., Nedrud, D., Okorafor, S., He, Y. & Schmidt, D. Targeted insertional mutagenesis libraries for deep domain insertion profiling. Nucleic Acids Research 48, e11–e11 (2019).

65. Atkinson, J.T., Jones, A.M., Zhou, Q. & Silberg, J.J. Circular permutation profiling by deep sequencing libraries created using transposon mutagenesis. Nucleic Acids Research 46, e76–e76 (2018).

66. Baker, T.A., Mizuuchi, M., Savilahti, H. & Mizuuchi, K. Division of labor among monomers within the Mu transposase tetramer. Cell 74, 723–733 (1993).

67. Haapa-Paananen, S. & Savilahti, H. Applications of the Bacteriophage Mu In Vitro Transposition Reaction and Genome Manipulation via Electroporation of DNA Transposition Complexes. Methods in Molecular Biology 1681, 279–286 (2018).

68. Engler, C., Kandzia, R. & Marillonnet, S. A One Pot, One Step, Precision Cloning Method with High Throughput Capability. PLOS ONE 3, e3647 (2008).

69. Cock, P.J.A. et al. Biopython: freely available Python tools for computational molecular biology and bioinformatics. Bioinformatics 25, 1422–1423 (2009).

70. Kunzmann, P. & Hamacher, K. Biotite: a unifying open source computational biology framework in Python. BMC Bioinformatics 19, 346 (2018).

71. Rueden, C.T. et al. ImageJ2: ImageJ for the next generation of scientific image data. BMC Bioinformatics 18, 529 (2017).

72. Bienert, S. et al. The SWISS-MODEL Repository—new features and functionality. Nucleic Acids Research 45, D313–D319 (2016).

73. Virtanen, P. et al. SciPy 1.0: fundamental algorithms for scientific computing in Python. Nature Methods 17, 261–272 (2020).

74. McLaughlin, J.A. et al. SynBioHub: A Standards-Enabled Design Repository for Synthetic Biology. ACS Synthetic Biology 7, 682–688 (2018).

