## Supplementary Information File for "Intein-assisted bisection mapping systematically splits proteins for Boolean logic and inducibility engineering"

**Supplementary Figures and Table**

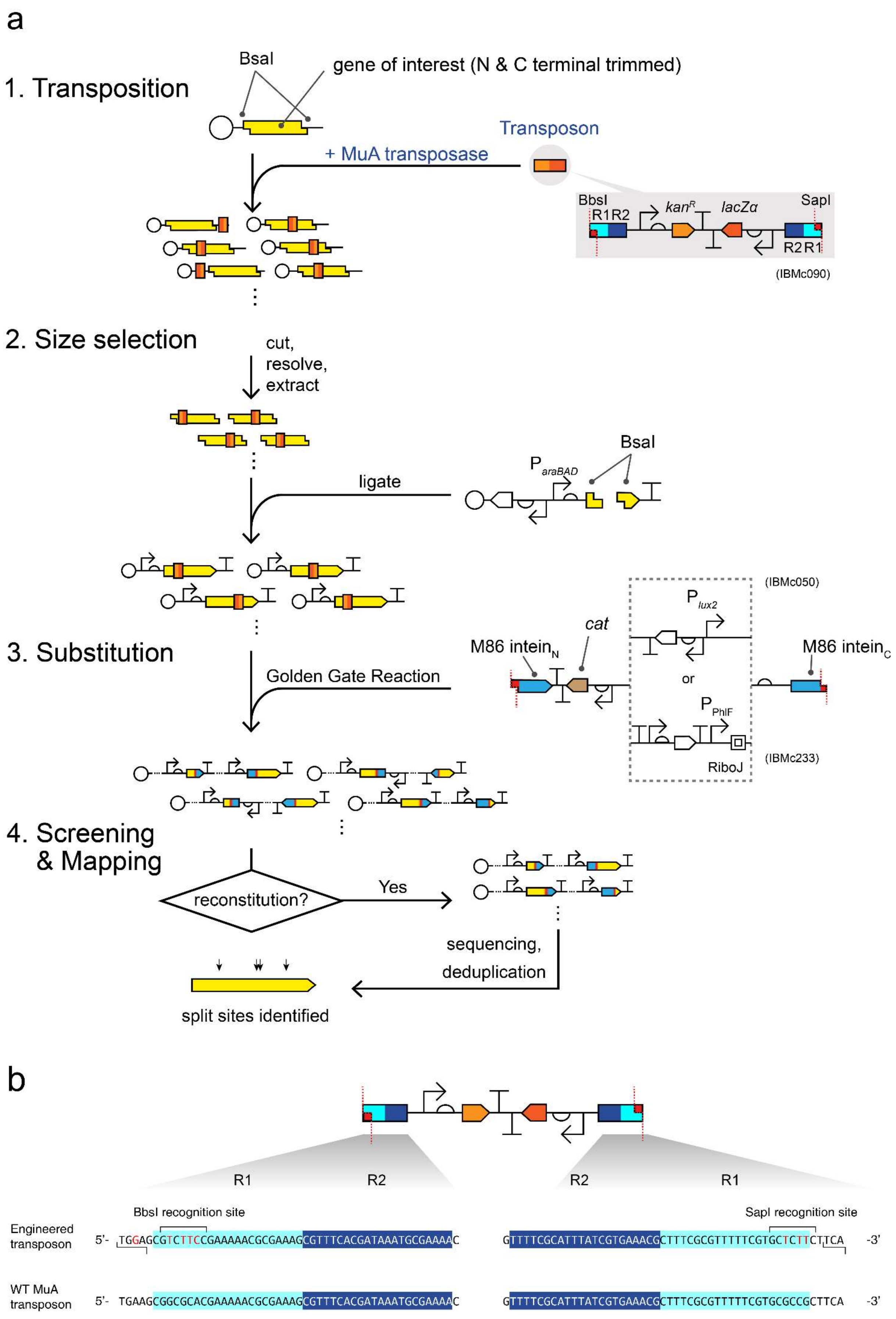

a
1. Transposition
BsaI
gene of interest (N & C terminal trimmed)
Transposon
+ MuA transposase
BbsI
R1R2
kanR
lacZα
SapI
R2R1
(IBMc090)
2. Size selection
cut,
resolve,
extract
BsaI
ParaBAD
ligate
3. Substitution
Golden Gate Reaction
M86 inteinN
cat
Plux2
or
PPhlF
RiboJ
(IBMc050)
(IBMc233)
M86 inteinC
4. Screening
& Mapping
reconstitution?
Yes
sequencing,
deduplication
split sites identified
b
R1
R2
R2
R1
BbsI recognition site
SapI recognition site
Engineered transposon
5'- TGGAGCGTCTTCCGAAAAACGCGAAAGCGTTTCACGATAAATGCGAAAAC
GTTTTCGCATTTATCGTGAAACGCTTTCGCGTTTTTCGTGCTCTTCTTCA -3'
WT MuA transposon
5'- TGAAGCGGCGCACGAAAAACGCGAAAGCGTTTCACGATAAATGCGAAAAC
GTTTTCGCATTTATCGTGAAACGCTTTCGCGTTTTTCGTGCGCCGCTTCA -3'

**Supplementary Figure 1 Workflow of intein-assisted bisection mapping (IBM). a.** Execution-wise, IBM is an augmentation of the bisection mapping method by adding a Golden Gate substitution step. It starts with a transposition reaction: a staging plasmid carries a 5' and 3' trimmed gene of interest (GOI) with the internal BsaI, BbsI and SapI sites removed. The staging plasmid is mixed in vitro with the MuA transposase and the mini-Mu transposon[1, 2], which generates the initial insertion library. After in vivo amplification (transformation into *E. coli* and overnight growth), the insertion library is cut by BsaI and resolved on a DNA agarose gel. The band with the correct size corresponding to GOI with the insertion is purified and ligated into a linearized vector with BsaI-generated overhangs. This produces the Open Reading Frame (ORF) insertion library[3]. The ORF insertion library is again amplified in vivo and then an aliquot is mixed with a DNA fragment carrying the split *Ssp* DnaB$^{M86}$ intein in a Golden Gate reaction, using the restriction enzymes BbsI and SapI. In later experiments we replaced the $P_{lux2}$ promoter by the $P_{PhlF}$ promoter which was reported to be very tight[4]. The DNA fragment contains a different selection marker (*cat*) to facilitate selection of the library with the transposon replaced. In the end, the final library is screened for individual strains that showed proper reconstitution of protein function upon expression of both the N- and C-lobes (AND or NAND logic gate behavior). Those strains were then subjected to Sanger sequencing at one of the joints to map the split sites. Identical split sites were then deduplicated and consolidated with activity data to generate the intein-bisection maps. **b.** Illustration of modifications on wild type (WT) Mu transposon R1R2 recognition sequences[5] to accommodate BbsI and SapI Type IIS restriction sites. R1 sequences are highlighted in cyan and R2, dark blue. Mutations introduced are colored in red. Recognition sites are marked by square brackets and overhangs positions are denoted by staggered lines.

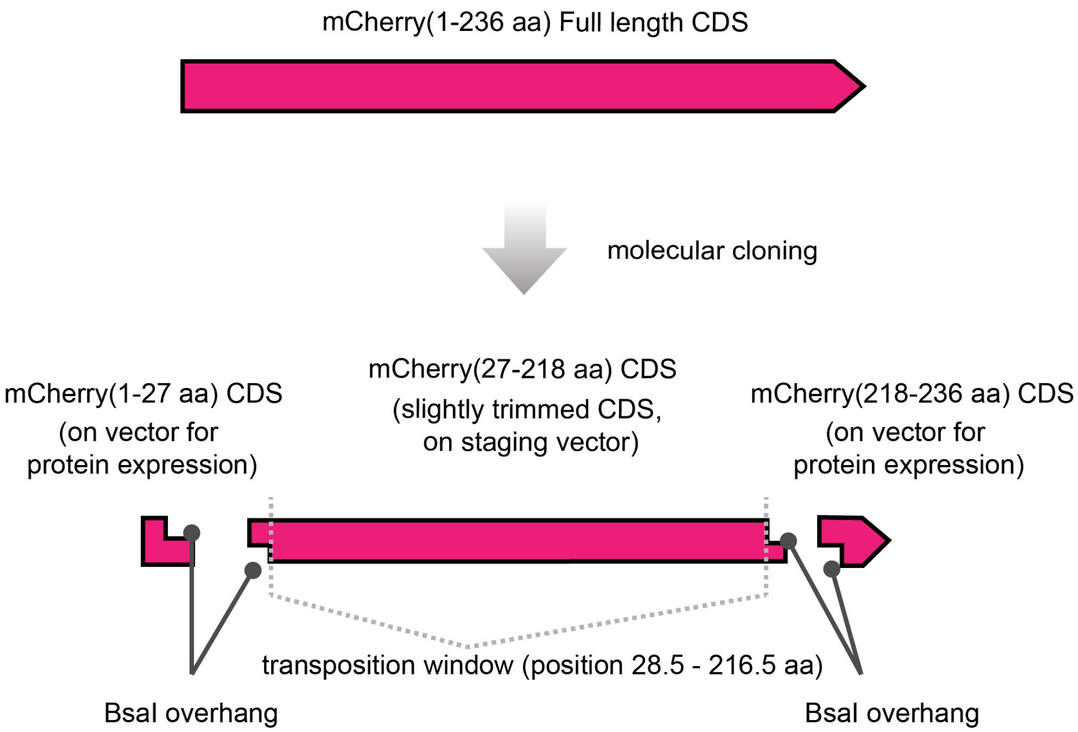

**Supplementary Figure 2 Illustration on slightly trimmed CDS using mCherry as an example.** Before IBM was performed on a protein of interest, most part of its CDS, excluding the very N terminus and the C terminus, would be subcloned onto a staging plasmid and flanked by BsaI sites. If transposition happens within the BsaI sites or beyond them, the transposed fragments would be unable to be selected for before being subcloned into the protein expression plasmid. The sequence spaces between the two BsaI sites thus defines the transposition window.

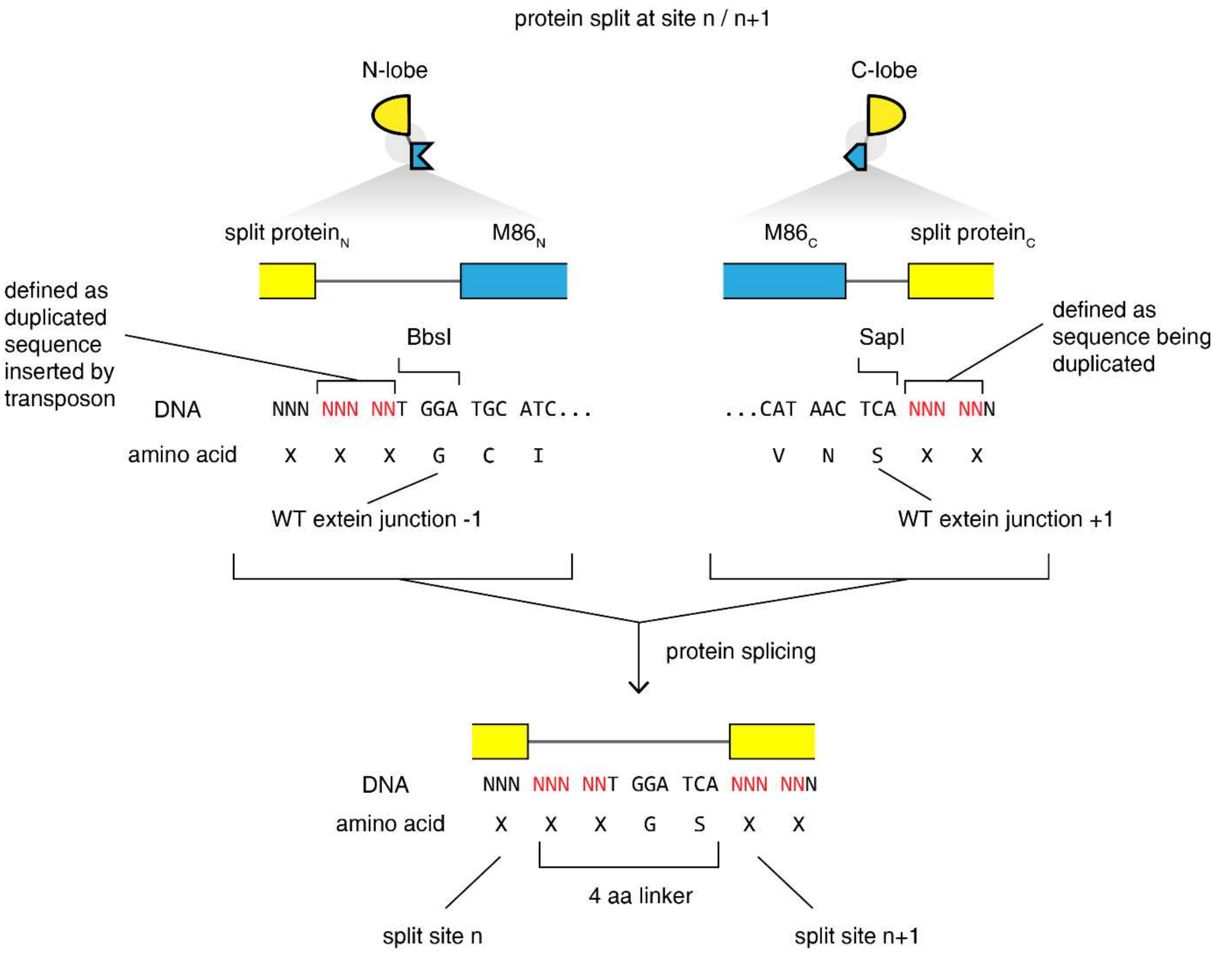

**Supplementary Figure 3 Illustration of effect on transposition and Golden Gate substitution on amino acid linker in spliced product from IBM.** The BbsI and SapI overhangs are at the very ends of the R1 recognition sites on the mini-Mu transposon. As a result, the number of amino acids inserted at each junction, between the extein and the intein, is minimal. Transposition by MuA duplicates 5 bp at the insertion site (colored in red)[2]. The duplicated sequence at the 3' end is arbitrary defined as the part of the original sequence and that at the 5' is defined as the extra bases. The extra bases together with the DNA of the BbsI overhang code for three amino acid residues, which includes the -1 extein junction for the M86 intein[6]. At the 3' end, bases of the SapI overhang encodes a serine, which is the required +1 extein junction. Splicing thus leaves behind a 4 bp scar.

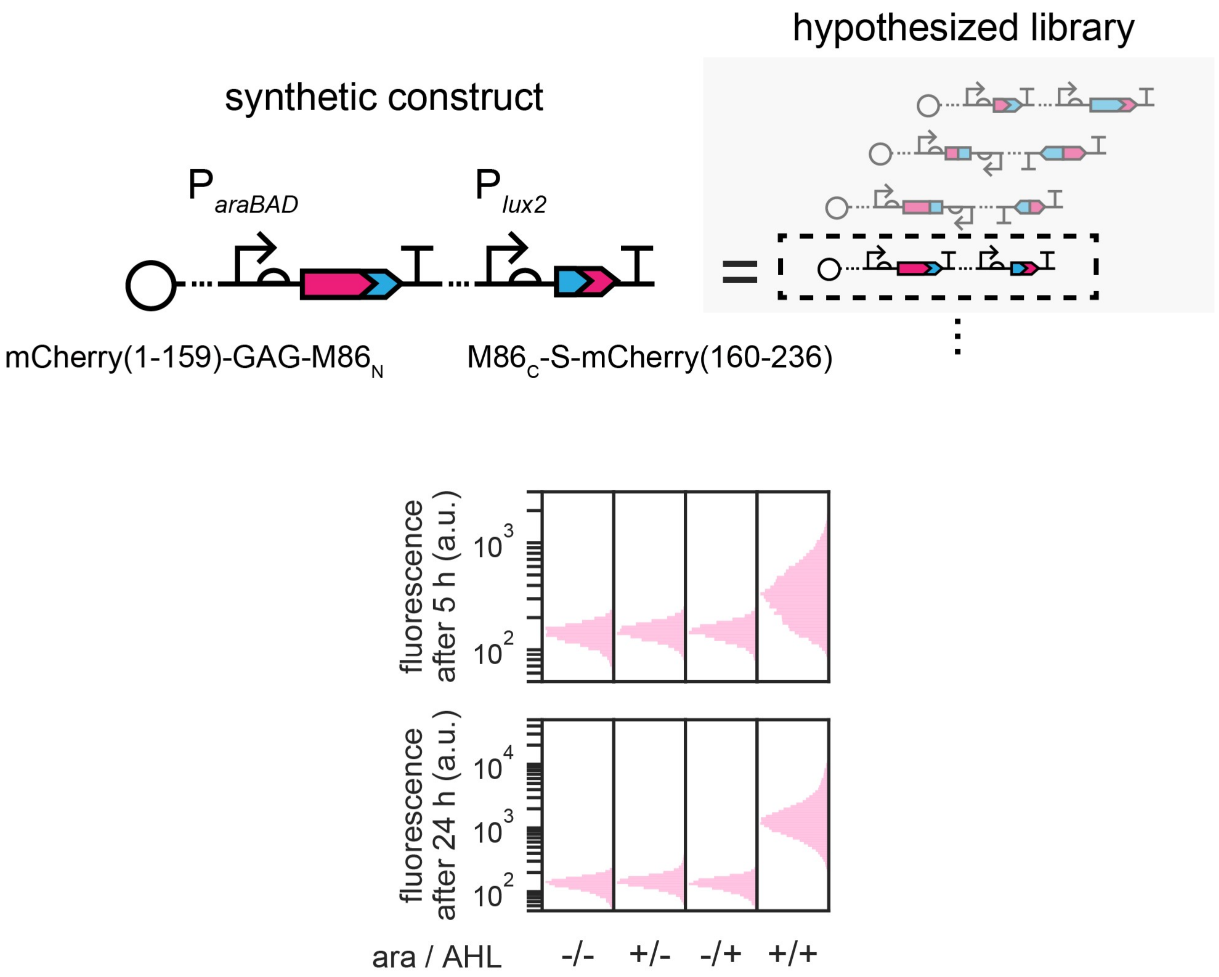

**Supplementary Figure 4 Outcome simulating control suggested that IBM should recover split site 159/160 on mCherry.** A synthetic construct was built to mimic one of the members within the final library if IBM had been performed on mCherry. mCherry would have been split at site 156/160, with a total of four extra amino acid residues added at the split site. This construct yielded the correct AND logic gate behavior with increased fluorescence only when both the N- and C-lobes were present. Proper execution of IBM on mCherry should in principle recover this construct and its split site. Inducers for N- and C-lobes were arabinose (ara, 1 mM) and acyl homoserine lactone (AHL, 1 µM). Single-cell fluorescence distributions shown were pooled from three biological replicates.

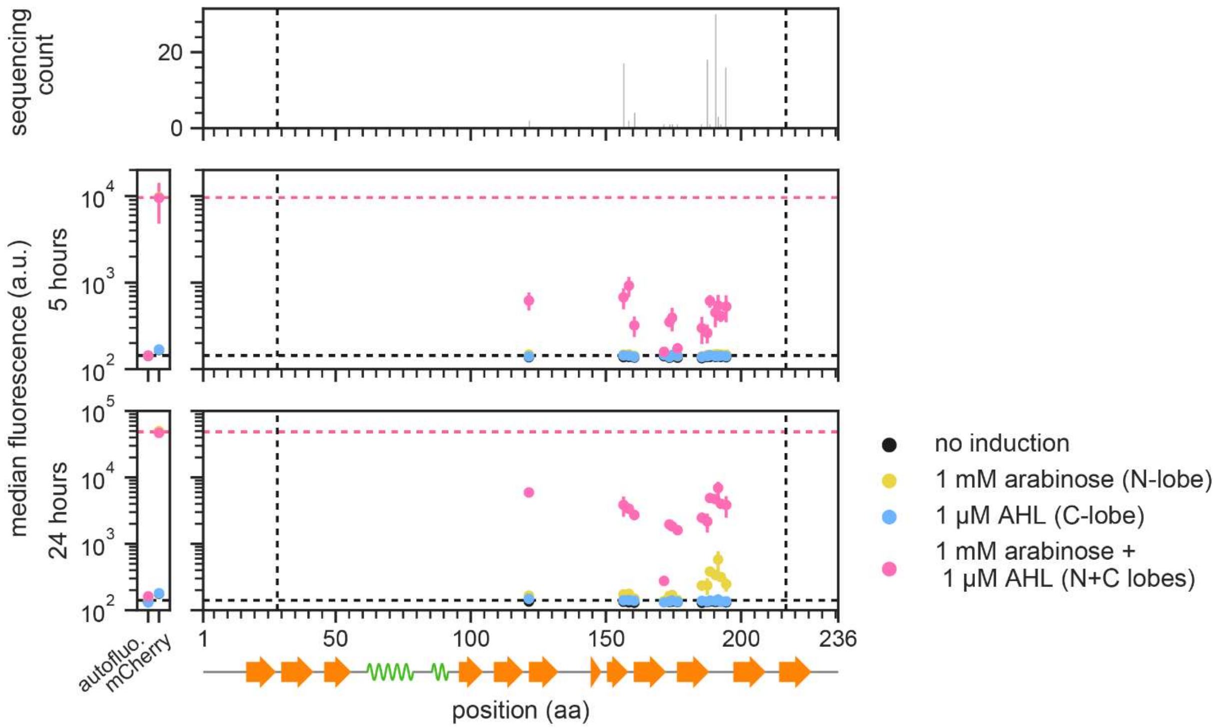

**Supplementary Figure 5 Full intein-bisection map of mCherry.** Full map of IBM on mCherry in which data for Figure 1b was drawn from. Left panel, the fluorescence of the controls that provide the references (horizontal dashed lines) for activities of the intact mCherry and the minimal activity in theory achievable by bipartite constructs. Right panel, bisection map of mCherry. Each vertical group of spots represents an identified split site on the x axis, aligned to the mCherry secondary structure (PDB: 2H5Q)[7] below. A total of 99 filtered candidate strains were characterized and sequenced to generate this map. y locations and error bars are mean and std of median fluorescence from independent experiments performed on three different days. Vertical dashed lines bound the permitted transposition window.

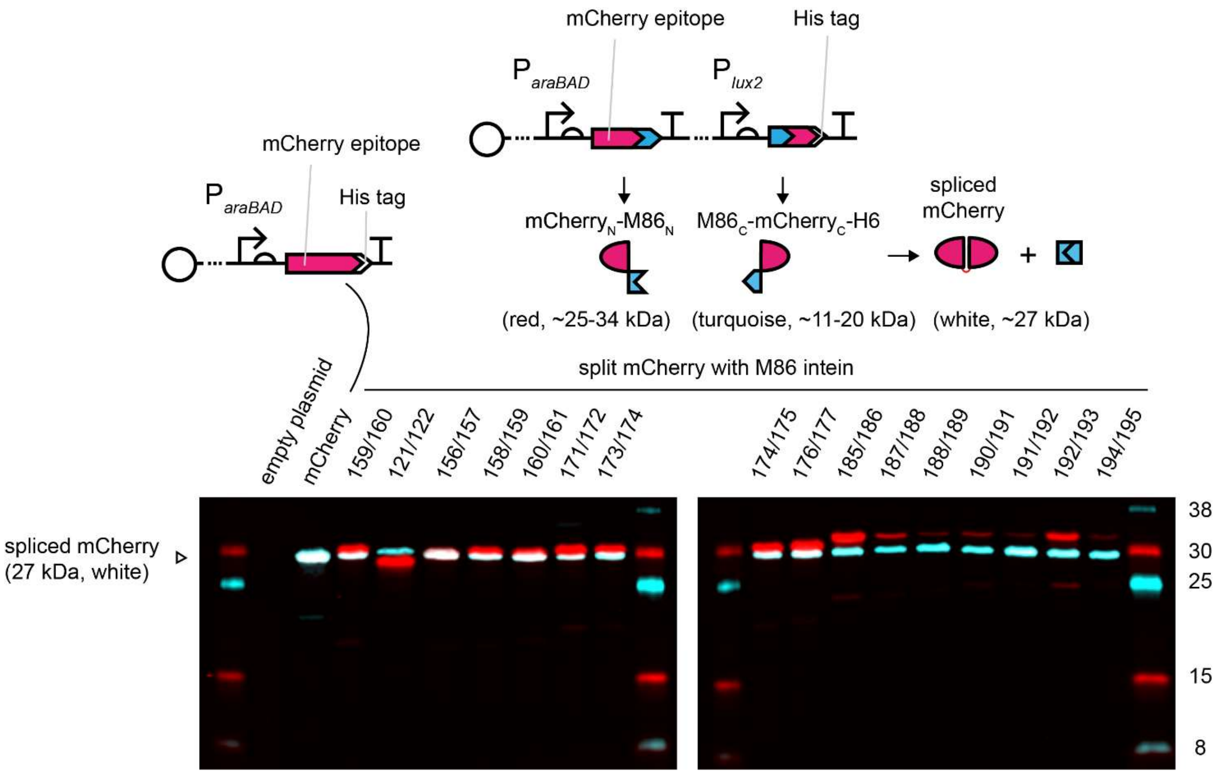

**Supplementary Figure 6 Proof of mCherry splicing at all split sites identified from IBM of mCherry for BiFC.** For all split sites identified for mCherry from IBM (Fig. 1b), a hexahistidine tag was added to the C-terminus of the split mCherry C-lobes by molecular cloning. Cells harboring the constructs were grown for 5 hours in the presence of 1 mM arabinose and 1 µM AHL, and were then harvested for cell lysis. Whole-cell lysates were subjected to a Western blots experiment. N- and C-lobes expression were probed using antibodies that target the mCherry epitope (red) and the hexahistidine tag (turquoise) respectively. In all constructs, the mCherry epitope is located within the N-lobes despite differences in split sites. Overlaps of red and turquoise bands gave white bands at a size between 25 and 30 kDa, indicating splice products formation. In each lane, a second red band with sizes corresponding to unspliced precursor could be found but only a turquoise band, at the size of the splice product, was present. This suggested the C-lobes were the limiting precursors in splicing and N-lobes were in excess. Our result proves that splicing has occurred at all split sites identified from IBM.

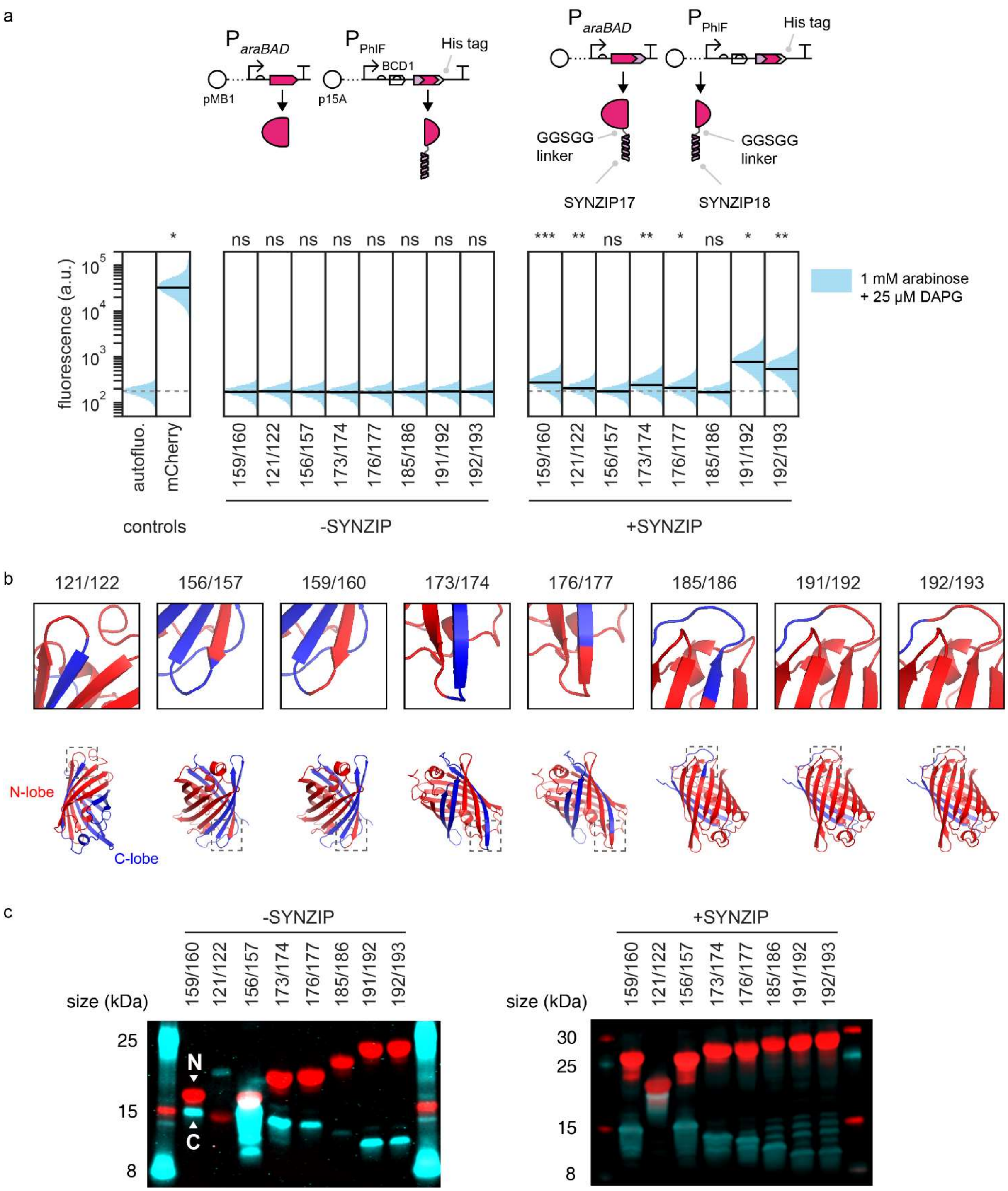
a
P araBAD
P PhlF
BCD1
His tag
pMB1
p15A
GGSGG linker
SYNZIP17
SYNZIP18
fluorescence (a.u.)
1 mM arabinose + 25 μM DAPG
autofluo.
mCherry
159/160
121/122
156/157
173/174
176/177
185/186
191/192
192/193
controls
-SYNZIP
+SYNZIP
*
ns
***
**
b
N-lobe
C-lobe
c
size (kDa)
N
C
25
15
8
30

**Supplementary Figure 7 Backward compatibility of split sites from IBM of mCherry for BiFC. a.** To test if split sites identified from IBM of mCherry were functional for the purpose of bimolecular fluorescence complementation (BiFC), mCherry was split at representative sites (121/122, 173/174, 191/192, 192/193) and fused with synthetic coil-coiled domains SYNZIP17 and SYNZIP18[8] through a 5-residue linker (right most panel). To test if the split mCherry alone could self-dimerize, the same assay was conducted, in which SYNZIP17 on the N-lobes were removed but SYNZIP18 were retained on the C-lobes (middle panel). In addition, some IBM-identified split sites locating within β-sheets (156/157, 176/177, 185/186) were tested. Cells were induced with 1 mM arabinose and 25 μM DAPG were grown for 16 h at 37 °C followed by incubation at room temperature for 9 h to maximize signals from complementation. Single-cell fluorescence distributions shown were pooled from three biological replicates. Solid black horizontal lines denote population median, except for autofluorescence which was denoted by dotted grey lines. Above each histogram, the statistics summary is a two-tailed t-test, assuming unequal variance, that compares the median fluorescence values between the test population and the autofluorescence populations (n = 3). BCD, bicistronic design to increase the likelihood that the C-lobes would be expressed at similar levels despite having different 5' coding DNA sequences[9]. Note that the same data for constructs with SYNZIP was used to generate **Fig. 1d**. In all constructs, mCherry carried a hexahistidine tag at the C-terminus. **b.** Illustration of chosen split sites on mChery 3D structures (PDB:2H5Q). Top panels in boxes are zoom-in views of subsections bounded by grey dashed lines on the bottom panel, which provide views to the whole structure. As shown, sites 121/122, 159/160, 173/174, 191/192, 192/193 are on loops and sites 156/157, 176/177, 185/186 are within β-sheets. **c**. The Western blot result on whole-cell lysates of cells from **a** on one of the three replicates. N- and C-lobes expression were probed using antibodies that target the mCherry epitope (red) and the hexahistidine tag (turquoise) respectively. In all constructs, the mCherry epitope is located within the N-lobes despite differences in split sites. This proves that lack of fluorescence from constructs missing SYNZIP17 was not due to absence of protein expression. It should be noted that, in the cases where SYNZIP17 was removed from the N-lobes, the C-lobes were generally more poorly expressed than the N-lobes and so the brightness of the turquoise channel was enhanced to visualize expression of the C-lobes. **a-c**. Split site 159/160 was not recovered from the IBM of mCherry but served as a reference of a known split site for comparison.

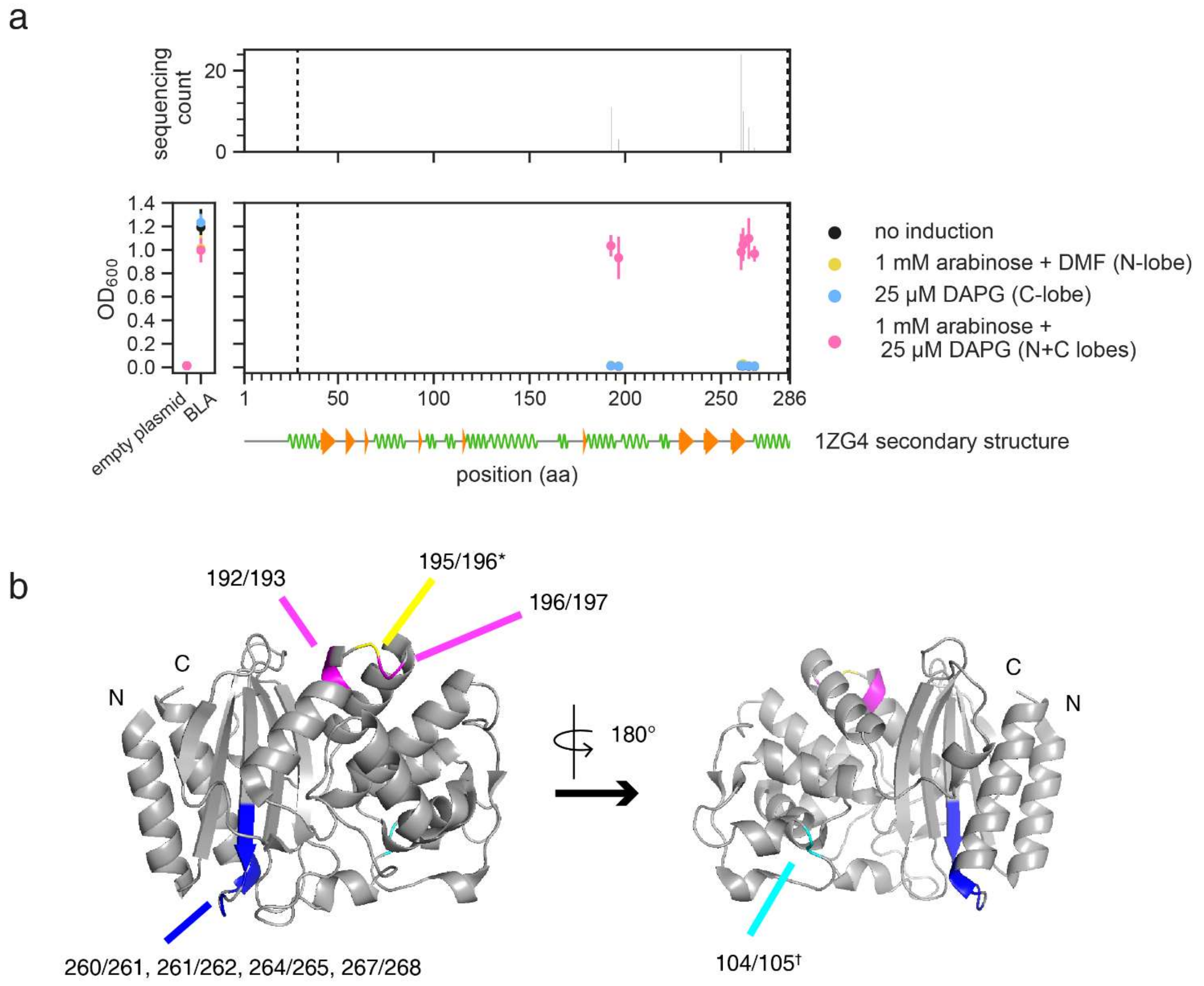

**Supplementary Figure 8 IBM on β-lactamase identified a new computationally unpredicted split site. a.** The intein-bisection map of β-lactamase. **a.** Left panel, optical densities of the controls that provide the references as to how cells with or without intact β-lactamase (BLA) would grow when challenged by ampicillin. Right panel, bisection map of β-lactamase split by the split gp41-1 intein. Each vertical group of spots represents an identified split site on the x axis, aligned to the β-lactamase secondary structure (from PDB: 1ZG4) below. A total of 55 candidate strains showing AND gate behavior were characterized and sequenced to generate this map. y locations and error bars are mean and std of optical densities from independent experiments performed on three different days. Vertical dashed lines bound the permitted transposition window. **b.** Known and IBM-identified split sites of β-lactamase mapped to the crystal structure (PDB: 1ZG4). Each split site has the -1 and the +1 amino acid residues colored. Sites 192/193 and 196/197 were colored in magenta, 260/261, 261/262, 264/265, 267/268 were colored in blue. (*) The established site 195/196[10, 11] was between the yellow and magenta colored residues. (†) The computationally predicted and then experimentally verified split site 104/105[12] was colored in cyan. It was not identified from this IBM experiment.

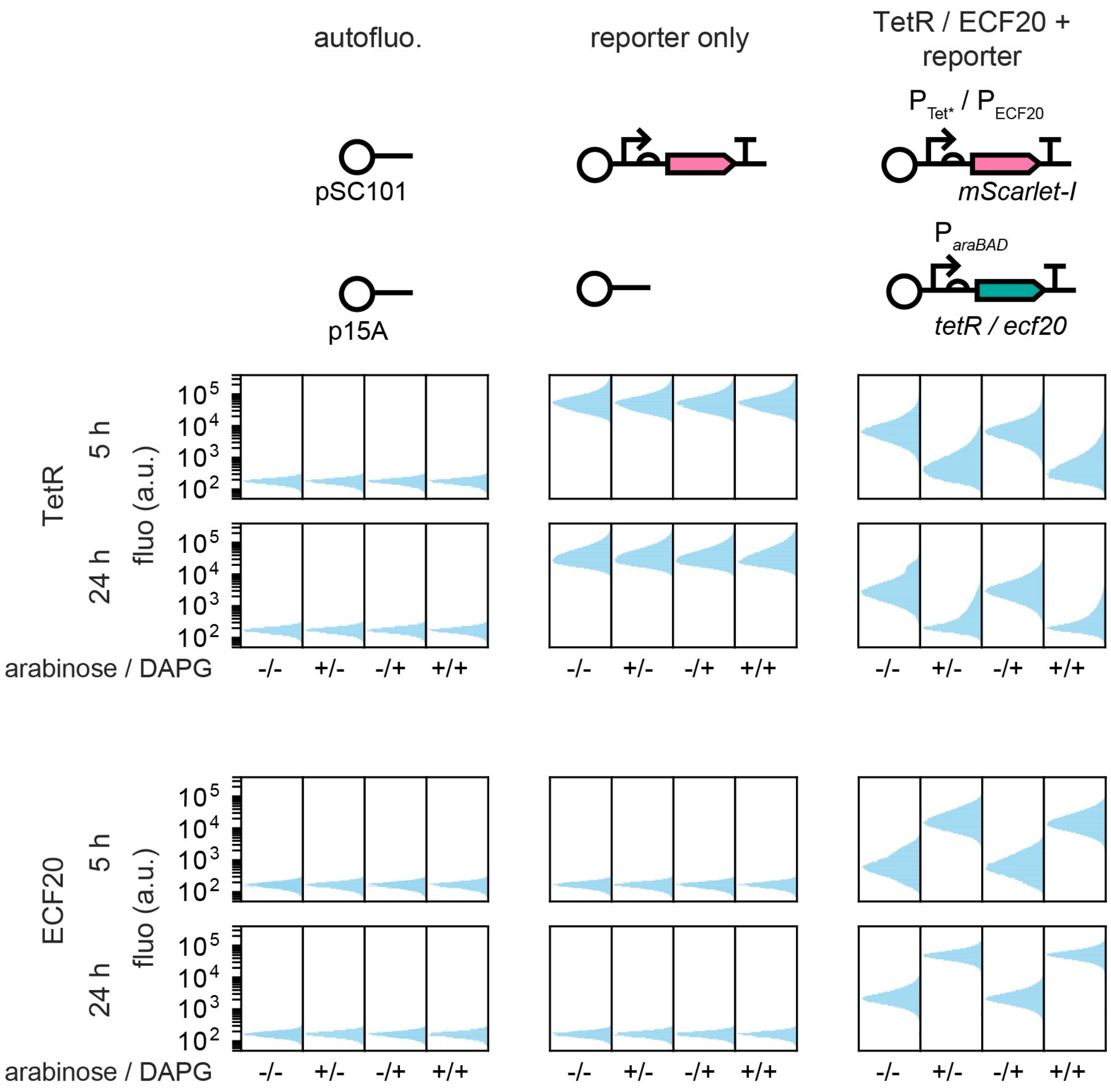

**Supplementary Figure 9 Explanation of controls of in intein-bisection maps using TetR and ECF20.** In the left subplots of TetR and ECF20 intein-bisection maps, there are three sets of vertical dots that were "autofluo.", "reporter only" and "TetR/ECF20 + reporter". They are, respectively, cells co-transformed with empty plasmids, cells co-transformed with the reporter plasmid and an empty plasmid, and cells co-transformed with the reporter plasmid and the intact protein expression plasmid. Their fluorescence profiles, under different combinations of inducers, are shown underneath the schematics. Single-cell fluorescence distributions shown were pooled from experiments performed on three different days. Addition of inducers had no influence on the basal activity or unrepressed expression of the reporter. Expression of the intact protein was under the control of $P_{araBAD}$ and thus repression or activation activities, in theory, should only be observed when arabinose was added. Results showed that induction by arabinose led to clear repression (TetR) and stronger activation (ECF20). Yet, in absence of any induction, the presence of intact TetR and ECF20 already caused some levels of repression and activation, respectively, at 5 h post-induction, and the effects were stronger at 24 h post-induction.

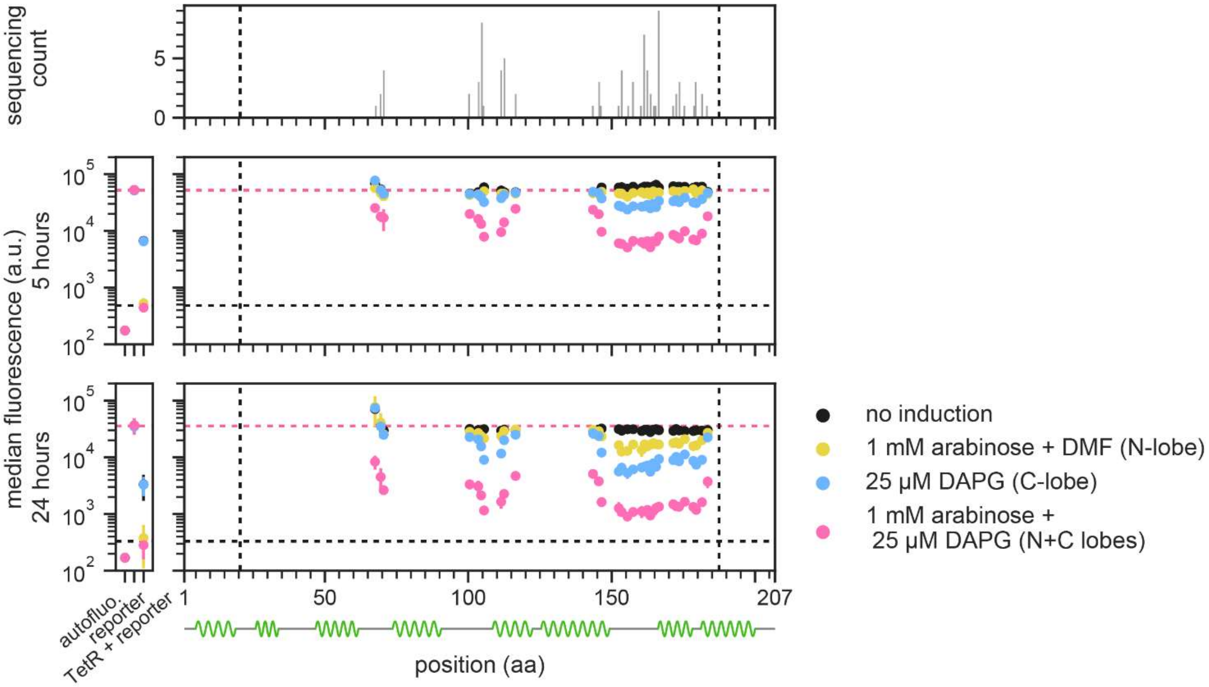

**Supplementary Figure 10 Full intein-bisection map of TetR.** Full map of IBM on TetR in which data for Figure 2b was drawn from. Left panel, the fluorescence of the controls that provide the references (horizontal dashed lines) for the activities of the intact TetR and hence the maximum repression activity in theory achievable by bipartite TetR. Right panel, bisection map of TetR. Each vertical group of spots represents an identified split site on the x axis, aligned to the TetR secondary structure (PDB: 4AC0) below. A total of 85 filtered candidate strains were characterized and sequenced to generate this map. y locations and error bars are mean and std of median fluorescence from independent experiments performed on three different days. Vertical dashed lines bound the permitted transposition window.

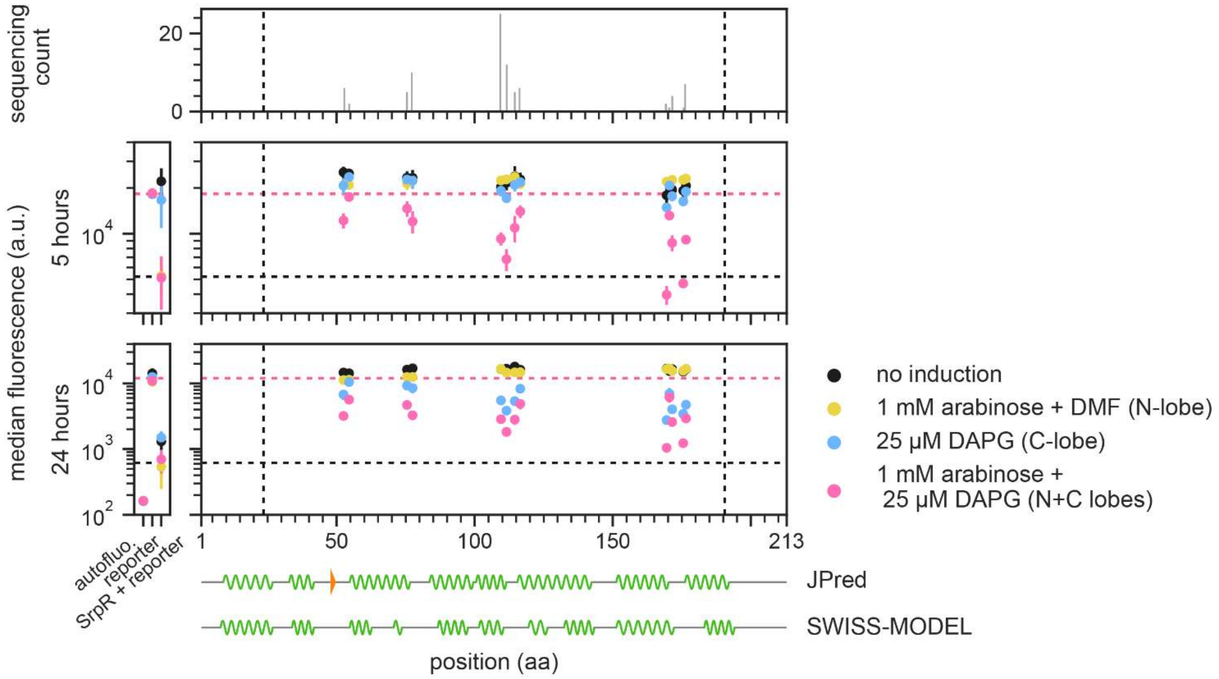

**Supplementary Figure 11 Full intein-bisection map of SrpR.** Full map of IBM on SrpR in which data for Figure 2c was drawn from. Left panel, the fluorescence of the controls that provide the references (horizontal dashed lines) for the activities of the intact SrpR and hence the maximum repression activity in theory achievable by bipartite SrpR. Right panel, bisection map of SrpR. Each vertical group of spots represents an identified split site on the x axis, aligned to two predicted SrpR secondary structures below. A total of 86 filtered candidate strains were characterized and sequenced to generate this map. JPred, the de novo secondary structure predicted from the primary sequence using the JPred 4 server[13]. SWISS-MODEL, the secondary structure predicted through the SWISS-MODEL homology modeling pipeline[14], based on PDB: 3BCG.1.A[15]. y locations and error bars are mean and std of median fluorescence from independent experiments performed on three different days. Vertical dashed lines bound the permitted transposition window.

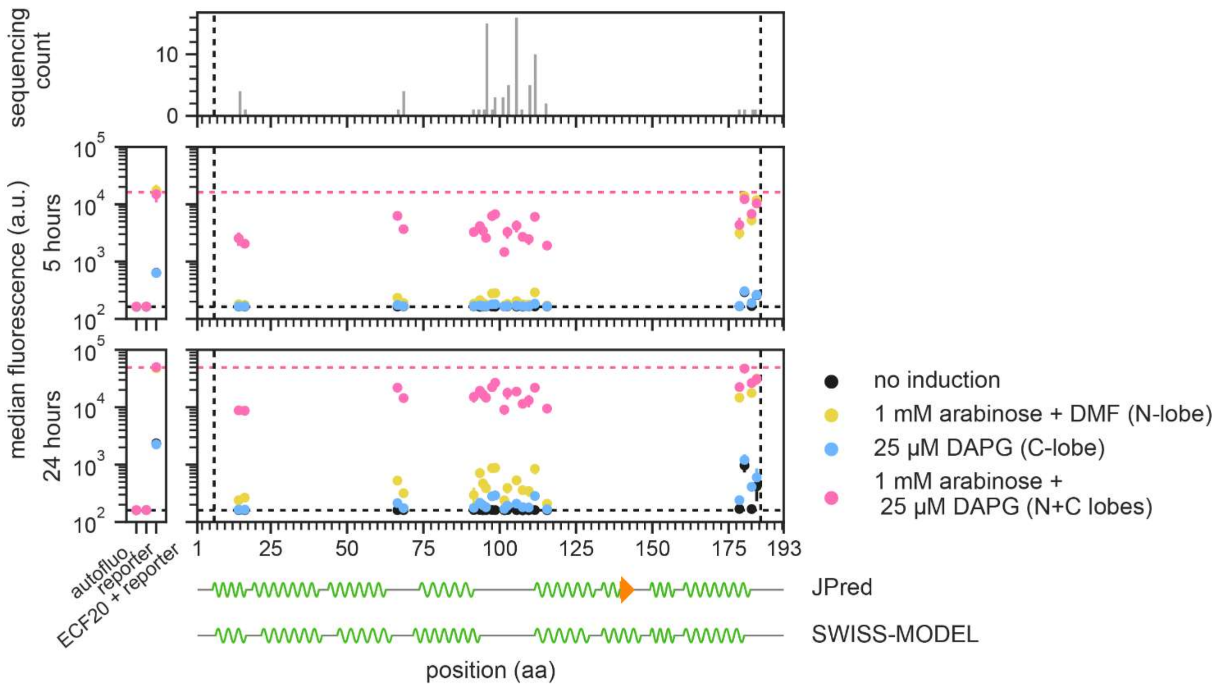

**Supplementary Figure 12 Full intein-bisection map of ECF20.** Full map of IBM on ECF20 in which data for Figure 2d was drawn from. Left panel, the fluorescence of the controls that provide the references (horizontal dashed lines) for activities of the intact ECF20 and hence the maximum activation activity in theory achievable by bipartite ECF20. Right pane, bisection map of ECF20. Each vertical group of spots represents an identified split site on the x axis, aligned to two predicted ECF20 secondary structures below. A total of 78 filtered candidate strains were characterized and sequenced to generate this map, of which 74 displayed AND logic and 4 were truncations. JPred, the de novo secondary structure predicted from the primary sequence using the JPred 4 server[13]. SWISS-MODEL, the secondary structure predicted through the SWISS-MODEL homology modeling pipeline[14], based on PDB: 6JBQ.1.F[16]. y locations and error bars are mean and std of median fluorescence from independent experiments performed on three different days. Vertical dashed lines bound the permitted transposition window.

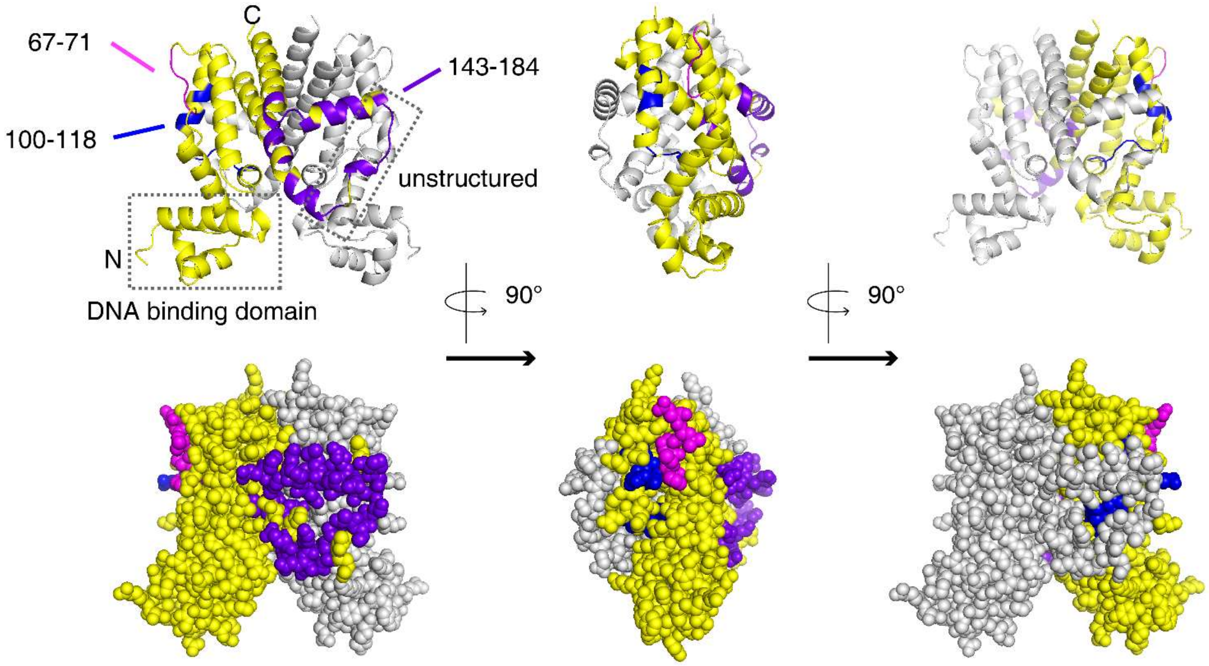

**Supplementary Figure 13 IBM-identified TetR split sites on TetR crystal structure.** Split sites identified from IBM on TetR using the M86 intein was mapped back to the TetR crystal structure (PDB: 4AC0) for illustration. Top panel: structure in cartoon. Bottom panel, structure in spheres. The crystal structure shows a TetR dimer, and each split site has the -1 and the +1 amino acid residues colored on one monomer (yellow, chain A) only. While the other monomer was shown in grey. Split sites from 67-71 were colored in magenta, 100-118, in blue, 143-184, in purple. The DNA binding domain for TetR consists of the first three alpha helices at the N-terminus. Note that the PDB model 4AC0 had missing residues that are likely unstructured regions. The model shown had those residues filled in by “predicting” the entire structure through the SWISS-MODEL homology modeling pipeline[14].

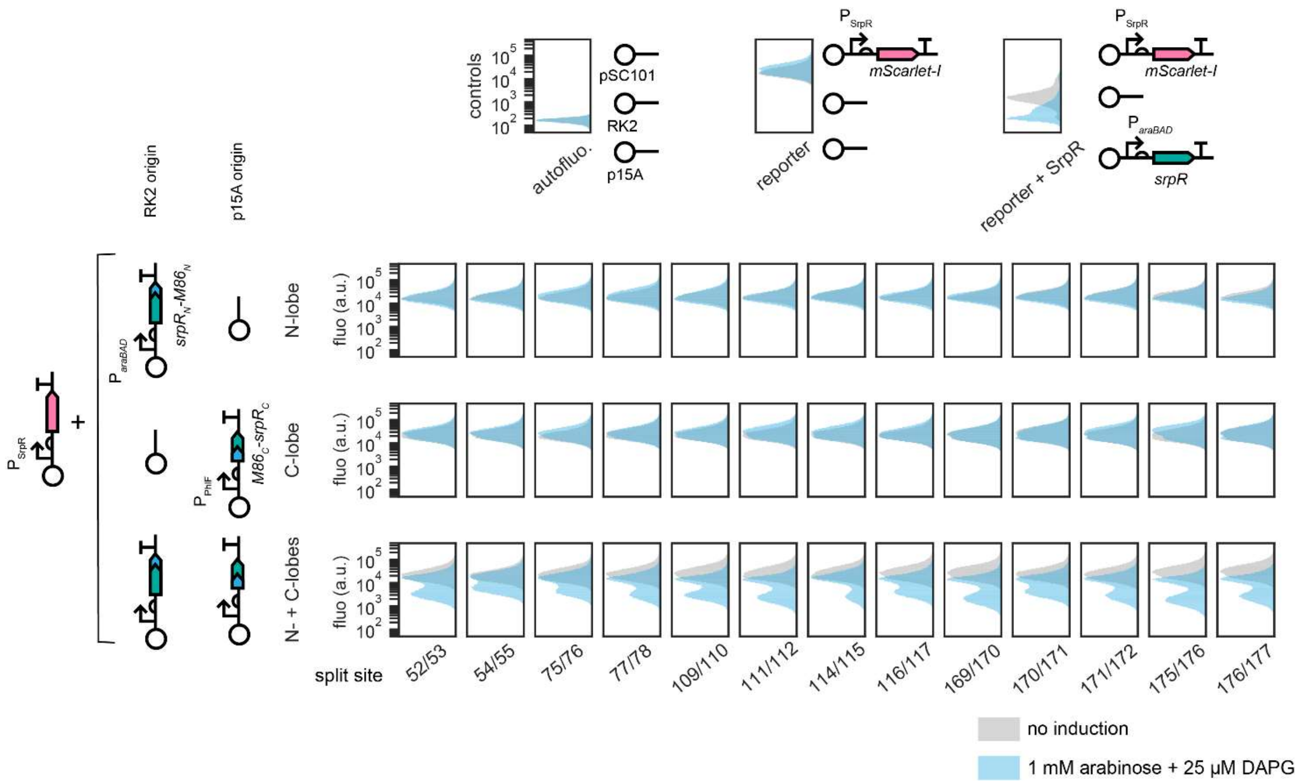

**Supplementary Figure 14 Split SrpR N- or C-lobes alone did not achieve repression.** In **Supplementary Fig. 11**, at 24 h post-induction, split SrpR at most split sites showed strong repression when C-lobes were induced alone. We hypothesized that was due to leaky expression led to N-lobes accumulation over time. To test this, the operons encoding N- and C-lobes were separated from one plasmid and subcloned into two different plasmids. Presence or absence of either lobe was achieved co-transforming a construct carrying plasmid or an empty plasmid, respectively, with the reporter plasmid and the expression plasmid of the other lobe into *E. coli*. Shown in the figure are single cell fluorescence data for the combinatorial transformations, which were performed for all split sites and controls. Results showed that at all split sites, both the N- and C-lobes were required for repression to happen. This proved that neither lobe alone was sufficient for repression and all split SrpR fused with split M86 intein were authentic NAND gates. Single-cell fluorescence distributions shown were pooled from experiments performed on three different days and cells were induced for 24 h prior to fluorescence assay.

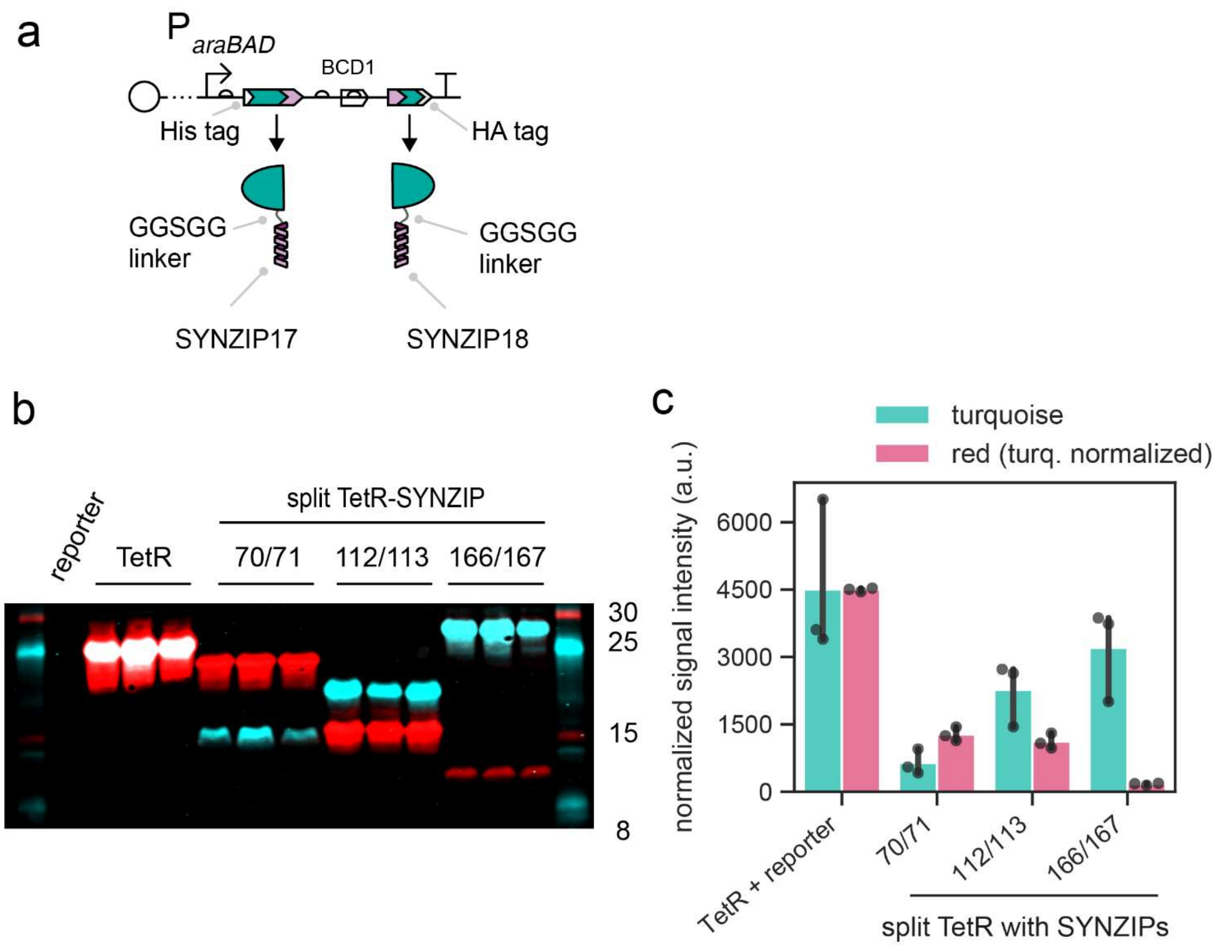

**Supplementary Figure 15 Quantification of Split TetR-SYNZIP at three split sites. a.** Construct schematics for data shown in **Fig. 3a**. The N-lobe contains a hexahistidine at the N-terminus and the C-lobe, a HA tag at the C-terminus. This allowed the abundances of the protein fragments to be measured semi-quantitatively from Western blots. The intact TetR construct also carries the two tags. BCD, bicistronic design to increase the likelihood that the C-lobes would be expressed at similar levels despite having different 5' coding DNA sequences[9]. **b.** Western blot on which quantification was performed. Each whole-cell lysate from one of the three biological replicates, from the same experiment where fluorescence measurements were taken, was loaded on one lane for quantification of protein abundances. N- and C-lobes expression were probed using antibodies that target the hexahistidine tag (turquoise) and the HA tag (red) respectively. The lane "reporter" was a whole-cell lysate from bacteria carrying the reporter and empty plasmid, and demonstrates endogenous proteins would not cross-react with either antibody. The reporter plasmid was present in all other whole-cell lysates as well. **c.** Quantification result from b. The turquoise and red signals were normalized using TetR since they must be in 1:1 ratio. Across the three split constructs, split TetR-SYNZIP at site 166/167 had the lowest expression on one of the lobes (C-lobe). Since the N- and C-lobes must reconstitute in a 1:1 ratio, split TetR-SYNZIP at site 166/167 should have the least abundance of reconstituted TetR. However, it still outperformed the other two split sites in the fluorescence repression assay. This result rejected the possibility that differences in repression strengths were due to inadequate protein expression at sites 70/71 and 112/113.

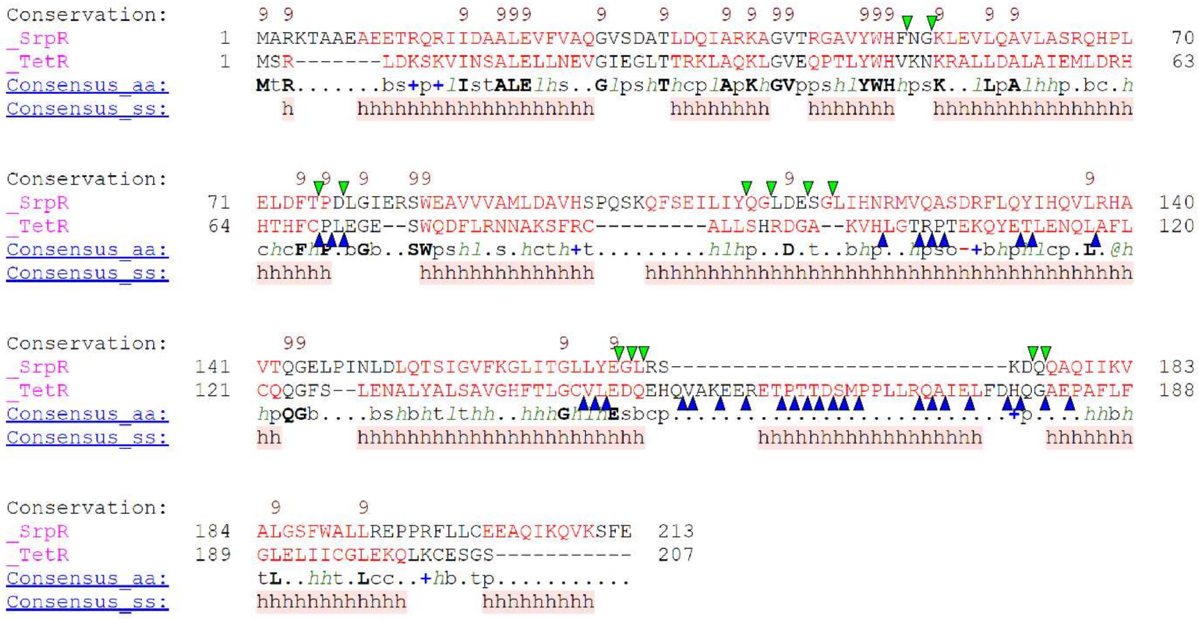

**Supplementary Figure 16 Secondary structure alignment of TetR and SrpR with identified split sites revealed limitation of inferring split sites from homology alignment.** SrpR and TetR amino acid sequences were aligned with the known TetR structure (PDB: 4AC0) using PROMALS3D with default parameters[17]. IBM identified split sites for SrpR and TetR were marked by green and blue triangles, respectively. The result was re-rendered into Figure 3b. As shown, only the second and the last split seams could be mutually inferred.

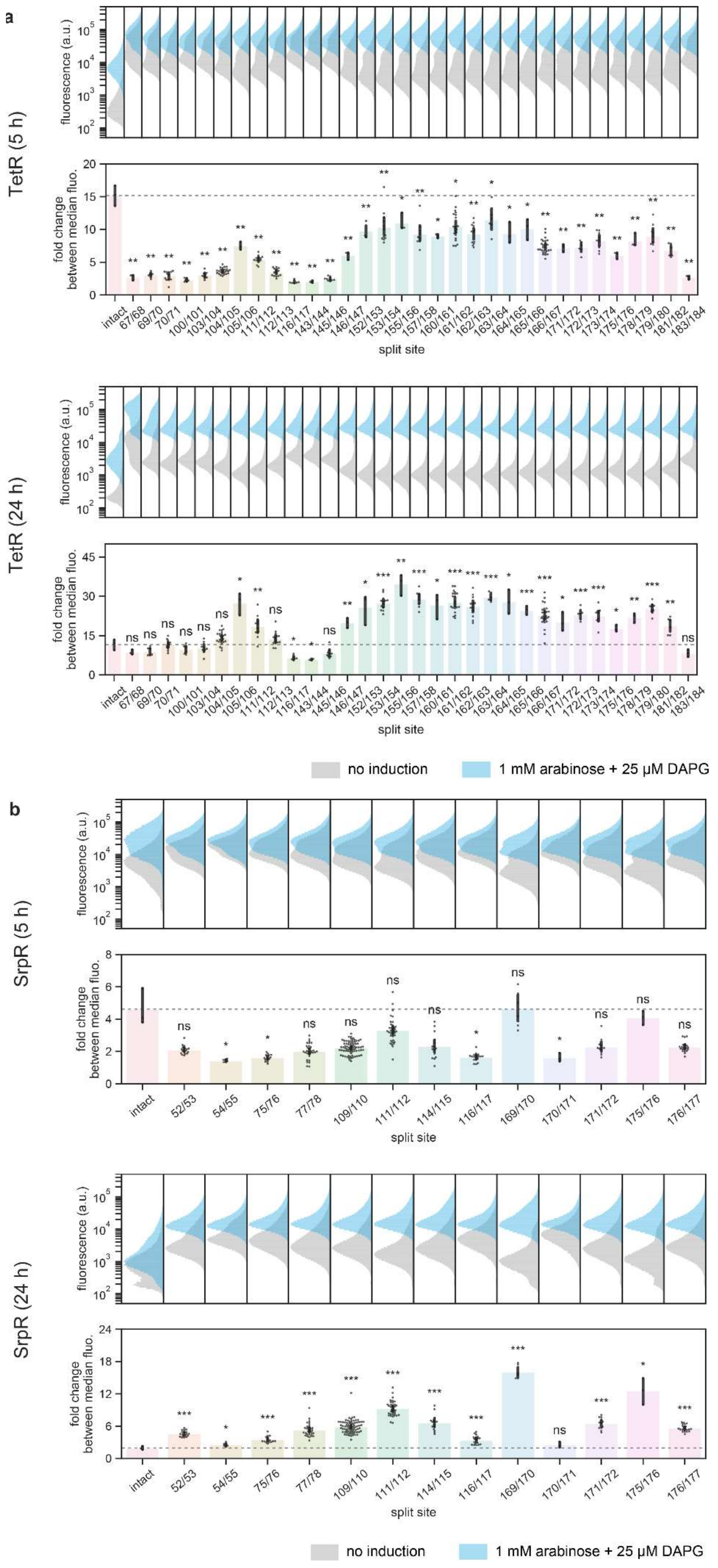
a
TetR (5 h)
fluorescence (a.u.)
fold change between median fluo.
split site
intact
67/68
69/70
70/71
100/101
103/104
104/105
105/106
111/112
112/113
116/117
143/144
145/146
146/147
152/153
153/154
155/156
157/158
160/161
161/162
162/163
163/164
164/165
165/166
166/167
171/172
172/173
173/174
175/176
178/179
179/180
181/182
183/184
TetR (24 h)
no induction
1 mM arabinose + 25 µM DAPG
b
SrpR (5 h)
SrpR (24 h)
intact
52/53
54/55
75/76
77/78
109/110
111/112
114/115
116/117
169/170
170/171
171/172
175/176
176/177
ns
no induction
1 mM arabinose + 25 µM DAPG

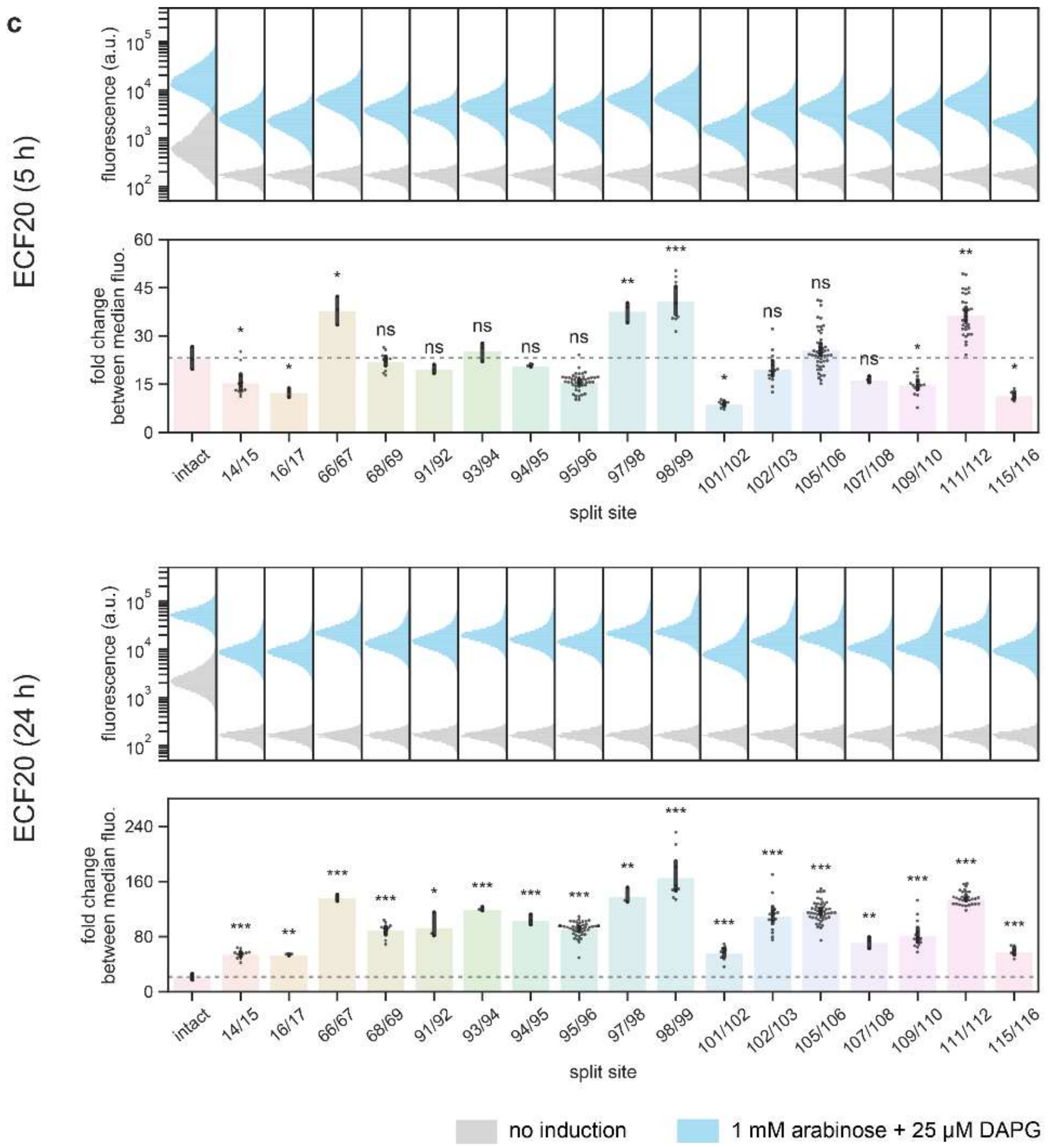

**Supplementary Figure 17 Bipartite proteins at different split sites have lower overall basal activities over prolonged growth.** For TetR (**a**), SrpR (**b**) and ECF20 (**c**), within each subpanel for 5 or 24 h post-induction, the upper panel are pooled single-cell fluorescence data from three independent experiments performed on 3 different days, with no induction, or expression of the intact protein, or both N- and C-lobes. The lower panels are fold changes obtained by dividing, for each strain and each experiment, the median fluorescence value of the high fluorescent population by that of the low fluorescent population. The horizontal dashed lines serve as visual guides for fold changes of the intact protein. For prolonged growth (24 h), most bipartite proteins had less leaky activities than their intact counterparts and hence greater fold changes. Data are from the same source data that was used to produce the intein-bisection maps in Supplementary Figures 6-8, and representative data for each protein was repeated in Figure 3c. For each fold change panel, statistics summaries above each bar is a two-tailed t-test, assuming unequal variance, that compares the fold changes between the intact protein (n = 3) and that of the bipartite construct split at different positions (various n, n ≥ 3).

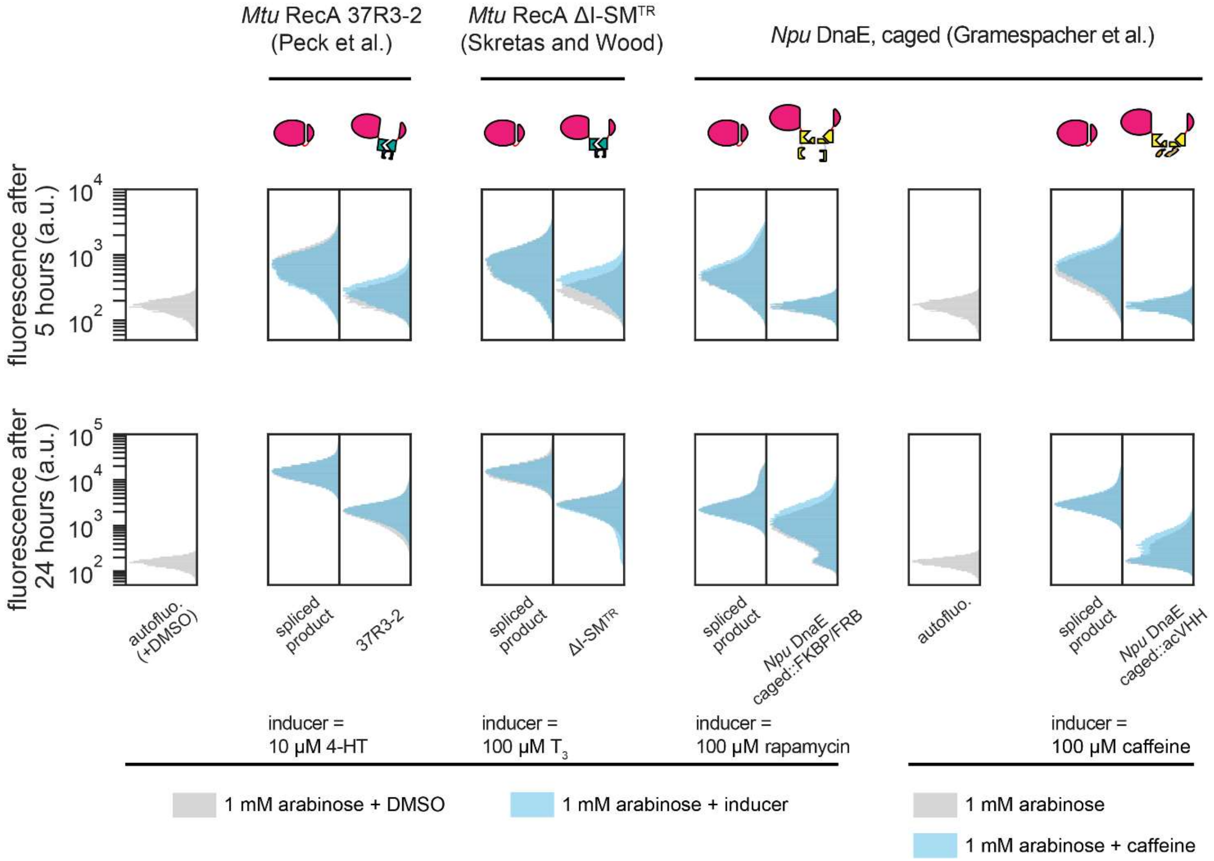

**Supplementary Figure 18 Testing literature reported chemically inducible inteins.** Three switchable inteins[18-20] were picked from the literature, synthesized with the known extein junctions, and were inserted into or used to split the mCherry at site 192/193. The resulting constructs were then assayed with or without the corresponding inducer alongside a control that simulated the spliced product with linker due to extein junctions. The result showed that different extein junctions were tolerated at the split site. Under this context, all assayed conditional inteins did not give sufficient differential responses to substantiate further bisection or insertion mapping. Single-cell fluorescence distributions shown were pooled from three biological replicates. It should be noted that, the intein *MtuA* RecA 37R3-2 was not reported to work in *E. coli* but since it also used the ER-LBD (albeit mutated) and ER-LBD should function in bacteria[21], it was included for testing.

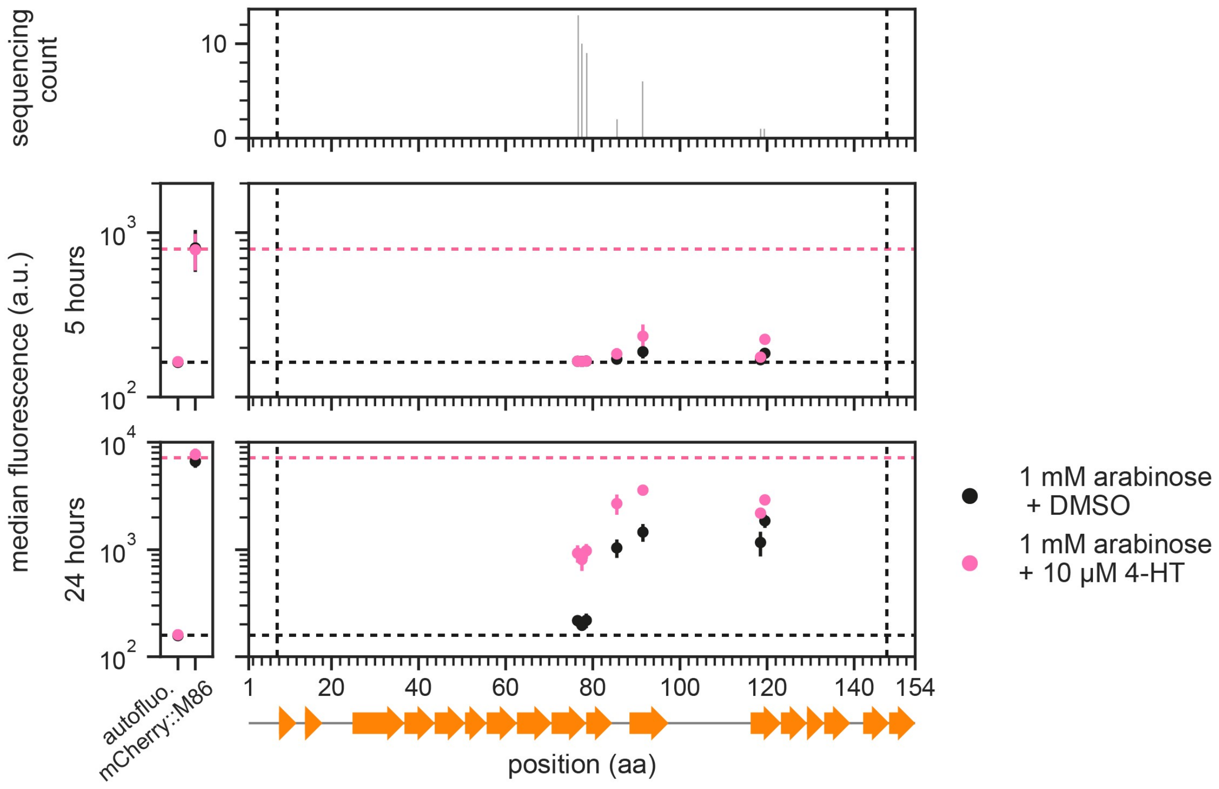

**Supplementary Figure 19 Full ER-LBD-insertion map of the M86 intein inserted in mCherry spilt at 192/193.** Full domain-insertion map for ER-LBD inserted into mCherry(1-192)-G-M86-S-mCherry(193-236) used to produced Figure 4b. Left panel, the fluorescence of the controls that provide the references (horizontal dashed lines) for the activities of the cis-M86 intein-inserted mCherry and hence the maximum fluorescence in theory achievable. Right panel, domain-insertion map. Each vertical group of spots represents an identified insertion site on the x axis, aligned to the secondary structure of the M86 intein below (PDB: 6FRH)[22]. A total of 42 filtered candidate strains were characterized and sequenced to generate this map. y locations and error bars are mean and std of median fluorescence from independent experiments performed on three different days. Vertical dashed lines bound the permitted transposition window.

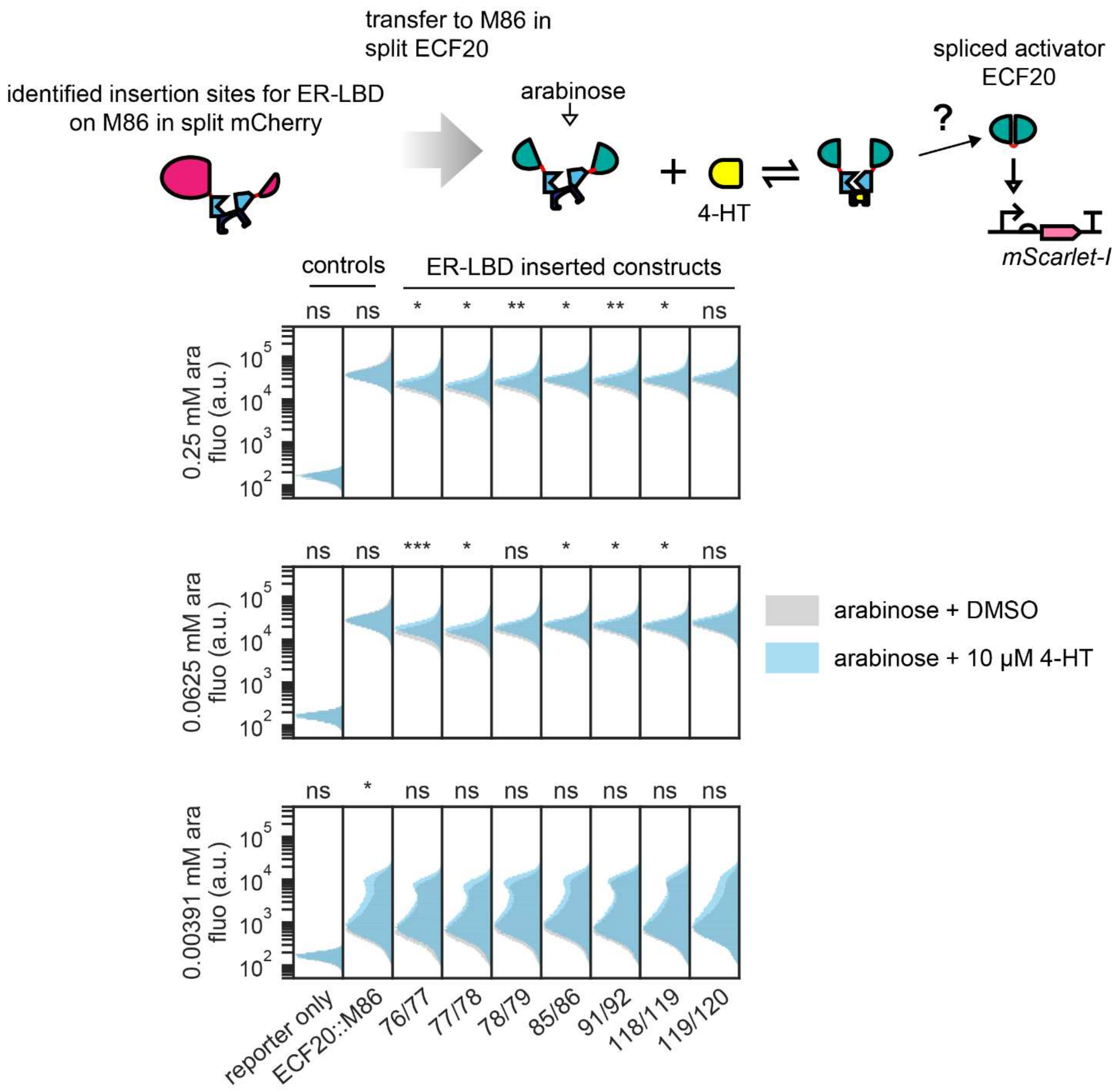

**Supplementary Figure 20 ER-LBD-inserted M86 intein from split mCherry was not transferrable to split ECF20.** All seven conditional ER-LBD-inserted M86 inteins were subcloned from the domain-inserted strains from their split mCherry contexts (site 192/193) into split ECF20 (site 101/102). Protein expression was induced by different concentrations of arabinose for 24 h in the presence or absence of 10 µM 4-HT. Results showed little to no shift in fluorescence population upon 4-HT induction. At 0.0625 mM and 0.25 mM arabinose, some constructs displayed very small, albeit statistically significant differences in the centers of mass of fluorescence populations. Histograms show single-cell fluorescence pooled from four biological replicates. Above each histogram, the statistics summary is a two-tailed t-test, assuming unequal variance, that compares the median fluorescence values between the uninduced and induced populations (n = 4).

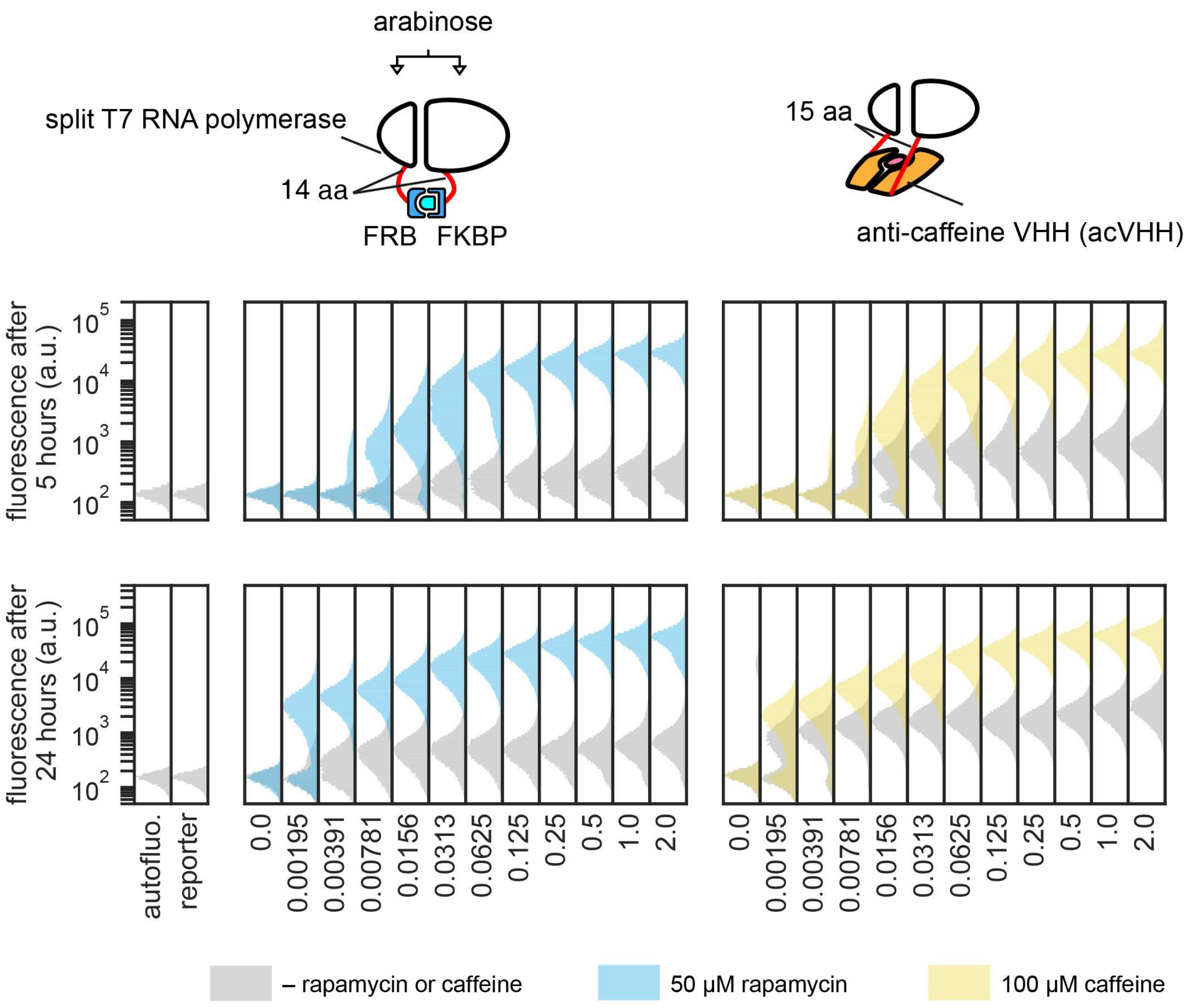

**Supplementary Figure 21 Chemically inducible dimerization by caffeine binding acVHH is similar to that by FRB/FKBP.** The evolved split T7 RNA polymerase[23] was used as a test platform to compare the efficacies and performance of small molecule-induced dimerization between anti-caffeine VHH[24] and FRB/FKBP domains. Strains carrying the constructs were induced under various arabinose concentrations with or without rapamycin (for FRB/FKBP) or caffeine (acVHH) for 5 h. Maximal achievable activation activities were similar between the different domains. In both cases, higher expression levels of the bipartite parts led to higher basal activities, likely due to an increase in local protein concentrations that increased spontaneous reconstitution of T7 RNAP by random collision. However, at lower protein expression strengths T7 RNA polymerase split by FRB/FKBP had lower tendencies to self-associate and gave less basal activities. Linker lengths for FRB/FKBP was chosen from the constructs that displayed strongest activation from Pu et al.[24], and linker lengths for acVHH were chosen to be made comparable to those of FRB/FKBP. It should be noted, however, based on the tertiary structure of acVHH[25], the linker lengths were reduced to 10 residues each in subsequent experiments. Single-cell fluorescence distributions shown were pooled from three independent experiments performed on different days.

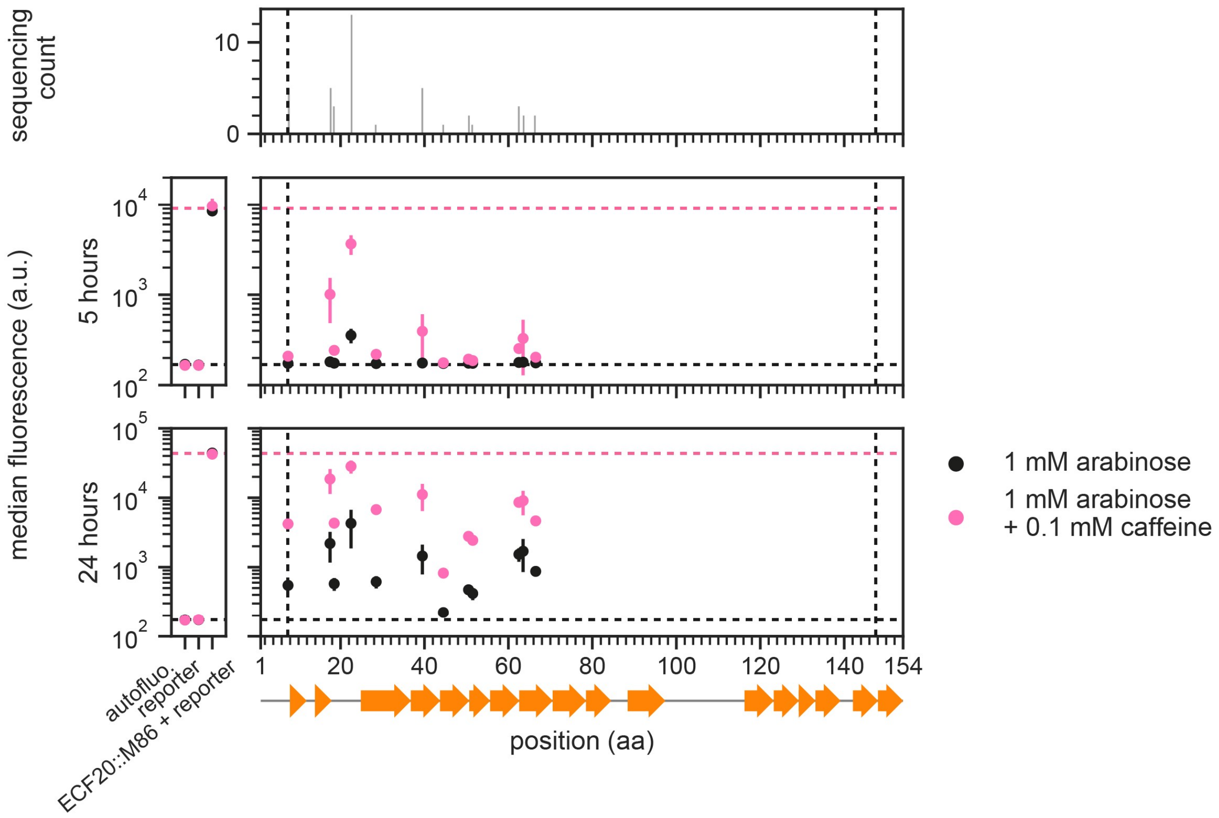

**Supplementary Figure 22 Full acVHH-assisted bisection map of the M86 intein inserted in mCherry spilt at ECF20 101/102.** Full bisection map for acVHH-inserted split sites within ECF20(1-101)-G-M86-S-ECF20(102-193) used to produced Figure 4d. Left panel, the fluorescence of the controls that provide the references (horizontal dashed lines) for the activities of cis-M86 intein-inserted ECF20 and hence the maximum activation activity in theory achievable. Right pane, acVHH-bisection map. aligned to the secondary structure of the M86 intein below (PDB: 6FRH)[22]. A total of 43 filtered candidate strains were characterized and sequenced to generate this map. y locations and error bars are mean and std of median fluorescence from independent experiments performed on three different days. Vertical dashed lines bound the permitted transposition window.

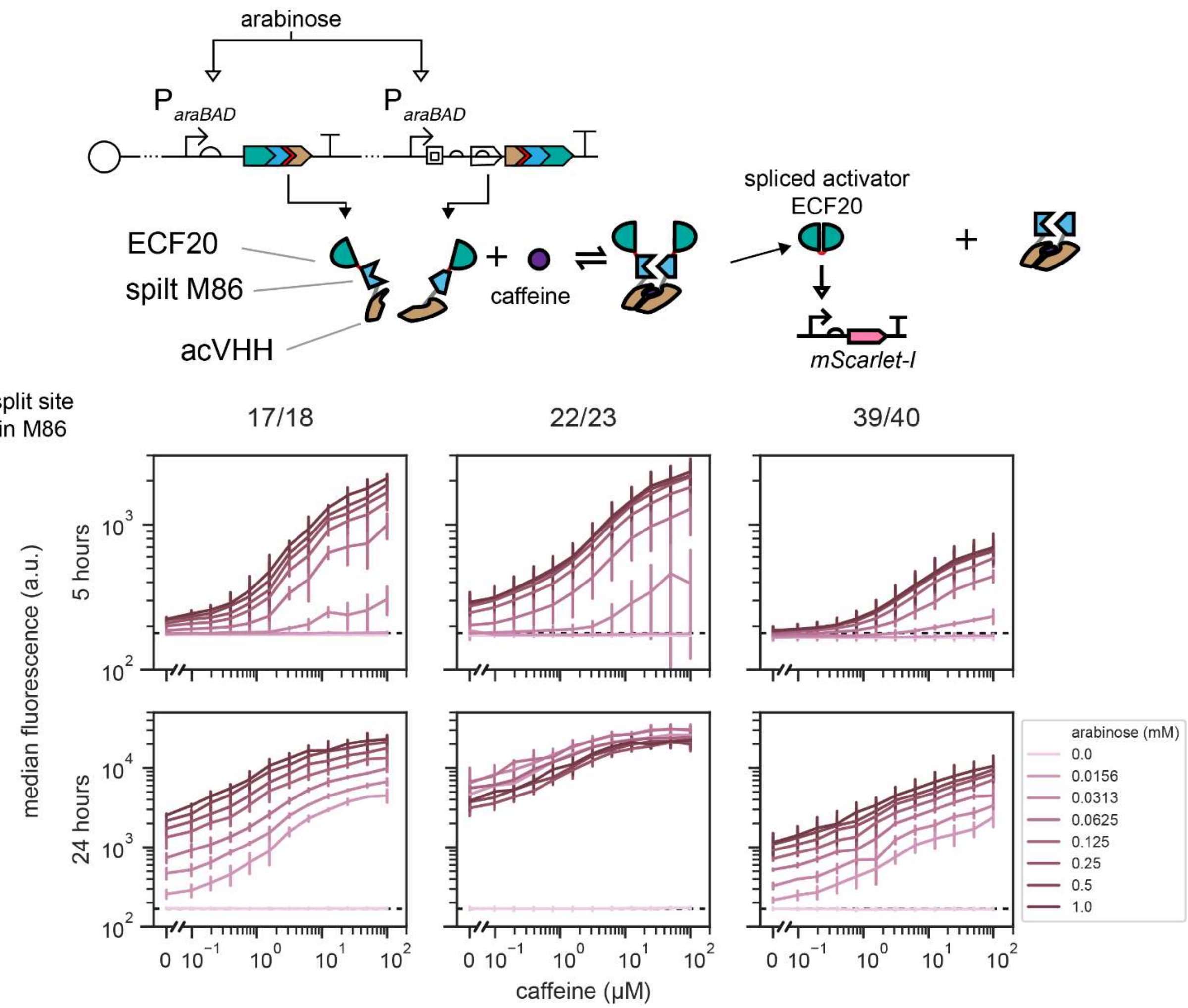

**Supplementary Figure 23 Dose-dependent responses of selected acVHH-bisected M86 intein within split ECF20.** All response curves of representative acVHH-inserted ECF20(1-101)-M86-ECF20(102-193) clones subjected to a 2D gradient induction of caffeine and arabinose. Selected curves were displayed in Figure 4e. Results showed that activation responses gradually increased with caffeine concentrations and magnitudes were dependent on protein expression strengths. There were 11 caffeine concentrations, diluted 2-fold starting from 100 μM plus no caffeine. y locations and error bars are mean and std of median fluorescence from experiments performed on 3 different days. Horizontal dashed lines marked the mean of median autofluorescence.

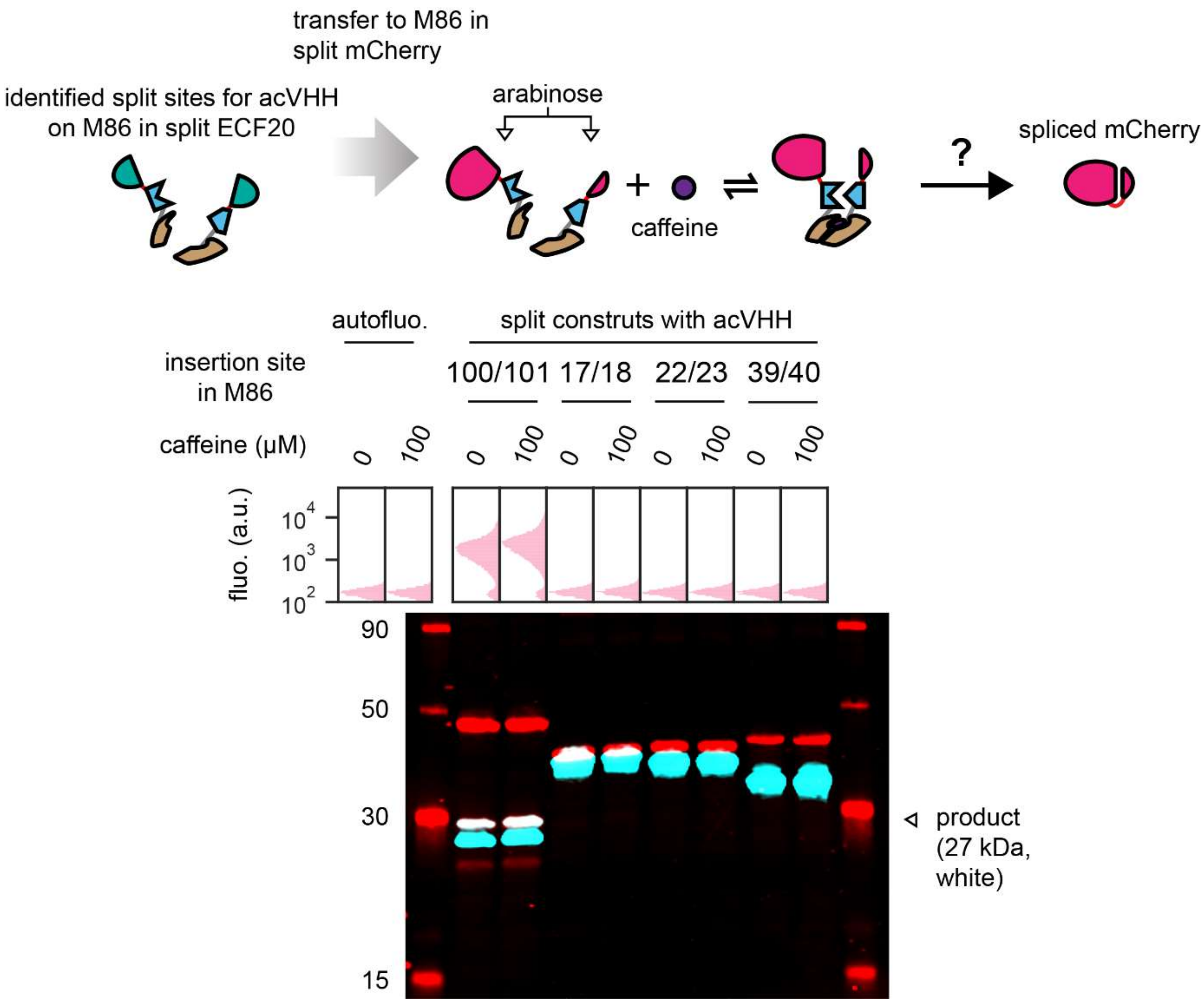

**Supplementary Figure 24 acVHH-bisected M86 intein from split ECF20 was not transferrable to split mCherry.** The three representative caffeine inducible acVHH-bisected M86 intein from Supplementary Figure 16 (sites 17/18, 22/23, 39/40) were subcloned from their split ECF20 (site 101/102) contexts into split mCherry (site 192/193). To provide a control, the M86 intein was split at the conventional split site that permits spontaneous re-assembly (site 100/101). In all constructs, mCherry carried a hexahistidine tag at the C-terminus. Cells were induced for protein expression at 1 mM arabinose for 24 h in the presence and absence of 100 µM caffeine. Top panel, single-cell fluorescence of induced or uninduced cells prior to cell lysis for Western blot. Fluorescence could be detected for the construct where acVHH was inserted at the split site within the M86 intein for spontaneous re-assembly but not in other cases. Each distribution comes from one biological sample. Bottom panel, the Western blot result for whole-cell lysates on the bipartite constructs after fluorescence was analyzed. N- and C-lobes expression were probed using antibodies that target the mCherry epitope (red) and the hexahistidine tag (turquoise) respectively. In all lanes both the N- and C-lobes could be detected, but spliced product formation (overlapped bands in white) could be observed at site 100/101 but not the rest, indicating that the tested caffeine inducible inteins were non-functional when moved out of their contexts in which they were screened out.

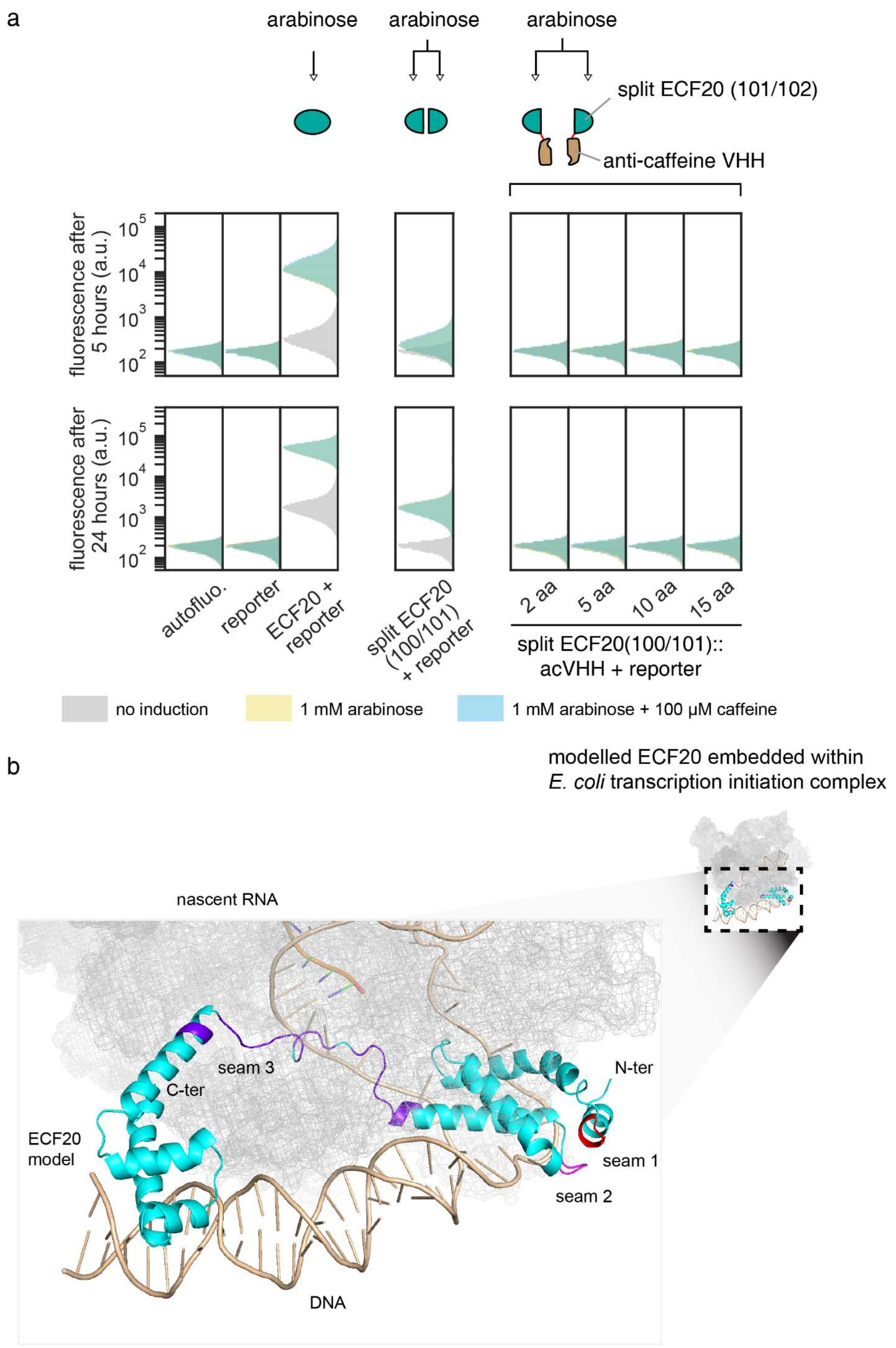
a
arabinose
arabinose
arabinose
split ECF20 (101/102)
anti-caffeine VHH
fluorescence after 5 hours (a.u.)
fluorescence after 24 hours (a.u.)
autofluo.
reporter
ECF20 + reporter
split ECF20 (100/101) + reporter
2 aa
5 aa
10 aa
15 aa
split ECF20(100/101):: acVHH + reporter
no induction
1 mM arabinose
1 mM arabinose + 100 µM caffeine
b
modelled ECF20 embedded within E. coli transcription initiation complex
nascent RNA
seam 3
C-ter
N-ter
ECF20 model
seam 1
seam 2
DNA

**Supplementary Figure 25 ECF20 as an example where additional domains are not tolerated at a split site.** **a.** ECF20 was split at site 101/102 and fused to acVHH domains using various linker lengths between the acVHH and the spilt ECF20 parts, but none was functional. However, expression of bipartite parts alone without any fusion yielded a much weaker but still functional activation. For this experiment, cells carrying constructs were induced for 24 h to allow accumulation of any activation activities such that they could be observed. Single-cell fluorescence distributions shown were pooled from three biological replicates. Note that populations induced with arabinose, with or without caffeine appear almost identical and thus give a green appearance due to overlap of colors. **b.** The results obtained could be explained by a predicted 3D model of ECF20. During transcription initiation, the linker of a sigma factor is normally embedded within the RNA polymerase (gray mesh). Addition of acVHH domains at linker (site 101/102) thus obstructs transcription initiation complex formation. Model is generated by the SWISS-MODEL homology modeling pipeline with PDB: 6JBQ.1.F[16] as the template.

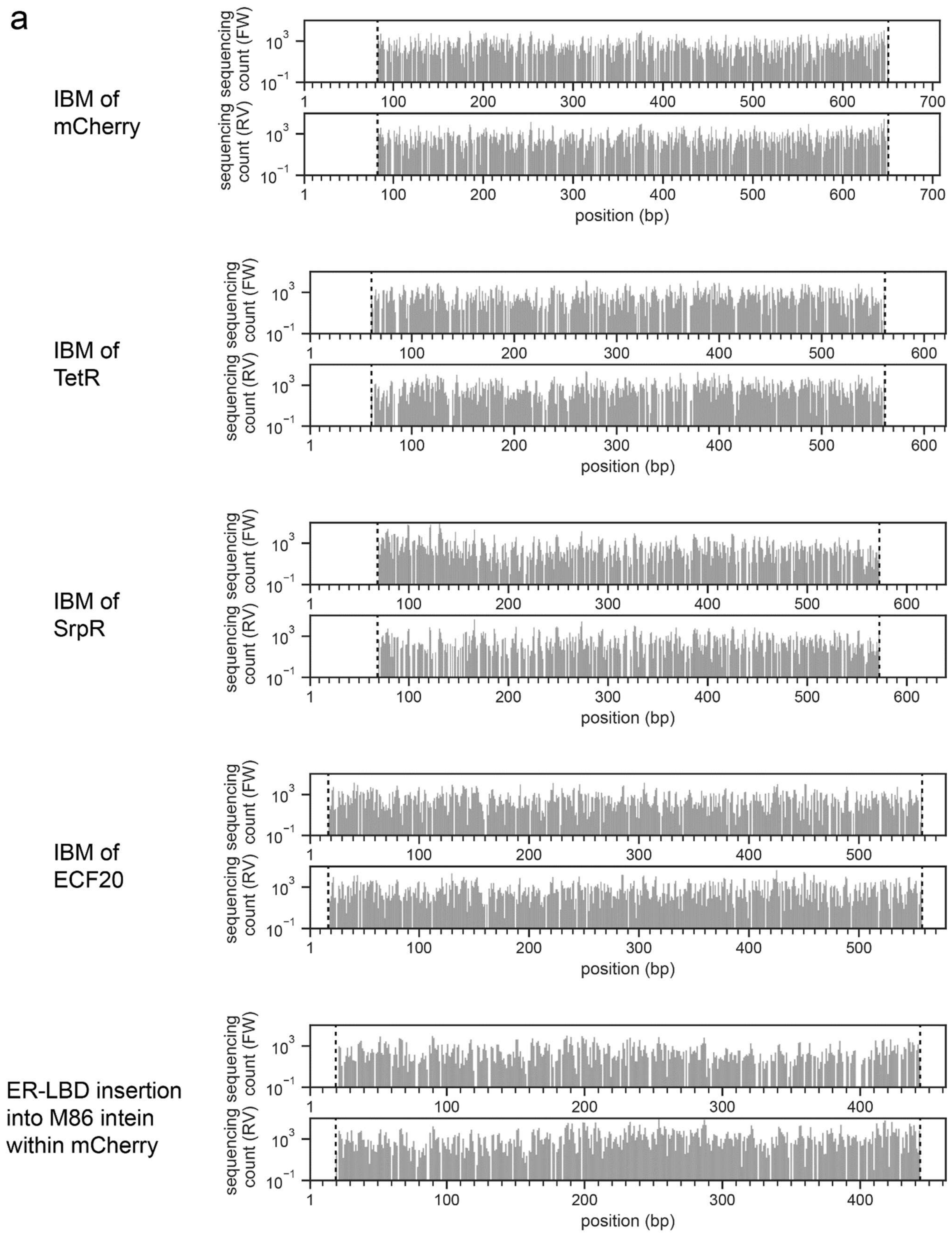
a
IBM of mCherry
IBM of TetR
IBM of SrpR
IBM of ECF20
ER-LBD insertion into M86 intein within mCherry
sequencing count (FW)
sequencing count (RV)
position (bp)

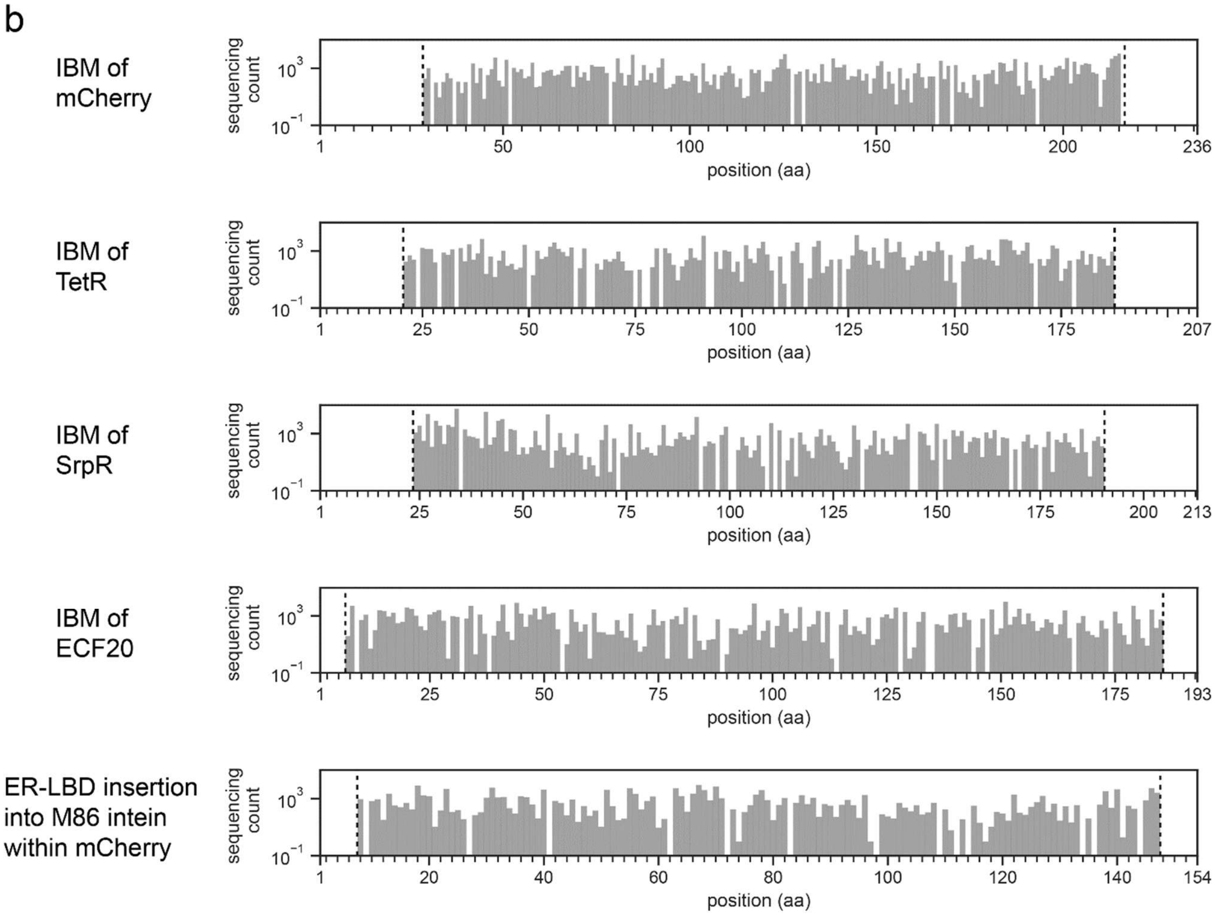

**Supplementary Figure 26 NGS confirms that most split/insertion sites are covered when using the mini-Mu transposon. a.** Histograms showing DNA coverages of five final libraries prior to screening. Gaps represent missing DNA insertion positions. As shown, random insertion by transposition could cover most DNA insertion points on both the forward and reverse orientations. Starting DNA positions are the first base pairs in the CDS of the protein. Note that we define the 5', 5 bp duplicated sequence as the non-native inserted sequence regardless of insertion direction. **b.** Histograms showing the coverages of split sites of the five final libraries prior to screening. Gaps represent missing protein split/insertion sites. NGS results showed that most split/insertions sites were covered. For ER-LBD insertion into M86 intein, the starting amino acid position refers to the first amino acid of the intein. **a, b.** Vertical lines denote transposition windows, which are inclusive.

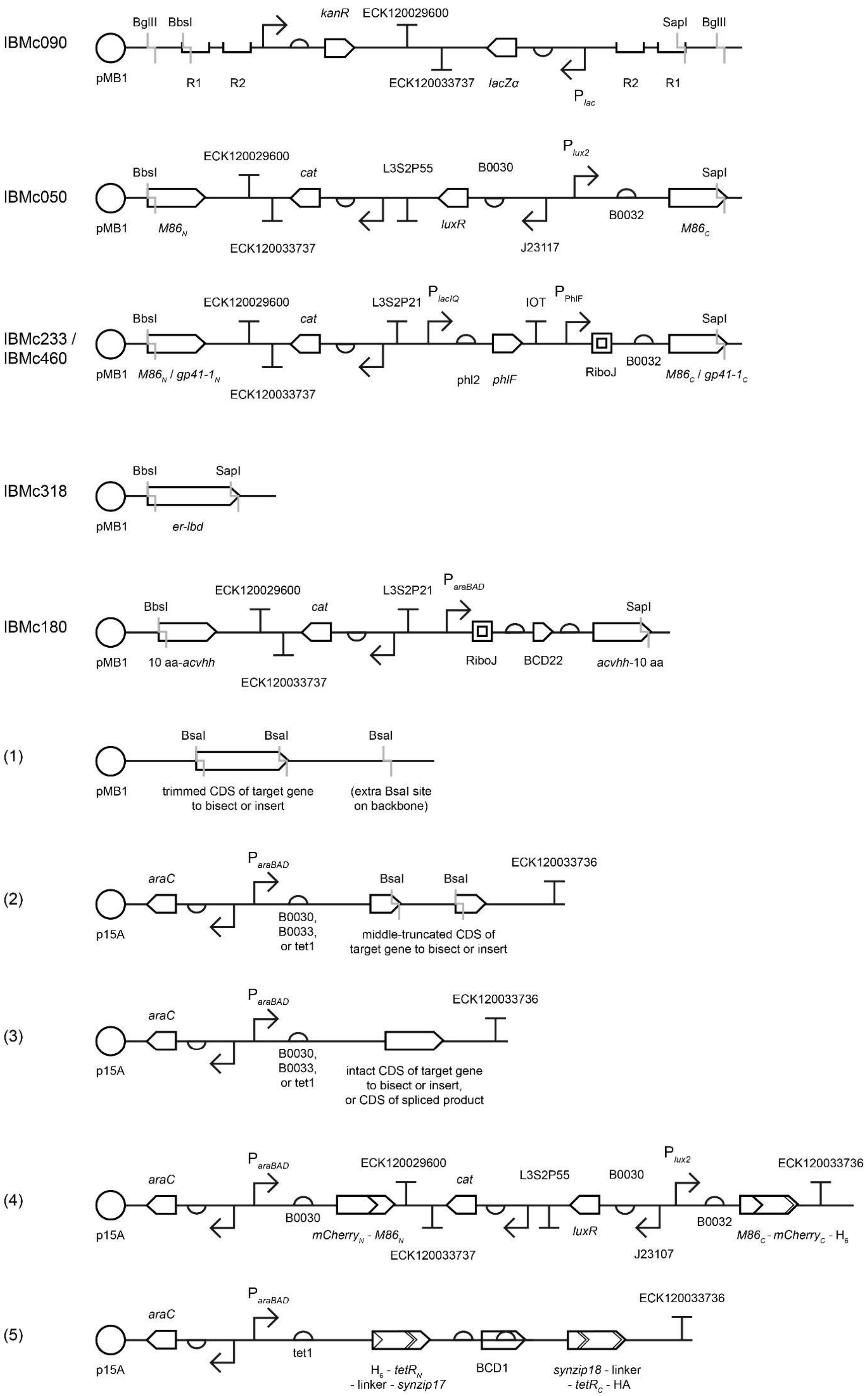
IBMc090
BglII BbsI
kanR ECK120029600
SapI BglII
pMB1 R1 R2 ECK120033737 lacZα $P_{lac}$ R2 R1
IBMc050
BbsI ECK120029600 cat L3S2P55 B0030 $P_{lux2}$ SapI
pMB1 $M86_N$ ECK120033737 luxR B0032 J23117 $M86_C$
IBMc233 / IBMc460
BbsI ECK120029600 cat L3S2P21 $P_{lacIQ}$ IOT $P_{PhlF}$ SapI
pMB1 $M86_N$ / $gp41\text{-}1_N$ ECK120033737 phl2 phlF RiboJ B0032 $M86_C$ / $gp41\text{-}1_C$
IBMc318
BbsI SapI
pMB1 er-lbd
IBMc180
BbsI ECK120029600 cat L3S2P21 $P_{araBAD}$ SapI
pMB1 10 aa-acvhh ECK120033737 RiboJ BCD22 acvhh-10 aa
(1)
BsaI BsaI BsaI
pMB1 trimmed CDS of target gene to bisect or insert (extra BsaI site on backbone)
(2)
araC $P_{araBAD}$ BsaI BsaI ECK120033736
p15A B0030, B0033, or tet1 middle-truncated CDS of target gene to bisect or insert
(3)
araC $P_{araBAD}$ ECK120033736
p15A B0030, B0033, or tet1 intact CDS of target gene to bisect or insert, or CDS of spliced product
(4)
araC $P_{araBAD}$ ECK120029600 cat L3S2P55 B0030 $P_{lux2}$ ECK120033736
p15A B0030 $mCherry_N$ - $M86_N$ ECK120033737 luxR J23107 B0032 $M86_C$ - $mCherry_C$ - $H_6$
(5)
araC $P_{araBAD}$ ECK120033736
p15A tet1 $H_6$ - $tetR_N$ - linker - synzip17 BCD1 synzip18 - linker - $tetR_C$ - HA

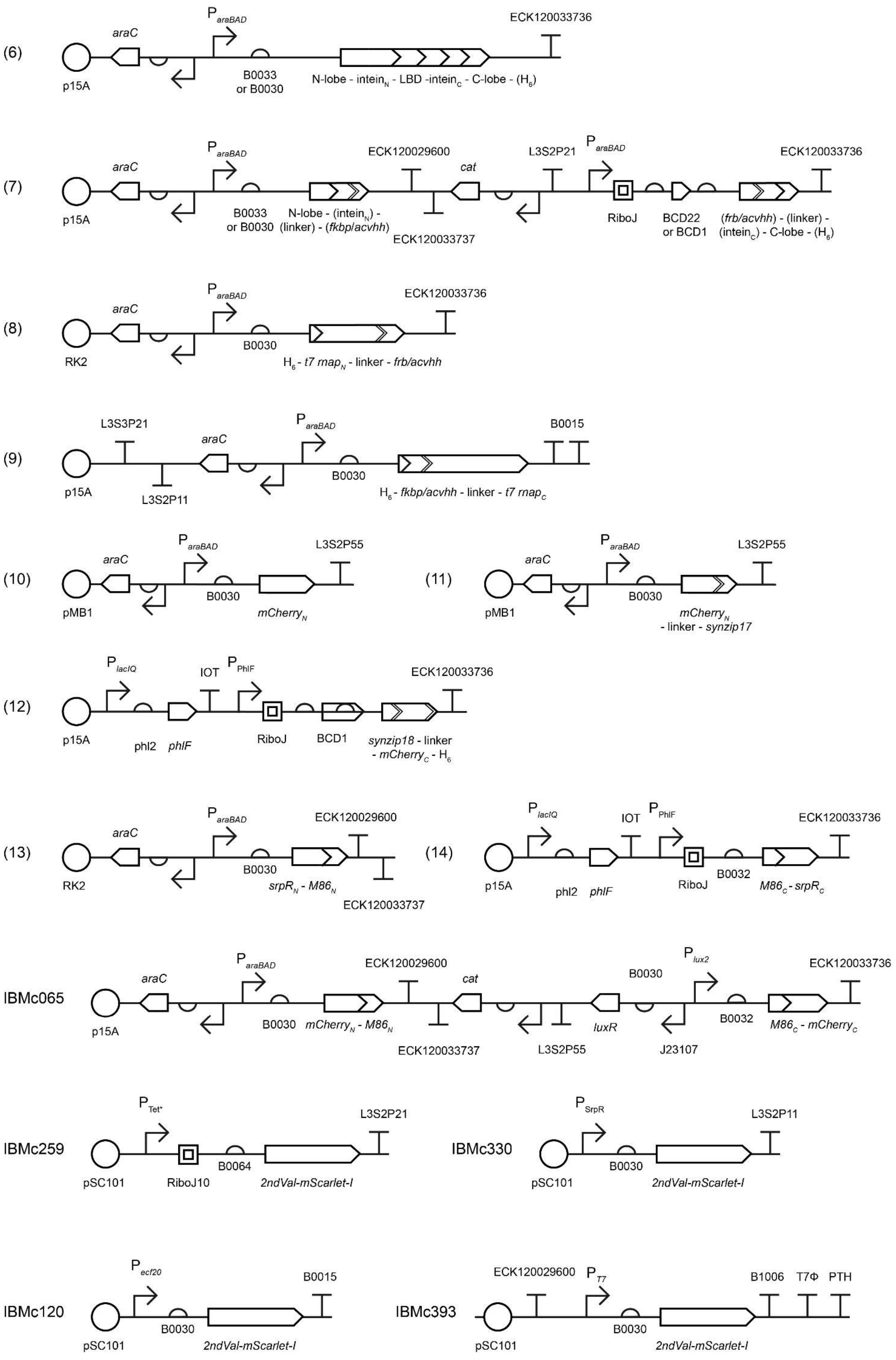
(6)
p15A
araC
ParaBAD
B0033 or B0030
N-lobe - inteinN - LBD -inteinC - C-lobe - (H6)
ECK120033736
(7)
p15A
araC
ParaBAD
B0033 or B0030
N-lobe - (inteinN) - (linker) - (fkbp/acvhh)
ECK120029600
ECK120033737
cat
L3S2P21
ParaBAD
RiboJ
BCD22 or BCD1
(frb/acvhh) - (linker) - (inteinC) - C-lobe - (H6)
ECK120033736
(8)
RK2
araC
ParaBAD
B0030
H6 - t7 rnapN - linker - frb/acvhh
ECK120033736
(9)
p15A
L3S3P21
L3S2P11
araC
ParaBAD
B0030
H6 - fkbp/acvhh - linker - t7 rnapC
B0015
(10)
pMB1
araC
ParaBAD
B0030
mCherryN
L3S2P55
(11)
pMB1
araC
ParaBAD
B0030
mCherryN - linker - synzip17
L3S2P55
(12)
p15A
PlacIQ
phl2
phlF
IOT
PPhlF
RiboJ
BCD1
synzip18 - linker - mCherryC - H6
ECK120033736
(13)
RK2
araC
ParaBAD
B0030
srpRN - M86N
ECK120029600
ECK120033737
(14)
p15A
PlacIQ
phl2
phlF
IOT
PPhlF
RiboJ
B0032
M86C - srpRC
ECK120033736
IBMc065
p15A
araC
ParaBAD
B0030
mCherryN - M86N
ECK120029600
ECK120033737
cat
L3S2P55
luxR
B0030
J23107
Plux2
B0032
M86C - mCherryC
ECK120033736
IBMc259
pSC101
PTet*
RiboJ10
B0064
2ndVal-mScarlet-I
L3S2P21
IBMc330
pSC101
PSrpR
B0030
2ndVal-mScarlet-I
L3S2P11
IBMc120
pSC101
Pecf20
B0030
2ndVal-mScarlet-I
B0015
IBMc393
pSC101
ECK120029600
PT7
B0030
2ndVal-mScarlet-I
B1006
T7Φ
PTH

**Supplementary Figure 27 Schematics for constructs used in this study.** Schematics were drawn according to SBOL Visual v2.2.0[26]. See **Supplementary Data 1** to match constructs to numbered schematics and figures. Glyphs for restriction sites are colored in grey. Elements in parenthesis, for instance the hexahistidine tag $H_6$, are unique to some constructs and do not appear in others. Short amino acid insertions (< 10 residues) that are not linkers are omitted in all schematics. Plasmid backbones[27], promoters[4, 28], ribosome binding sites[4], bicistronic designs[9] (BCDs), insulators[29], *mScarlet-I*[30] and terminators[31, 32] were obtained or engineered from various studies or the Registry of Standard Biological Parts (http://parts.igem.org/Main_Page). Other genetic elements have been cited in the main text or their respective supplementary figures. In all cases orthogonal terminators were used for constructs that co-existed within the same cell. The only exception was the TetR reporter plasmid which, due to cloning issues, reused the terminator L3S2P21 once. Plasmid sequences are available at SynBioHub[33] (See Data Availability).

| Target protein | Total reads count * | Aligned reads count | Possible DNA insertion sites count | Library fold coverage † | Missing DNA insertion sites count | DNA insertion coverage | Possible amino acid split / insertion sites count | Missing amino acid split / insertion sites count | Amino acid split / insertion site coverage | Missing amino acid split / insertion sites |
|---|---|---|---|---|---|---|---|---|---|---|
| mCherry | 17,851,597 | 1,012,384 | 1138 | 890 | 125 | 89% | 189 | 11 | 94.2% | 30/31, 36/37, 40/41, 51/52, 78/79, 127/128, 130/131, 166/167, 170/171, 193/194, 216/217 |
| TetR | 18,343,364 | 1,129,105 | 1004 | 1124 | 102 | 89.8% | 168 | 21 | 87.5% | 23/24, 28/29, 32/33, 49/50, 60/61, 63/64, 64/65, 74/75, 76/77, 77/78, 80/81, 91/92, 92/93, 107/108, 110/111, 114/115, 121/122, 123/124, 150/151, 168/169, 177/178 |
| SrpR | 19,336,076 | 1,284,795 | 1010 | 1272 | 140 | 86.1% | 168 | 16 | 90.5% | 34/35, 72/73, 92/93, 96/97, 99/100, 100/101, 108/109, 110/111, 112/113, 131/132, 143/144, 144/145, 150/151, 167/168, 169/170, 174/175 |
| ECF20 | 20,904,174 | 1,380,547 | 1082 | 1276 | 58 | 94.6% | 180 | 13 | 92.8% | 8/9, 31/32, 37/38, 53/54, 88/89, 113/114, 127/128, 133/134, 134/135, 143/144, 146/147, 165/166, 173/174 |
| M86 (domain insertion by ER-LBD) | 13,258,758 | 2,015,735 | 850 | 2371 | 80 | 90.6% | 141 | 13 | 90.8% | 8/9, 26/27, 40/41, 61/62, 71/72, 82/83, 97/98, 108/109, 111/112, 113/114, 133/134, 135/136, 143/144 |

**Supplementary Table 1 Summary of NGS results of the five screenable libraries created in IBM and DIM experiments.** (*) Total reads count includes both the forward and reverse reads from paired-ends. (†) Library fold coverage = Aligned reads count / Possible DNA insertion sites count. See Methods on how aligned reads were called.

### References

1. Manna, D., Deng, S., Breier, A.M. & Higgins, N.P. Bacteriophage Mu Targets the Trinucleotide Sequence CGG. *Journal of Bacteriology* **187**, 3586-3588 (2005).
2. Haapa, S., Taira, S., Heikkinen, E. & Savilahti, H. An efficient and accurate integration of mini-Mu transposons in vitro : a general methodology for functional genetic analysis and molecular biology applications. *Nucleic Acids Research* **27**, 2777-2784 (1999).
3. Nadler, D.C., Morgan, S.-A., Flamholz, A., Kortright, K.E. & Savage, D.F. Rapid construction of metabolite biosensors using domain-insertion profiling. *Nature Communications* **7**, 12266 (2016).
4. Meyer, A.J., Segall-Shapiro, T.H., Glassey, E., Zhang, J. & Voigt, C.A. Escherichia coli "Marionette" strains with 12 highly optimized small-molecule sensors. *Nature Chemical Biology* **15**, 196-204 (2019).
5. Goldhaber-Gordon, I., Williams, T.L. & Baker, T.A. DNA Recognition Sites Activate MuA Transposase to Perform Transposition of Non-Mu DNA. *Journal of Biological Chemistry* **277**, 7694-7702 (2002).
6. Appleby-Tagoe, J.H. et al. Highly Efficient and More General cis- and trans-Splicing Inteins through Sequential Directed Evolution. *Journal of Biological Chemistry* **286**, 34440-34447 (2011).
7. Shu, X., Shaner, N.C., Yarbrough, C.A., Tsien, R.Y. & Remington, S.J. Novel Chromophores and Buried Charges Control Color in mFruits. *Biochemistry* **45**, 9639-9647 (2006).
8. Thompson, K.E., Bashor, C.J., Lim, W.A. & Keating, A.E. SYNZIP Protein Interaction Toolbox: in Vitro and in Vivo Specifications of Heterospecific Coiled-Coil Interaction Domains. *ACS Synthetic Biology* **1**, 118-129 (2012).
9. Mutalik, V.K. et al. Precise and reliable gene expression via standard transcription and translation initiation elements. *Nature Methods* **10**, 354-360 (2013).
10. Galarneau, A., Primeau, M., Trudeau, L.-E. & Michnick, S.W. β-Lactamase protein fragment complementation assays as in vivo and in vitro sensors of protein–protein interactions. *Nature Biotechnology* **20**, 619-622 (2002).
11. Wehrman, T., Kleaveland, B., Her, J.-H., Balint, R.F. & Blau, H.M. Protein–protein interactions monitored in mammalian cells via complementation of β-lactamase enzyme fragments. *Proceedings of the National Academy of Sciences* **99**, 3469-3474 (2002).
12. Palanisamy, N. et al. Split intein-mediated selection of cells containing two plasmids using a single antibiotic. *Nature Communications* **10**, 4967 (2019).
13. Drozdetskiy, A., Cole, C., Procter, J. & Barton, G.J. JPred4: a protein secondary structure prediction server. *Nucleic Acids Research* **43**, W389-W394 (2015).
14. Bienert, S. et al. The SWISS-MODEL Repository—new features and functionality. *Nucleic Acids Research* **45**, D313-D319 (2016).
15. Gu, R. et al. Conformational change of the AcrR regulator reveals a possible mechanism of induction. *Acta Crystallographica Section F* **64**, 584-588 (2008).
16. Fang, C. et al. Structures and mechanism of transcription initiation by bacterial ECF factors. *Nucleic Acids Research* **47**, 7094-7104 (2019).
17. Pei, J., Kim, B.-H. & Grishin, N.V. PROMALS3D: a tool for multiple protein sequence and structure alignments. *Nucleic Acids Research* **36**, 2295-2300 (2008).
18. Peck, Sun H., Chen, I. & Liu, David R. Directed Evolution of a Small-Molecule-Triggered Intein with Improved Splicing Properties in Mammalian Cells. *Chemistry & Biology* **18**, 619-630 (2011).
19. Skretas, G. & Wood, D.W. Regulation of protein activity with small-molecule-controlled inteins. *Protein Science* **14**, 523-532 (2005).
20. Gramespacher, J.A., Burton, A.J., Guerra, L.F. & Muir, T.W. Proximity Induced Splicing Utilizing Caged Split Inteins. *Journal of the American Chemical Society* **141**, 13708-13712 (2019).

21. Oakes, B.L. et al. Profiling of engineering hotspots identifies an allosteric CRISPR-Cas9 switch. *Nature Biotechnology* **34**, 646 (2016).

22. Friedel, K. et al. A functional interplay between intein and extein sequences in protein splicing compensates for the essential block B histidine. *Chemical Science* **10**, 239-251 (2019).

23. Pu, J., Zinkus-Boltz, J. & Dickinson, B.C. Evolution of a split RNA polymerase as a versatile biosensor platform. *Nature Chemical Biology* **13**, 432-438 (2017).

24. Chang, H.-J. et al. A Modular Receptor Platform To Expand the Sensing Repertoire of Bacteria. *ACS Synthetic Biology* **7**, 166-175 (2018).

25. Lesne, J. et al. Structural basis for chemically-induced homodimerization of a single domain antibody. *Scientific Reports* **9**, 1840 (2019).

26. Mısırlı, G. et al. SBOL Visual 2 Ontology. *ACS Synthetic Biology* **9**, 972-977 (2020).

27. Martínez-García, E., Aparicio, T., Goñi-Moreno, A., Fraile, S. & de Lorenzo, V. SEVA 2.0: an update of the Standard European Vector Architecture for de-/re-construction of bacterial functionalities. *Nucleic Acids Research* **43**, D1183-D1189 (2014).

28. Stanton, B.C. et al. Genomic mining of prokaryotic repressors for orthogonal logic gates. *Nature Chemical Biology* **10**, 99-105 (2014).

29. Nielsen, A.A. et al. Genetic circuit design automation. *Science* **352**, aac7341 (2016).

30. Balleza, E., Kim, J.M. & Cluzel, P. Systematic characterization of maturation time of fluorescent proteins in living cells. *Nature Methods* **15**, 47 (2017).

31. Chen, Y.-J. et al. Characterization of 582 natural and synthetic terminators and quantification of their design constraints. *Nature Methods* **10**, 659-664 (2013).

32. Zong, Y. et al. Insulated transcriptional elements enable precise design of genetic circuits. *Nature Communications* **8**, 52 (2017).

33. McLaughlin, J.A. et al. SynBioHub: A Standards-Enabled Design Repository for Synthetic Biology. *ACS Synthetic Biology* **7**, 682-688 (2018).
